# Nuclear Receptor Transcription Factors promote axon regeneration in the Adult Corticospinal Tract

**DOI:** 10.1101/2025.06.19.660319

**Authors:** Yogesh Sahu, Manojkumar Kumaran, Soupayan Banerjee, PS Athul Narayan, Katha Sanyal, Meghana Konda, Shringika Soni, Dhruva Kumar, Anisha S Menon, Prakash Chermakani, Aarthi Sukumar, Faheem Farooq, Sneha Manjunath, Deepta Susan Beji, Arupam Biswas, Ishwariya Venkatesh

## Abstract

Transcription factors are potent levers for neural repair, but which factors govern regenerative capacity in the corticospinal tract remains largely unknown. By intersecting developmental RNA-seq with ATAC-seq footprinting, we identified two nuclear-receptor transcription factors, NR2F1 and NR2F6, neither previously linked to CNS axon growth, whose chromatin occupancy at pro-growth enhancers is progressively lost as neurons mature. Forced expression of either factor significantly increased neurite outgrowth in single-neuron tracing assays, and each drove strong cross-midline sprouting after pyramidotomy and long-tract CST regeneration after complete thoracic crush, with concordant recovery of hip-rise kinematics and grip strength. Parallel multi-omic profiling (CUT&RUN, snRNA-seq and Ribo-seq) of both factors together with NR2F6 Hi-C revealed distinct mechanisms: NR2F1 reactivated chromatin-remodeling and cytoskeletal programs, whereas NR2F6, via a conserved corepressor domain, re-occupied developmental enhancers, reorganized three-dimensional chromatin architecture into new topologically associating domains, and imposed a transient translational down-shift in which growth-relevant modules were selectively preserved through translational buffering. Together, these data identify NR2F nuclear receptors as regulators of corticospinal regeneration, acting through enhancer redeployment, translational reprogramming and three-dimensional genome reorganization.

## Introduction

Central-nervous-system (CNS) neurons progressively lose their intrinsic capacity to regrow injured axons, whereas regeneration-competent populations such as peripheral sensory neurons, zebrafish CNS neurons and embryonic mammalian neurons mount robust transcriptional programs that re-activate developmental regulators (Mahar & Cavalli, 2018; Moore *et al*, 2009; Sun & He, 2010; Venkatesh & Blackmore, 2017; Wang *et al*, 2025; Williams *et al*, 2015). Transcription factors (TFs) sit atop these programs and are therefore strong therapeutic candidates(Moore *et al*, 2009; Williams *et al*, 2015; Wang *et al*, 2018; Bareyre *et al*, 2011; Mehta *et al*, 2016; Seijffers *et al*, 2007; Li *et al*, 2015; Belin *et al*, 2015; Wang *et al*, 2015; Norsworthy *et al*, 2017; Avraham *et al*, 2022; Lindborg *et al*, 2021); indeed, forced expression of individual TFs such as KLF7 or SOX11 can improve axon extension and functional recovery(Moore *et al*, 2009; Wang *et al*, 2018, 2015; Kramer *et al*, 2021; Venkatesh *et al*, 2021; Blackmore *et al*, 2012). Nevertheless, regeneration is governed by cooperative TF networks, and combinatorial approaches have generally outperformed single-factor treatments(Venkatesh & Blackmore, 2017; Venkatesh *et al*, 2021; Kramer *et al*, 2021).

In peripheral neurons, hierarchically organized waves of TF families orchestrate injury responses(Li *et al*, 2015; Chandran *et al*, 2016; Avraham *et al*, 2021), but only a handful of TFs have been shown to promote long-tract regeneration in the adult CNS, like KLF6, KLF7 and SOX11. We recently demonstrated that TF co-occupancy of developmental enhancers can be exploited to discover cooperative pairs: KLF6 and its binding partner NR5A2 synergized to enhance corticospinal-tract (CST) sprouting after pyramidotomy(Venkatesh *et al*, 2021). However, major gaps remain. Beyond the KLF family, no lineage of related TFs has been shown to act in concert to drive CNS regeneration, and an unbiased screen for factors that co-occupy pro-growth regulatory DNA across development has not been performed.

Here we integrate developmental RNA-seq with ATAC-seq footprinting to identify the nuclear-receptor family as previously unrecognized regulators of corticospinal repair, pinpointing NR2F1 and NR2F6 as cooperative drivers of axon regrowth. Forced expression of either factor significantly increased neurite outgrowth in single-neuron tracing assays and, *in vivo*, promoted midline sprouting after pyramidotomy and long-distance CST regeneration across a complete thoracic crush, with corresponding gains in hip-rise kinematics and grip strength. Parallel multi-omic dissection of both factors (CUT&RUN, snRNA-seq, Ribo-seq) along with Hi-C of NR2F6 revealed distinct but convergent mechanisms: NR2F1 re-engaged chromatin-remodeling and cytoskeletal networks, whereas NR2F6, via a conserved corepressor motif, re-occupied its developmental enhancer targets, imposed a transient translational down-shift that selectively preserved growth-relevant modules through translational buffering, and reorganized three-dimensional chromatin architecture into newly formed topologically associating domains and switched compartments. These findings implicate nuclear-receptor transcription factors in CNS regeneration and reveal translational reprogramming, enhancer redeployment and three-dimensional genome reorganization as previously unrecognized contributors to neuronal repair.

## Results

### Developmental gene-chromatin filtering nominates NR2F1 and NR2F6 as potential gene candidates to re-activate intrinsic axon-growth programs

To identify transcription factors that control the large axon-elongation capacity of embryonic neurons and could potentially be re-engaged in the adult CNS, we combined age-matched RNA-seq and ATAC-seq datasets retrieved from the open-access ENCODE resource(The ENCODE Project Consortium *et al*, 2020), together with Gene-Ontology (GO) analysis, across mouse cortical development (E11 to P0 to adult).

To minimize noise from generic transcription-factor (TF) networks, we first isolated only those genes that (i) decline ≥2-fold from embryonic day 11 to adulthood (adjusted p-value < 0.05) and (ii) map to Gene-Ontology terms directly linked to neurite outgrowth. This focused list of 194 transcripts was heavily enriched for growth-related biology (Supplementary Table S1). DAVID over-representation analysis (Biological Process domain, EASE p ≤ 0.05) identified 24 significantly enriched GO terms. Mapping these terms onto the Gene Ontology hierarchy shows that they span several related growth-associated lineages, including protein synthesis, nervous-system development and axon guidance (Fig. 1a). The most prominent terms were “cytoplasmic translation” (40 genes), “nervous-system development” (35 genes) and “axon guidance” (19 genes); gene counts indicate the number of the 194 input genes annotated to each term. Full enrichment results are provided in Supplementary Table S2.

**Figure 1.**
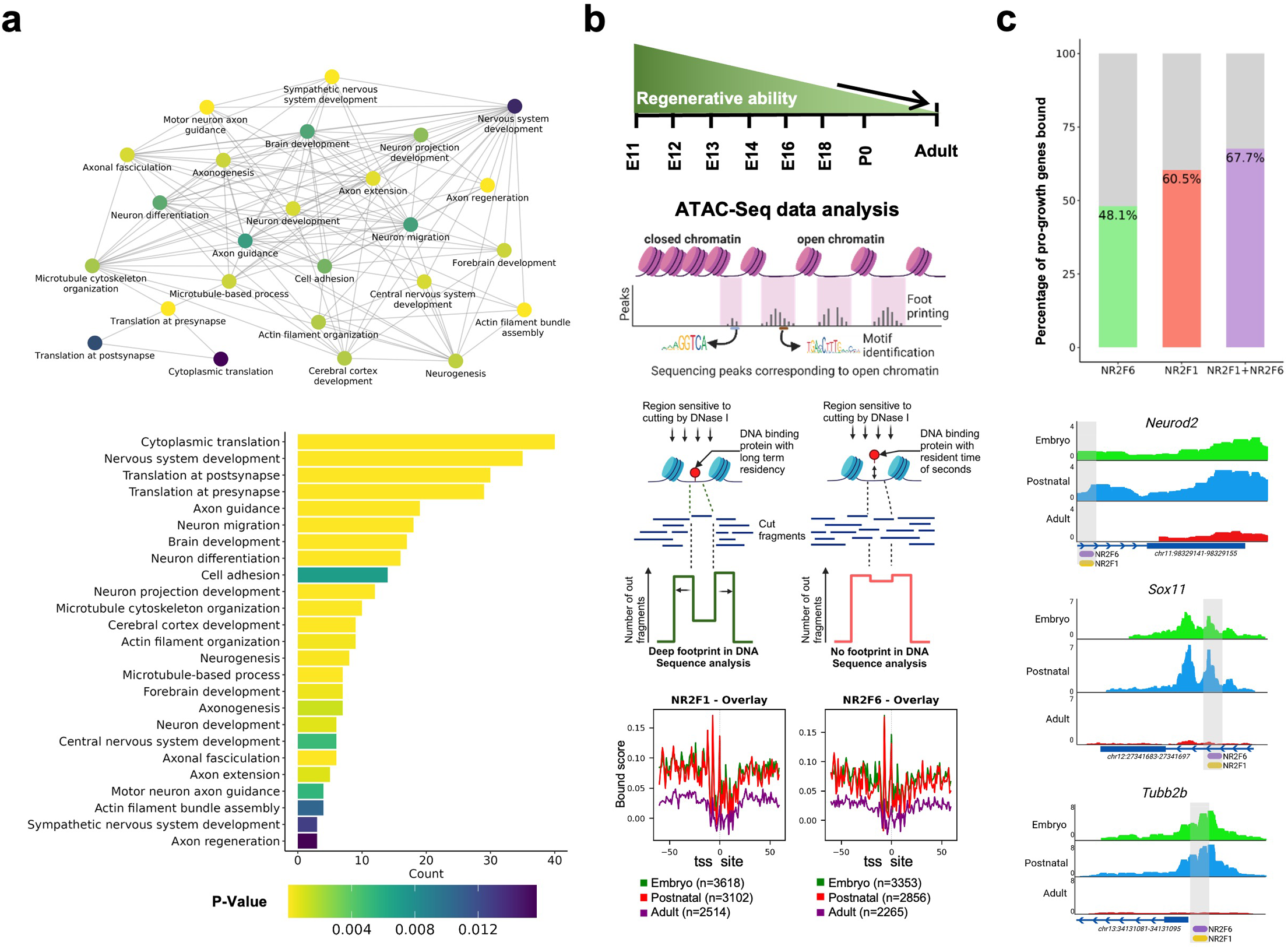
Identification of NR2F1 and NR2F6 as potential pro-growth transcription factors in silico analyses. (a) Gene Ontology (GO) analysis reveals the enrichment of growth-relevant biological processes among genes that are developmentally downregulated in mouse cortical neurons. The network visualization connects related biological processes, while the bar plot ranks the terms by significance (p-value <= 0.05) and gene counts among the 194 candidate genes. (b) Time-series ATAC-seq data from mouse cortical neurons were analysed to investigate changes in chromatin accessibility and transcription factor binding across developmental stages. The upper panel illustrates the decline in regenerative capacity of neurons with age, which correlates with a shift from open to closed chromatin states. The middle schematic demonstrates the ATAC-seq workflow used to identify transcription factor footprints, focusing on motifs associated with chromatin regions that transition from open to closed states. The lower panel highlights developmental changes in motif accessibility for NR2F1 and NR2F6, showing significant footprints during embryonic stages and progressive reduction in postnatal and adult neurons. (c) Quantitative analysis of pro-growth genes targeted by NR2F1 and NR2F6. The bar graph indicates that 60.5% of growth-associated genes are contain NR2F1, 48.1% and NR2F6, 67.7% foot printing. Genome browser snapshots further illustrate the dynamic binding of NR2F1 and NR2F6 at representative growth-associated loci across developmental stages. Strong binding signals are observed in embryonic neurons, with reduced accessibility and binding in postnatal and adult neurons. This integration of gene regulatory and chromatin data highlights the developmental downregulation of NR2F1 and NR2F6 activity and their critical roles in driving early neuronal growth programs.

We performed TF footprinting with the TOBIAS pipeline (BINDetect)(Bentsen *et al*, 2020) on the consensus ATAC-seq peaks of a three-stage cortical series (embryonic, postnatal, adult). To focus on the pro-growth program, footprints were then intersected with the accessible regions of the 194 candidate genes, defined as ATAC-seq peaks overlapping the gene body or lying within ±3 kb of the transcription start site (mm10 annotation). Among the highest-scoring motifs were those for the nuclear-receptor paralogues NR2F1 and NR2F6, whose aggregate binding scores are shown in Fig. 1b. Notably, per-base cleavage profiles displayed deep, well-defined NR2F1/NR2F6 footprints in embryonic nuclei that progressively flattened postnatally, consistent with a loss of stable binding as regenerative capacity wanes.

Cross-referencing footprints with the 194 growth-linked genes showed that NR2F1 binds 117 genes (60.5 %), NR2F6 binds 93 (48.1 %), and together they co-occupy 131 genes (67.7 %) (Fig. 1c, Supplementary Table S3). Genome browser tracks confirmed strong embryonic occupancy at hallmark growth drivers (Neurod2, Sox11, Tubb2b), followed by chromatin closure and signal loss in adult neurons. Consistent with this chromatin-level decline, Nr2f1 and Nr2f6 transcript levels in the murine forebrain are highest during the growth-competent period: Nr2f6 peaks at E12 and falls progressively to its lowest level in the adult (a reduction of approximately 78% from its developmental maximum), while Nr2f1 remains elevated throughout embryonic and perinatal stages and is then markedly reduced in the adult (approximately 69% reduction from peak; Supplementary Fig. S1). For both factors, adult expression is therefore a fraction of the level present during the window of high intrinsic axon-growth capacity. Taken together, this gene-chromatin approach suggests that NR2F1 and NR2F6 are potential candidates for reactivating the embryonic growth program in mature cortical neurons.

### Forced expression of NR2F1 or NR2F6 augments neurite outgrowth in cultured neurons

To test whether the *in silico* nominated candidates can directly stimulate neurite outgrowth, we used over-expressed NR2F1 and NR2F6 in two complementary culture systems using AAV-vectors. Structural integrity of AAV vector was confirmed using SDS-PAGE, data provided in Supplementary Fig. S2. In the Neuro-2A screen (Fig. 2a–c), cells were transduced with AAV-t tdTomato (control), AAV-NR2F6, AAV-NR2F1 or both vectors, switched to differentiation medium after 48 h and fixed a further 48 h later. Across all AAV-transduced neurons with tdTomato signal, mean neurite length rose from 68.7 µm in controls to 111.39 µm with NR2F6 and 98.13 µm with NR2F1; co-delivery of the two factors yielded the strongest effect at 113.35 µm. Statistical analysis was performed using (Kruskal–Wallis test, p < 0.0001), Post-hoc analysis using Dunn’s multiple comparisons test demonstrated that all treatment groups showed a significant increase in neurite length compared to the control (AAV-tdTomato) group (p < 0.0001) for all comparisons (Supplementary Table S4).

**Figure 2.**
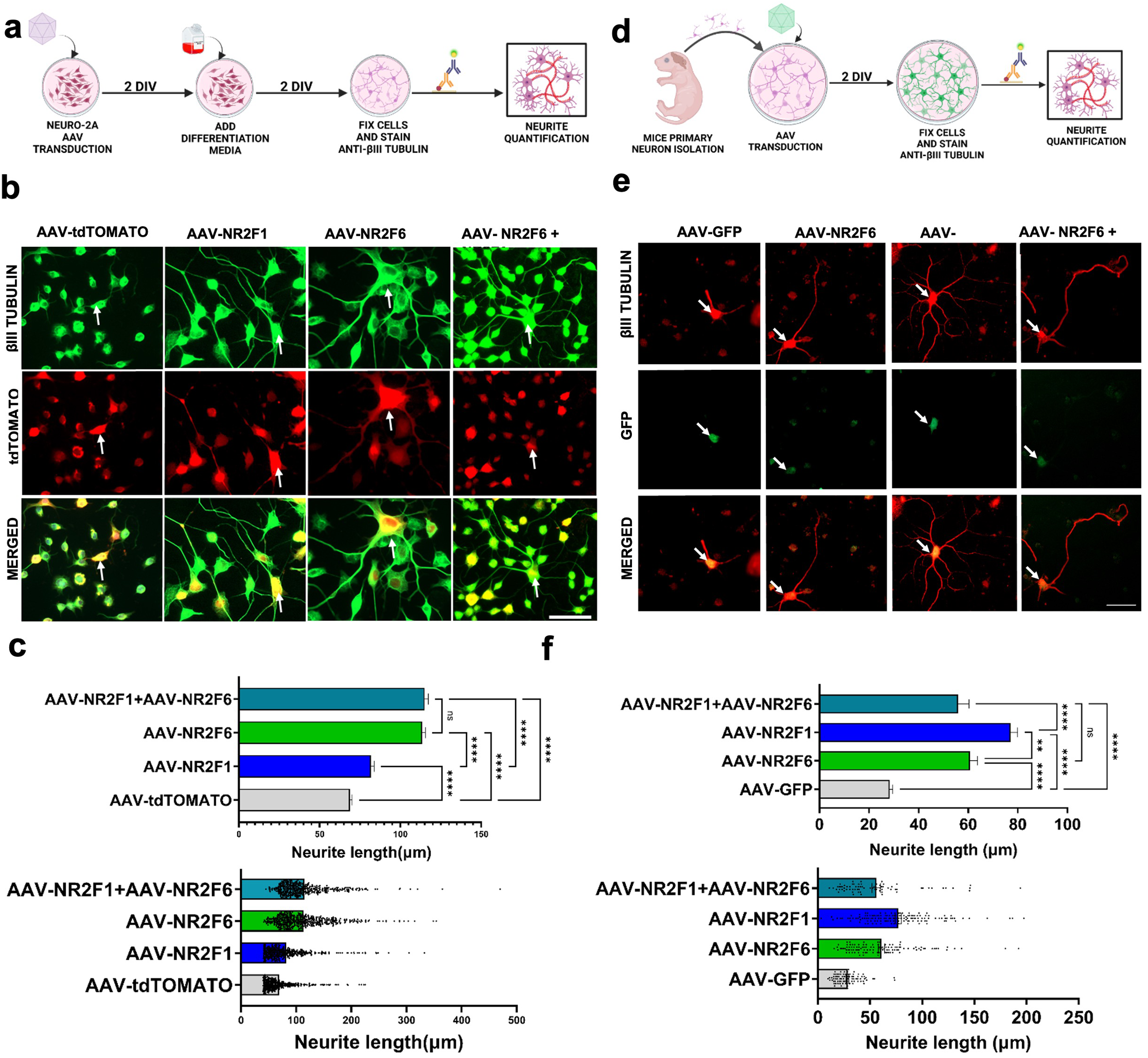
In vitro screening identifies NR2F6 and NR2F1 as potentially significant pro-growth transcription factors. (a) Schematic of the experimental timeline for Neuro-2A in vitro AAV transduction and neurite quantification. Neuro-2A cells were transduced with AAV vectors expressing tdTOMATO as a marker along with NR2F1, NR2F6, or a combination of the two. Following differentiation for 2 DIV, cells were fixed and stained for βIII Tubulin to quantify neurite outgrowth. (b) Representative immunofluorescence images of Neuro-2A cells transduced with AAV-tdTOMATO (control), AAV-NR2F1, AAV-NR2F6, or a combination of AAV-NR2F1 + AAV-NR2F6. Cells were stained for βIII Tubulin(green), and tdTOMATO signal is shown in red. Arrowheads highlight representative cells. (c) Quantification of neurite length in transduced Neuro-2A cells. Statistical analysis was performed using (Kruskal–Wallis test, p < 0.0001), indicating that at least one group differed significantly from the others. Post-hoc analysis using Dunn’s multiple comparisons test demonstrated that all treatment groups showed a significant increase in neurite length compared to the control (AAV-tdTOMATO) group (p < 0.0001 for all comparisons), data represent mean ± SEM. The number of neurons analysed per group was as follows: control (n = 474), NR2F6 (n = 544), NR2F1 (n = 489), and NR2F1+NR2F6 (n = 441) (Scale bar = 50 µm) (d) Schematic of the experimental timeline for mouse primary neuron culture, cells were fixed and stained for βIII Tubulin to quantify the extent of neurite outgrowth. (e) Representative image of neurons transfected with AAV-GFP (control), AAV-NR2F6, AAV-NR2F1, or a combination of AAV-NR2F1 and AAV-NR2F6, and neurite length was quantified. Arrowheads highlight representative cells. (f) Statistical analysis was performed using the Kruskal–Wallis test followed by Dunn’s multiple comparisons test. A significant overall effect of treatment was observed (Kruskal–Wallis test, p < 0.0001). Post-hoc analysis revealed that all treatment groups showed a significant increase in neurite length compared to control (p < 0.0001 for all comparisons). Each data point represents an individual neuron. Data are presented as mean ± SEM. The number of neurons analysed per group was as follows: control (n = 90), NR2F6 (n = 107), NR2F1 (n = 139), and NR2F1+NR2F6 (n = 70)(Scale bar = 50 µm)

We next validated these findings in a more physiologically relevant assay using postnatal day 0/1 mouse primary cortical neurons (Fig. 2d-f). Neurons were plated at low density to enable spatial isolation of individual cells, transduced immediately after plating with AAV-GFP (control), AAV-NR2F6, AAV-NR2F1 or a combination of AAV-NR2F1 and AAV-NR2F6, and analyzed at two days *in vitro*. For each GFP-positive neuron, all processes were traced individually to their terminal tips and summed to give total neurite length per cell. Across three independent culture experiments, all three treatment groups showed significantly greater neurite length compared with GFP control (Kruskal-Wallis p < 0.0001; Dunn’s post-hoc p < 0.0001 for each; GFP ( n = 90), NR2F6 (n=107), NR2F1 (n=139), NR2F1+NR2F6 (n= 70). NR2F1 alone produced significantly longer neurites than NR2F6 (p = 0.0011), confirming the stronger single-factor effect observed in the primary neuron (Supplementary Table S4). In addition, high-density culture data with βIII-tubulin intensity as readout are provided in Supplementary Fig. S3.

These data show that NR2F1 and NR2F6 overexpression cell-autonomously enhanced neurite extension *in vitro*, with NR2F1 exerting the stronger single-factor effect and the combination yielded a robust pro-growth response.

### NR2F1 and NR2F6 enhance corticospinal-tract (CST) sprouting across the midline after unilateral pyramidotomy

Having established that NR2F1 and NR2F6 accelerate neurite extension in vitro, we next asked whether the same factors could promote compensatory growth of mature CST axons *in vivo*. Adult mice received a unilateral cortical injection of AAVs encoding NR2F1, NR2F6, both factors, or GFP alone (Fig. 3a) specifically into layer 5 of the motor cortex, data provided in Supplementary Fig. S4. Seven days later, the ipsilateral CST was transected at the medullary pyramid; lesion completeness was verified by the loss of PKCγ immunoreactivity within the severed tract (Fig. 3b). Eight weeks after injury, cervical spinal cords were examined for GFP-labelled sprouts that crossed from the intact side into the denervated hemicord.

**Figure 3.**
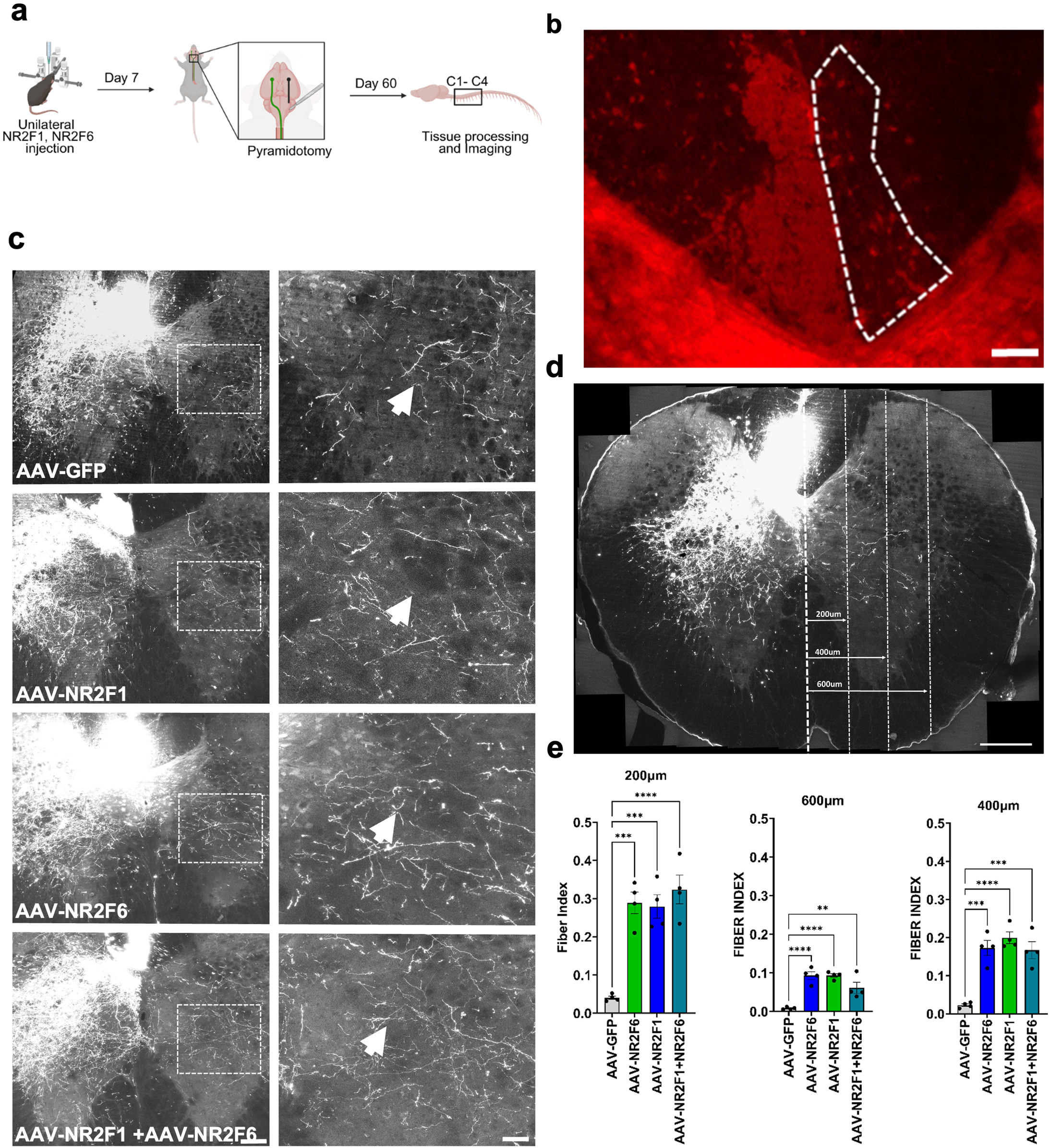
NR2F1 and NR2F6 significantly promote CST sprouting in pyramidotomy model. (a)Schematics representing the timeline and methodology of unilateral pyramidotomy experiment. (b) Unilateral ablation of CST axon (dotted box) was confirmed by PKCγ immunohistochemistry (Scale bar = 100 µm). (c) Representative image of cervical spinal cord in coronal section (Scale bar = 200 µm) and zoomed (25 µm) showing enhanced sprouting across midline in animals treated with NR2F1, NR2F6 and combination of both along with GFP as tracer. (d-e) CST crossing midline was quantified from 200 um, 400 um, and 600 um to midline. Data are shown as mean ± SEM. Groups were compared by one-way ANOVA followed by Dunnett’s post-hoc test against the AAV-GFP control. Variance homogeneity was confirmed by Brown-Forsythe test (all distances p > 0.05). At 200 µm: F(3,12) = 21.18, p < 0.0001. At 400 µm: F(3,12) = 22.47, p < 0.0001. At 600 µm: F(3,12) = 18.79, p < 0.0001. All three treatment groups showed significantly increased fiber index compared to AAV-GFP at every distance measured. **p < 0.01, ***p < 0.001, ****p < 0.0001 vs. AAV-GFP.

All three gain-of-function groups displayed dense GFP-positive fiber extending beyond the midline, whereas controls showed only sparse midline approach (Fig. 3c); high-magnification images of the crossing zone are provided in Supplementary Fig. S5. Sprouting was quantified by a fiber index that normalizes the number of GFP-positive axons crossing parasagittal planes at 200, 400 and 600 µm from the midline to the total number of labelled CST fibers in the medullary pyramids of the same animal, thereby controlling for transduction efficiency (Fig. 3d). One-way ANOVA confirmed a significant treatment effect at every distance (200 µm: F(3,12) = 21.18, p < 0.0001; 400 µm: F(3,12) = 22.47, p < 0.0001; 600 µm: F(3,12) = 18.79, p < 0.0001; variance homogeneity confirmed by Brown-Forsythe test, all p > 0.05). Dunnett’s post-hoc test showed that all three treatment groups had significantly elevated fiber indices compared with the AAV-GFP control at every distance measured (Fig. 3e, Supplementary Table S5).

These results demonstrate that NR2F1 and NR2F6, delivered pre-injury, promote CST collateral sprouting after pyramidotomy, with fibers extending hundreds of micrometers into denervated territory. The fiber index quantifies axons crossing defined planes and therefore reports midline crossing directly, not simply elongation on a permissive substrate. This is consistent with our finding that NR2F6 significantly represses RhoA (snRNA-seq log2FC −0.83, p-adj 4.7 × 10⁻¹⁶), the convergent intracellular node through which scar-derived and myelin-associated inhibitors collapse the growth cone, suggesting that the observed crossing reflects a lowered intrinsic sensitivity to midline inhibitory cues in addition to enhanced growth capacity.

### NR2F1 and NR2F6 promote corticospinal-tract regeneration beyond a severe thoracic crush

To test whether the pro-growth effects of NR2F1 and NR2F6 extend to a complete spinal-cord injury, adult mice received bilateral cortical injections of AAV-GFP (control; n = 9), AAV-NR2F1 (n = 9) or AAV-NR2F6 (n = 8) and, seven days later, underwent a force-calibrated T8 crush (Fig. 4a). Lesion completeness was confirmed by the continuous ring of GFAP-positive astroglia encircling the injury core in every animal (Fig. 4b, Supplementary Fig. S6). Twelve weeks after injury, while larger segments T6-T10 were processed to see properly GFP fluorescence corticospinal-tract (CST) axons before and after site of injury. Expression of the AAV constructs was validated by western blotting in transduced Neuro-2A cells (Supplementary Fig. S7)

**Figure 4.**
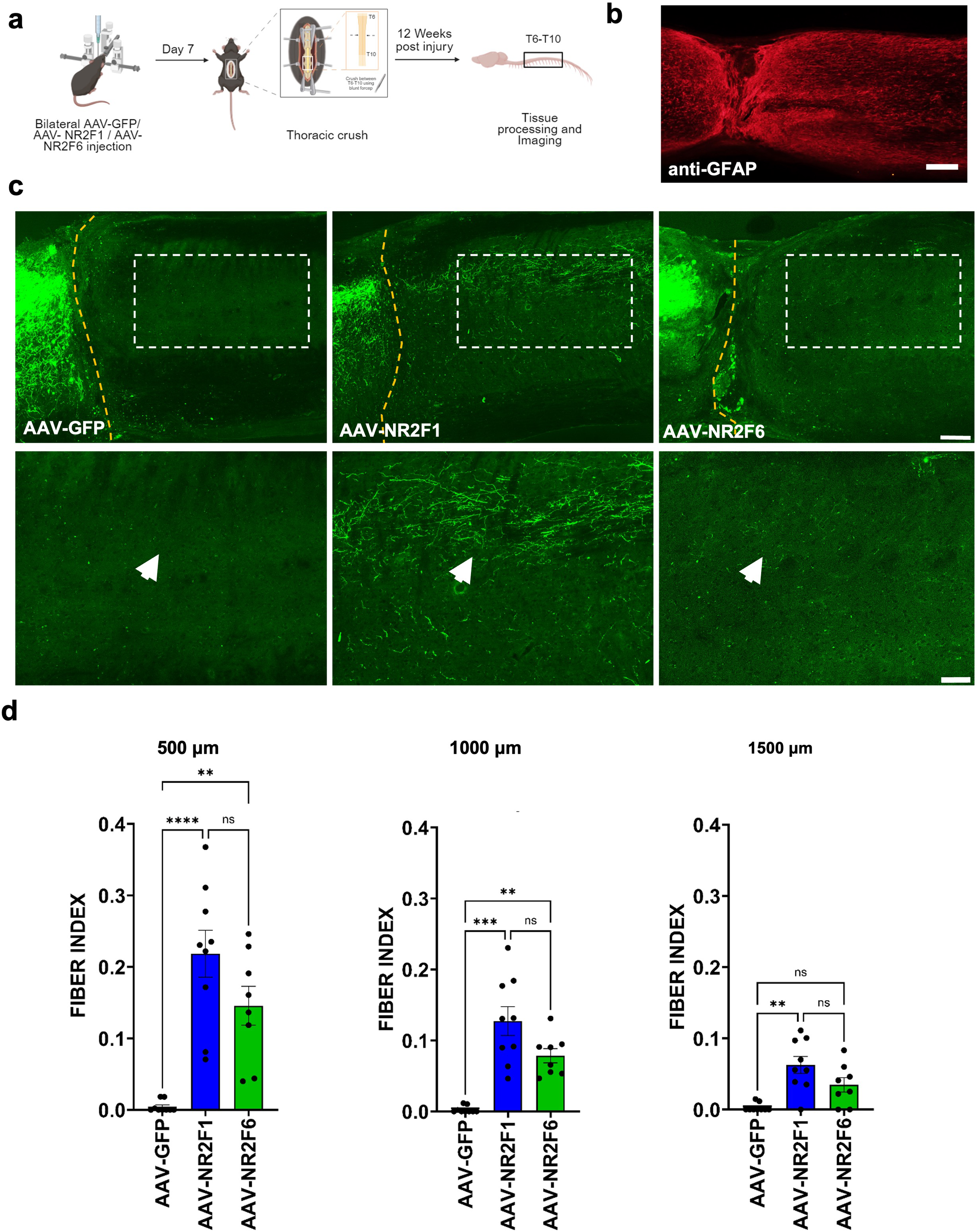
NR2F1 and NR2F6 promote axon regeneration following thoracic spinal cord injury. (a) Schematic of the experimental timeline: bilateral cortical injections of AAV-GFP (control), AAV-NR2F1, or AAV-NR2F6 were performed on day 0, followed by thoracic crush injury at the T8 spinal level on day 7. Animals were sacrificed at 12 weeks post-injury for tissue processing and imaging.(b) Representative sagittal section of the thoracic spinal cord stained for GFAP protein(scale bar =50 µm), showing the injury site at the epicenter of the crush.(c) Sagittal spinal cord sections showing CST axon regeneration distal to the lesion site in control (GFP), NR2F1-, and NR2F6-treated animals. The dashed box indicates the quantified region. Bottom panels show higher magnification of the boxed region, with white arrows indicating regenerating axons (scale bar = 100 µm) and zoomed (scale bar = 75µm). Enhanced axonal regrowth is evident in NR2F1 and NR2F6 groups compared to control.(d) Fiber index was calculated as the number of GFP⁺ axons intersecting parasagittal lines at 500, 1000 and 1500 um caudal to the lesion edge, normalized to total GFP+ axons labelled in the medullary pyramids. Mice received AAV-GFP (control; n = 9), AAV-NR2F1 (n = 9) and AAV-NR2F6 (n = 8). Data are shown as median ± interquartile range. Due to non-normal distribution of the data, groups were compared using the Kruskal-Wallis test followed by Dunn’s post-hoc test with Bonferroni correction. Kruskal-Wallis: 500 µm H = 18.39, p = 0.0001; 1000 µm H = 18.25, p = 0.0001; 1500 µm H = 12.67, p = 0.0018. AAV-NR2F1 showed significantly elevated fiber index versus AAV-GFP at all three distances. AAV-NR2F6 was significant at 500 µm and 1000 µm but not at 1500 µm. No significant difference was observed between AAV-NR2F1 and AAV-NR2F6 at any distance. **p < 0.01, ***p < 0.001, ****p < 0.0001; ns = not significant.

In GFP controls, only a few abortive sprouts approached the glial scar without traversing it (Fig. 4c). In contrast, NR2F1 overexpression produced GFP-positive fibers that crossed the lesion and extended hundreds of micrometers into caudal grey matter; NR2F6 induced a qualitatively similar, though less extensive, response. Regeneration was quantified with a fiber index that divides the number of GFP-labelled CST axons intersecting parasagittal lines at 500, 1000 and 1500 µm distal to the lesion edge by the total number of labelled CST fibers in the medullary pyramids of the same animal, thereby normalizing for transduction efficiency. Because fiber-index distributions were non-normal, groups were compared using the Kruskal-Wallis test followed by Dunn’s post-hoc test with Bonferroni correction. A significant treatment effect was observed at all three distances (500 µm: H = 18.39, p = 0.0001; 1000 µm: H = 18.25, p = 0.0001; 1500 µm: H = 12.67, p = 0.0018). NR2F1 showed significantly elevated fiber indices versus GFP at all three distances; NR2F6 was significant at 500 and 1000 µm but did not reach significance at 1500 µm. No significant difference was detected between NR2F1 and NR2F6 at any distance (Fig. 4d, Supplementary Table S6).

These results show that NR2F1 and NR2F6 promote CST regeneration across and beyond a complete thoracic crush. NR2F1 produced more anatomically extensive regeneration, reaching significance at all distances measured, while NR2F6 regeneration was significant proximally but attenuated at the greatest distance. This anatomical difference is notable in light of the behavioural data presented below, where the two factors produce comparable functional recovery.

### NR2F1 and NR2F6 improve hind-limb kinematics and grip strength after thoracic crush injury

To determine whether the anatomical regeneration driven by NR2F1 or NR2F6 translates into functional benefit, we tracked freely moving mice on a horizontal ladder every seven days from week 1 to week 12 post-injury (Fig. 5a; n = 4 AAV-GFP, 5 AAV-NR2F1, 4 AAV-NR2F6). Locomotor trajectories were captured at 30 fps and analyzed with DeepLabCut, which returned sub-pixel coordinates for six hind-limb landmarks; custom scripts reconstructed stance-swing stick plots (Fig. 5d).

**Figure 5.**
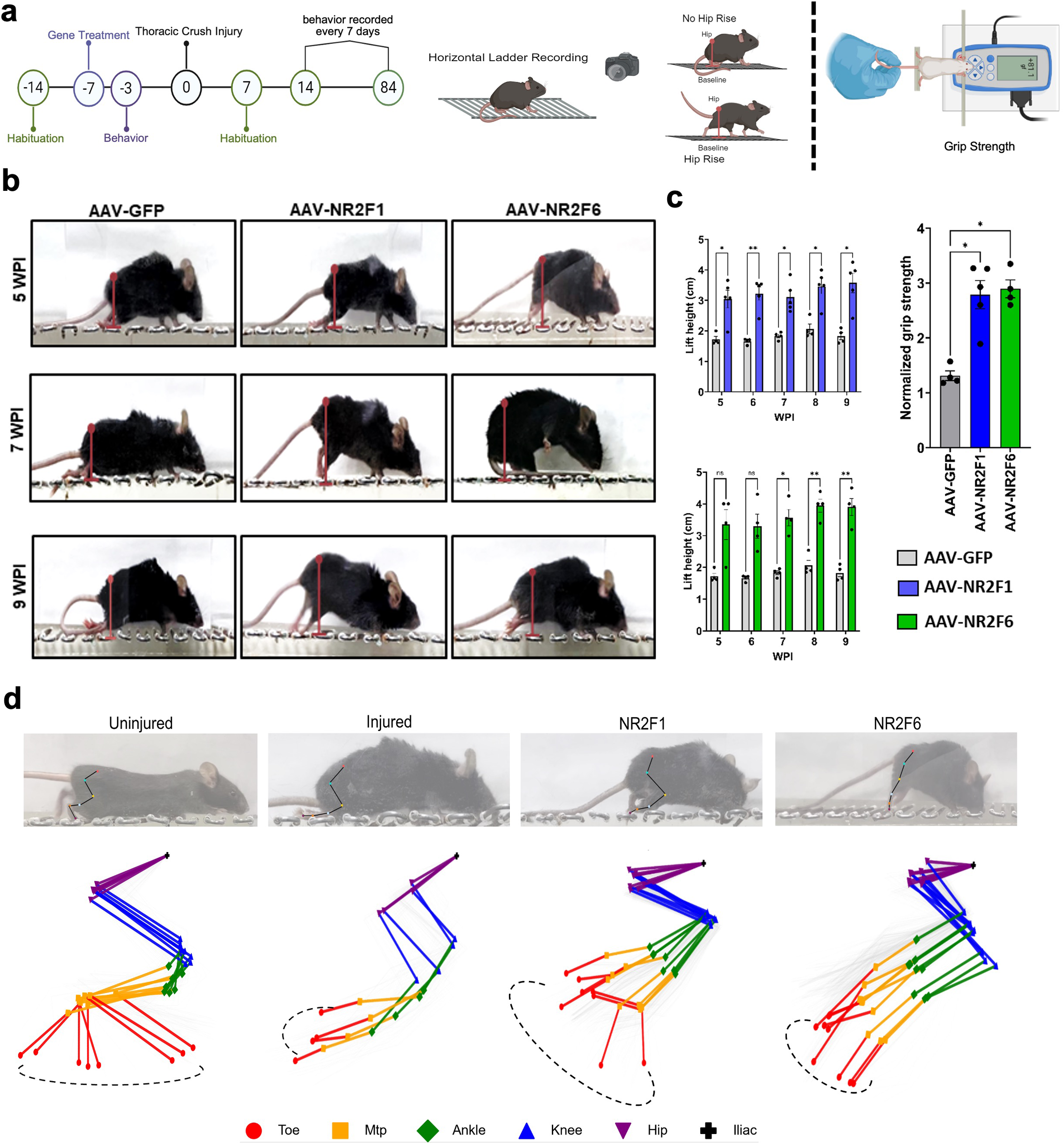
Assessment of motor function recovery following thoracic crush injury and gene treatment. (a) Timeline of the spinal cord injury (SCI) experiment and schematic of the horizontal ladder rung walking test setup, along with the methodology for measuring hip rise. Although animals were recorded weekly from 1 to 12 WPI, statistical comparison of hip rise was performed across the 5-9 WPI recovery window (b) Representative images showing hip rise analysis during horizontal ladder walking at 5, 7, and 9 weeks post-injury (WPI) in AAV-GFP (control), AAV-NR2F1, and AAV-NR2F6 treated animals. (c) Grip strength was compared across AAV-GFP, AAV-NR2F1 and AAV-NR2F6 groups using a Kruskal-Wallis test followed by Dunn’s multiple comparisons test. Overall Kruskal-Wallis p = 0.0113; AAV-GFP vs AAV-NR2F1, p = 0.0257; AAV-GFP vs AAV-NR2F6, p = 0.0365. n = 4 AAV-GFP, n = 5 AAV-NR2F1 and n = 4 AAV-NR2F6 animals. Hip-rise performance was analyzed from 5 to 9 WPI using two-way repeated-measures ANOVA with time as the repeated factor, followed by Bonferroni-Sidak multiple comparisons comparing treatment groups at each time point. For AAV-NR2F1 vs AAV-GFP, treatment effect p = 0.00098; adjusted p values were 5 WPI p = 0.0407, 6 WPI p = 0.0079, 7 WPI p = 0.0154, 8 WPI p = 0.0116 and 9 WPI p = 0.0198. n = 4 AAV-GFP and n = 5 AAV-NR2F1 animals. For AAV-NR2F6 vs AAV-GFP, treatment effect p = 0.00115; adjusted p values were 5 WPI p = 0.1776, 6 WPI p = 0.1070, 7 WPI p = 0.0257, 8 WPI p = 0.0024 and 9 WPI p = 0.0097. n = 4 animals per group.Although animals were recorded weekly from 1 to 12 WPI, statistical comparison of hip rise was performed across the 5-9 WPI recovery window (d) The stick plot illustrates gait patterns for each condition, depicting the stance and swing trajectories of key anatomical landmarks during locomotion. Marker tracking was performed using DeepLabCut (DLC), a deep learning-based pose estimation tool that enables precise and automated extraction of joint positions from video recordings. The markers serve as anatomical landmarks to track limb motion and provide insight into biomechanical changes across experimental conditions. Toe is represented by red circles, Metatarsophalangeal joint (Mtp) by orange squares, Ankle by green diamonds, Knee by blue upward-pointing triangles, Hip by purple downward-pointing triangles, and the I liac Crest by black crosses.

Hip-rise height Fig. 5b, a sensitive surrogate of descending supraspinal control, was significantly elevated in both treatment groups across the recovery period. Two-way repeated-measures ANOVA with time as the repeated factor confirmed a significant treatment effect for both NR2F1 versus GFP (p = 0.00098) and NR2F6 versus GFP (p = 0.00115). Post-hoc Bonferroni-Sidak comparisons showed significant gains emerging by week 5 and persisting through week 9 for both factors (per-timepoint adjusted p-values in Fig. 5c).

Grip strength mirrored the kinematic gains. Kruskal-Wallis testing confirmed a significant overall treatment effect (p = 0.0113); Dunn’s post-hoc comparisons showed that both NR2F1 (p = 0.0257) and NR2F6 (p = 0.0365) were significantly elevated relative to GFP controls (Fig. 5c, Supplementary Table S7).

Detailed stick-plot analysis revealed a graded restoration of the hind-limb stepping envelope. Uninjured mice displayed three discrete toe trajectories spanning the full stride cycle: a posterior backstroke, a vertical raise clearing the rung field, and an anterior forward swing (Fig. 5d, Supplementary Fig. S8). Injured controls collapsed to a single low-amplitude trace indicating loss of lift and forward propulsion. NR2F1- and NR2F6-treated animals re-established the first two components: toe tips executed a normal backstroke followed by a vertical lift, eliminating drag and restoring rung clearance. However, the anterior swing arc remained incomplete, with limbs adopting a pattern in which the paw was elevated and replaced near its origin without translating fully forward (Supplementary Fig. S8). Thus, two-thirds of the normal trajectory (retraction and lift) were recovered, whereas the terminal swing needed for complete stride was not. Quantification of spatiotemporal gait parameters (stride length, swing phase and stance phase) at 1 and 9 weeks post-injury confirmed significant improvements in treated versus injured animals (Supplementary Fig. S9, Supplementary Table S8).

These data show that NR2F1 and NR2F6 convert anatomical CST regeneration into measurable functional gains, with elevated hip rise, improved grip strength and partial restoration of coordinated stepping. Notably, the two factors produced comparable behavioural recovery despite the difference in anatomical regeneration extent reported above, indicating that NR2F6’s regeneration, though less extensive, is functionally productive.

### NR2F1 and NR2F6 reactivate intrinsic growth ability via differential strategies

Single-nucleus RNA-seq was performed one week after thoracic crush on GFP-labelled neurons isolated from motor cortex of AAV-GFP, AAV-NR2F1 and AAV-NR2F6 cohorts (Fig. 6a). GFP-positive nuclei were FACS-sorted prior to library preparation, so the entire dataset consists of virally transduced neurons. In the preprint version, transgene-expressing nuclei were identified using Seurat’s automatic read-assignment pipeline; we found that this approach discarded a substantial number of genuinely transgene-positive nuclei owing to inconsistent calling. We therefore developed a custom annotation script that reliably identifies transgene-expressing nuclei, recovering the previously excluded positives; this script has been deposited in our laboratory GitHub repository and the revised procedure is described in Methods.

**Figure 6.**
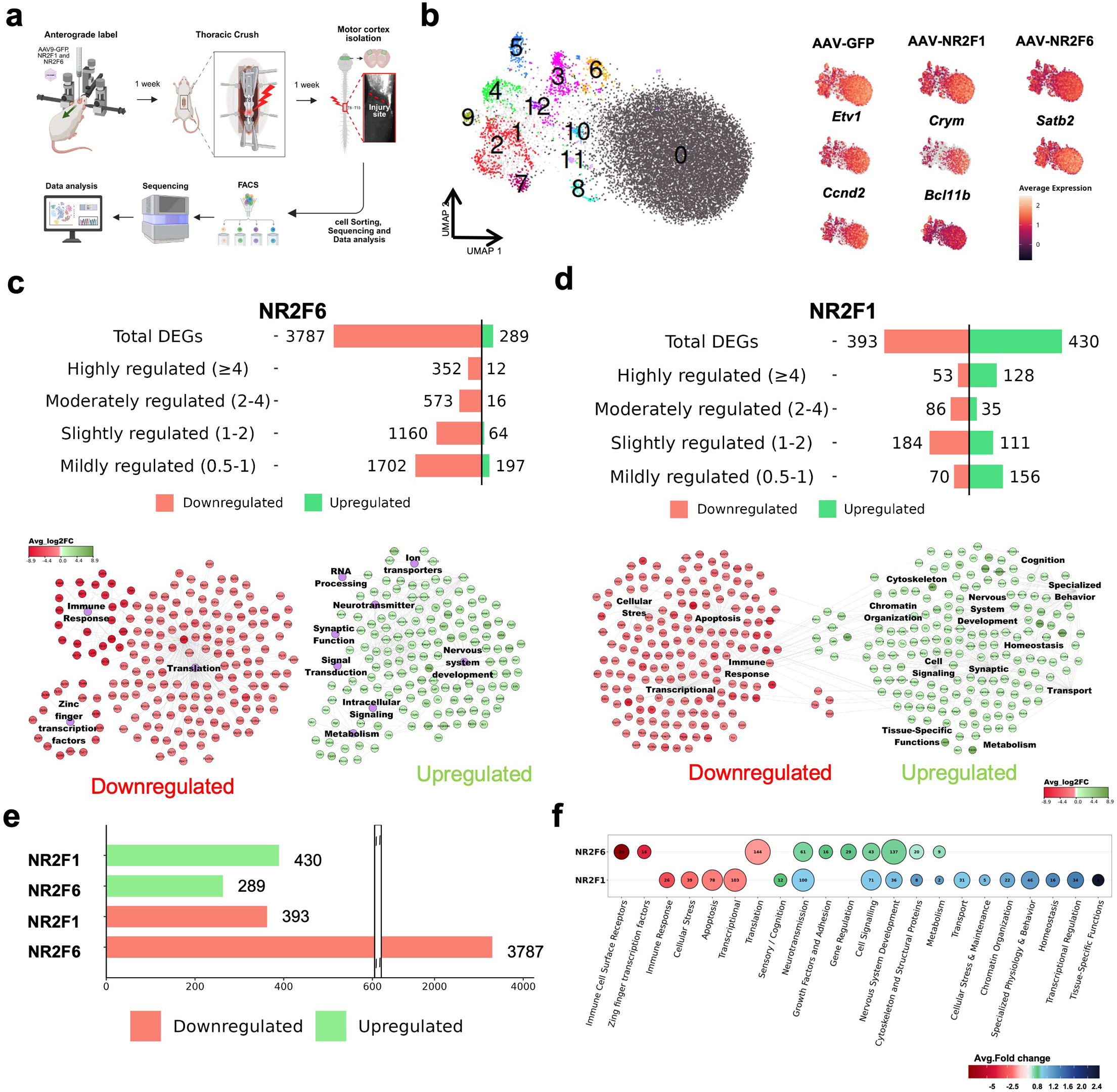
Transcriptional landscapes regulated by NR2F1 and NR2F6 in CST neurons following spinal cord injury, using single-nucleus RNA sequencing (snRNA-seq). (a) Schematic overview of the experimental workflow. Motor cortices were isolated from injured mice injected with AAV-NR2F1 and AAV-NR2F6. Nuclei were extracted and processed for snRNA-seq to assess transcriptional changes in corticospinal tract (CST) neurons. (b) UMAP projection illustrating transcriptional clustering of CST neuronal nuclei from NR2F1-and NR2F6-treated datasets. Distinct clusters are color-coded and labeled, with marker gene expression (Etv1, Crym, Satb2, Ccnd2, Spata16) displayed for selected clusters. (c) Differential gene expression (DEG) analysis of AAV-NR2F6 versus AAV-GFP samples. The bar plot quantifies upregulated (green) and downregulated (red) genes across four fold-change thresholds. The adjacent network diagram highlights enriched biological processes among upregulated genes, including synaptic function, intracellular signaling, and nervous system development. (d) DEG analysis of AAV-NR2F1 versus AAV-GFP samples. The bar plot shows significantly upregulated and downregulated genes, and the network plot illustrates enriched pathways among upregulated genes, such as transcriptional regulation, chromatin organization, and synaptic communication. (e) Comparative bar graph depicting the number of upregulated and downregulated DEGs in NR2F1 and NR2F6 conditions across fold-change categories. (f) Dot plot representing the number of genes involved in key biological pathways regulated by NR2F1 and NR2F6. Dot size corresponds to gene count, and color reflects average log2 fold change.

After quality control we retained 16,328 nuclei (AAV-GFP: 5,403; AAV-NR2F1: 4,748; AAV-NR2F6: 6,177), projected them with UMAP and recovered 13 transcriptional clusters (0-12; Fig. 6b). Cell-type identity was assigned by evaluating canonical marker expression across each cluster (Supplementary Fig. S10, Supplementary Table S9). The major clusters expressed corticospinal/layer-V projection-neuron markers (Satb2, Etv1, Crym, Ccnd2, Bcl11b) at high levels and lacked markers of non-cortical descending populations (paraventricular hypothalamus, red nucleus, locus coeruleus, raphe and others); these were annotated as layer-V extratelencephalic projection neurons of the motor cortex. The small remaining clusters expressed inhibitory-neuron or glial markers and were excluded from differential analysis. Neither NR2F1 nor NR2F6 measurably shifted cluster membership, consistent with transcriptional modulation without cell-identity conversion. Transgene expression was confirmed directly: in each condition the cognate transgene was the single most enriched transcript in the GFP-sorted nuclei (Nr2f6 log2FC 10.6 in the NR2F6 condition; Nr2f1 log2FC 10.5 in the NR2F1 condition; Supplementary Fig. S7).

NR2F6 produced a predominantly repressive transcriptional signature. Relative to GFP it altered 4,076 genes, of which 3,787 (93%) were down-regulated and only 289 were induced (p-adj < 0.05, | log2FC| > 0.5; Fig. 6c, Supplementary Table S10). The down-regulated pool encompassed translational machinery, metabolic enzymes and broad housekeeping programs, consistent with a metabolic reprogramming that pares back general biosynthesis. The up-regulated subset was enriched for synaptic function, signal transduction and nervous-system development (Fig. 6c), indicating that NR2F6 preserves specific circuit-remodeling programs while globally reducing biosynthetic demand.

NR2F1 operated differently. I t modulated 823 transcripts, with a slight bias toward activation (430 up, 393 down; Fig. 6d, Supplementary Table S11). Up-regulated genes mapped to chromatin organization, transcriptional regulation and cytoskeletal remodeling, whereas many metabolic and translational genes were concomitantly reduced, consistent with a targeted reactivation of structural and regulatory programs rather than broad repression.

To verify that these gene sets are not inflated by the cell-level statistical framework, we repeated the differential-expression analysis using the MAST framework with pseudobulk aggregation, which accounts for the correlation structure among nuclei from a pooled sample. The two approaches yielded concordant results with minimal differences in the reported gene sets (Supplementary Table S12), confirming that the transcriptional signatures described here are robust to the choice of DE method.

Direct comparison (Fig. 6e-f, Supplementary Table S13) underscores the complementarity of the two factors: NR2F6 enforces a large-scale transcriptional down-shift while preserving a focused pro-synaptic program, whereas NR2F1 up-regulates chromatin-remodeling and structural modules with more limited global suppression. These different strategies are consistent with the behavioural equivalence of the two factors despite their different anatomical potency: NR2F6 may set the metabolic conditions permissive for growth, while NR2F1 provides the chromatin and cytoskeletal activation that drives axon extension.

### The growth-promoting effect of NR2F6 requires an evolutionarily conserved corepressor domain

Sequence alignment of the NR2F protein family revealed that the well-characterized corepressor motif of NR2F1 and NR2F2 (LSGYISLLL) has a structurally analogous counterpart in NR2F6 (LGIDNVCEL, residues 184-192), located within the ligand-binding domain (Fig. 7a). A corepressor domain has been described for NR2F1 and NR2F2, but not previously for NR2F6, which is structurally and functionally divergent from the other two family members. We therefore treated the NR2F6 motif as a predicted corepressor domain and tested it functionally. We generated an NR2F6 construct lacking this 9-amino-acid motif (NR2F6-DD). Overlap PCR followed by Sanger sequencing confirmed precise excision, leaving the upstream DNA-binding domain and downstream ligand-binding domain in frame (Fig. 7b). Western blotting showed that both wild-type (WT) and DD proteins migrated at the expected size (∼37 kDa), ruling out degradation or truncation (Fig. 7e).

**Figure 7.**
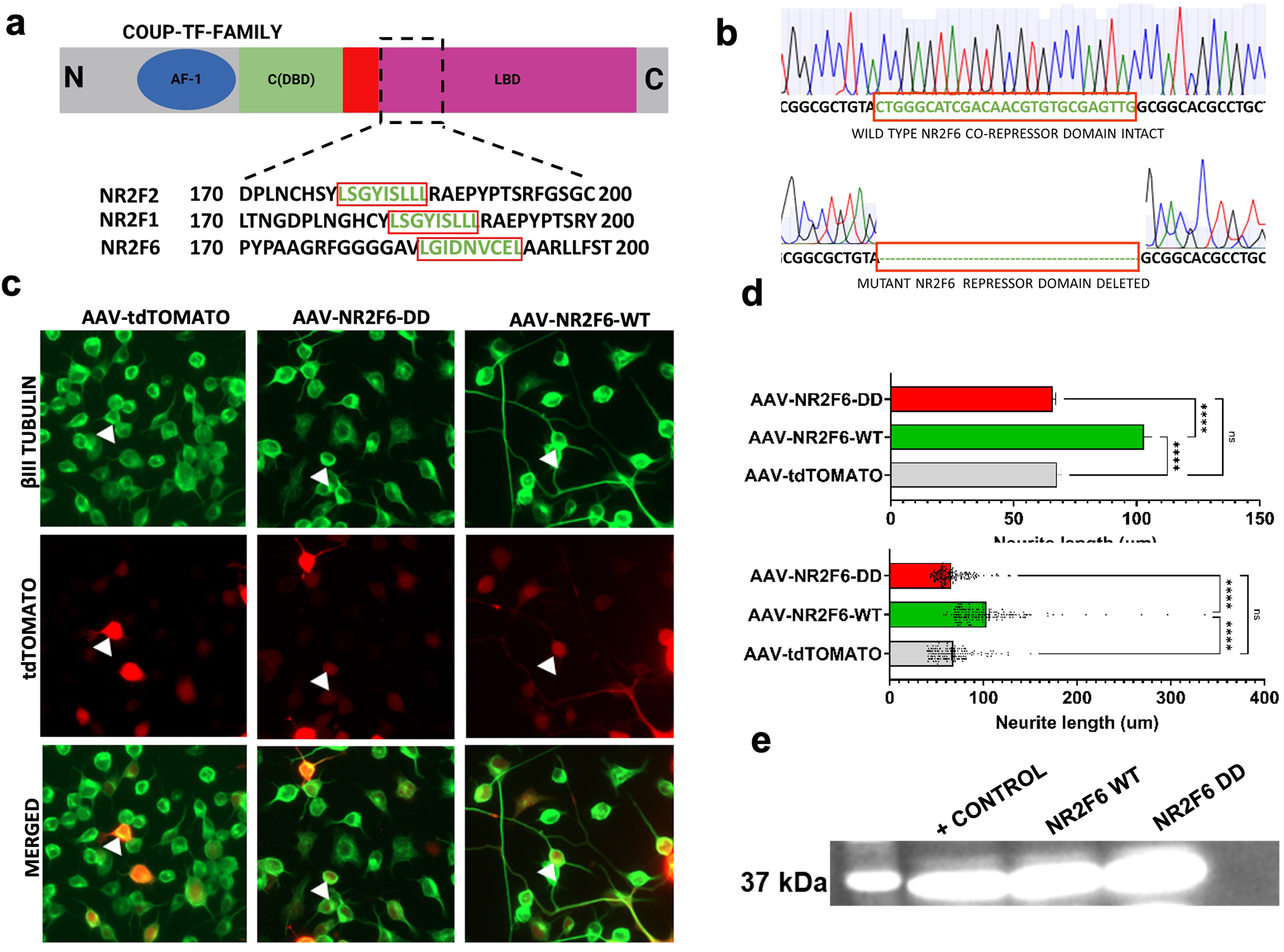
The NR2F6 Corepressor Domain is Essential for Neurite Growth Induction and Regulation. (a) Sequence alignment of the NR2F protein family (NR2F1, NR2F2, and NR2F6) reveals potential conservation of a corepressor domain. (b) Generation and validation of the NR2F6 corepressor domain deletion mutant(NR2F6-DD). Overlapping PCR was used to delete the corepressor domain from NR2F6. Sanger sequencing confirms the successful deletion, as indicated by the absence of the repressor domain sequence in the mutant (bottom) compared to the wild-type NR2F6 (top). (c) Functional evaluation of the NR2F6 corepressor domain in neurite growth using AAV constructs. N2A cells were transduced with AAV constructs expressing tdTOMATO (control), NR2F6-DD (NR2F6 corepressor domain mutant) and NR2F6-WT(wild-type NR2F6). Immunostaining for βIII-tubulin (green) reveals neurite outgrowth, while tdTOMATO fluorescence (red) identifies transduced cells. AAV-NR2F6-WT promotes significant neurite extension (right), while deletion of the corepressor domain (AAV-NR2F6-MUTANT, middle) abolishes this effect, similar to the control (AAV-tdTOMATO, left). Arrowheads highlight representative cells. Scale bar = 50 µm (d) Neurite length was quantified per neurite counted from Neuro-2A transfected with AAV-tdTOMATO (control; n = 135 neurites), AAV-NR2F6-WT (n = 158 neurites), or AAV-NR2F6-DD (domain deleted; n = 156 neurites). Data are shown as mean ± SEM. Groups were compared using the Kruskal-Wallis test followed by Dunn’s post-hoc test with Bonferroni correction (Kruskal-Wallis H = 163.1, p < 0.0001, total n = 449 neurites).AAV-NR2F6-WT showed significantly greater neurite values compared to both AAV-tdTOMATO (p < 0.0001) and AAV-NR2F6-DD (p < 0.0001). AAV-NR2F6-DD was not significantly different from AAV-tdTOMATO (p > 0.9999), indicating that the deleted domain is required for NR2F6-mediated activity. ****p < 0.0001; ns, not significant. (e) western blot was performed to check the formation of full length protein by NR2F6 domain deleted clone. Equal amount of protein 25ug was loaded in all three wells.

Neuro-2A cells were transduced with AAV- tdTomato (control; n = 135 neurites), AAV-NR2F6-WT (n = 158) or AAV-NR2F6-DD (n = 156) and scored 48 h after differentiation. βIII-tubulin immunostaining showed elongated neurites in NR2F6-WT cells, whereas DD-expressing cells resembled controls (Fig. 7c). Kruskal-Wallis testing confirmed a significant treatment effect (H = 163.1, p < 0.0001, total n = 449); Dunn’s post-hoc with Bonferroni correction showed that NR2F6-WT neurite length was significantly greater than both tdTomato and NR2F6-DD (p < 0.0001 for each), while NR2F6-DD did not differ from the control (p > 0.99; Fig. 7d, Supplementary Table S14).

These data demonstrate that the predicted corepressor motif is required for NR2F6-mediated neurite extension, providing the first functional evidence for a corepressor domain in NR2F6 and indicating that the factor operates primarily through transcriptional repression rather than direct activation of growth genes.

### NR2F6 and NR2F1 reshape the translational landscape of injured neurons through distinct mechanisms

To determine how the transcriptional changes imposed by NR2F6 and NR2F1 propagate to the translational level, we performed ribosome profiling (Ribo-seq) on GFP-labelled CST neurons isolated 7 days after thoracic crush from mice that had received AAV-NR2F6 or AAV-NR2F1 seven days before injury (Fig. 8a, Supplementary Fig. S11).

**Figure 8.**
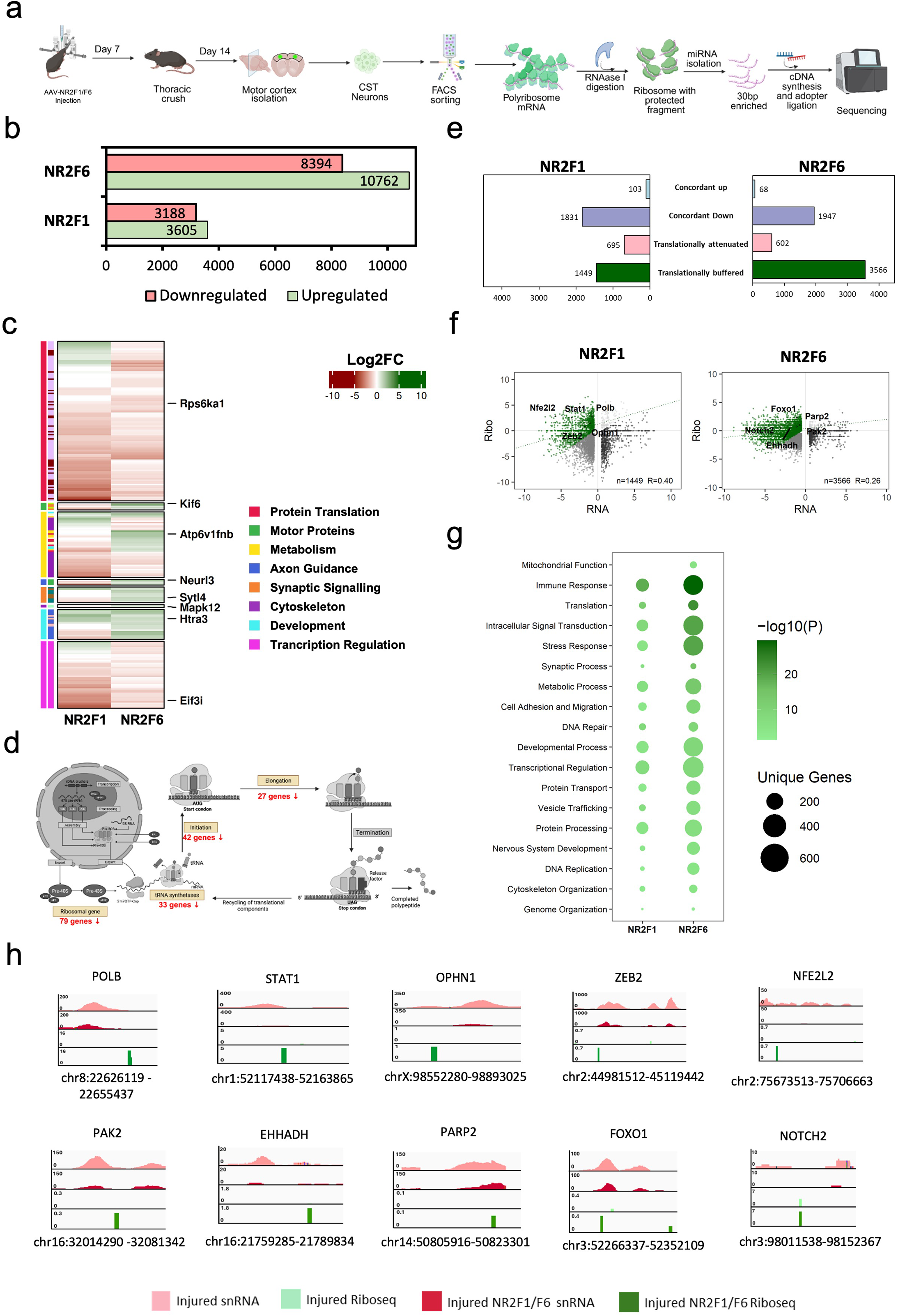
Ribosome profiling (Ribo-seq) reveals global translational alterations driven by NR2F6 and NR2F1 in CST neurons. (a) Schematic overview of the Ribosome profiling (Ribo-seq) experimental pipeline. The timeline illustrates AAV injection, subsequent thoracic crush injury at Day 14, motor cortex isolation, and FACS sorting of CST neurons. This is followed by the isolation of polyribosome mRNA, RNase I digestion to isolate ribosome-protected fragments, and sequencing library preparation. (b) Bar graph quantifying the total number of differentially translated transcripts. Green bars indicate upregulated genes, and salmon red bars indicate downregulated genes for both the NR2F6 and NR2F1. (c) Heatmap visualizing the log2 fold change (Log2FC) of selectively enriched genes across targeted functional categories for both NR2F1 and NR2F6 conditions. Upregulated transcripts are depicted in green, while downregulated transcripts are shown in red. Color-coded sidebars categorize the transcripts into specific functional roles: Protein Translation, Motor Protein, Metabolism, Axon Guidance, Synaptic Signaling, Cytoskeleton, Development, and Transcription Regulation. Representative key genes (e.g., Rps6ka1, Klf6, Atp6v1fnb, Neurl3, Sytl4, Mapk12, Htra3, Eif3i) are explicitly annotated on the right.(d) Schematic representation mapping the specific impact of downregulated transcripts onto the cellular translational machinery. The diagram highlights vulnerabilities across the translation cycle, annotating the precise number of suppressed transcripts associated with ribosomal genes (79 genes), tRNA synthetases (33 genes), translation initiation (42 genes), and translation elongation (27 genes).(e) Bidirectional bar plots showing the number of genes in each regulatory category — concordant up, concordant down, translationally attenuated, and translationally buffered for NR2F1 and NR2F6. NR2F6 exhibits a markedly larger translationally buffered gene set (3,566) compared to NR2F1 (1,449).(f) Scatter plots of snRNA-seq (RNA) versus ribosome occupancy (Ribo) log₂ fold changes for NR2F1 (n=1,449, R=0.40) and NR2F6 (n=3,566, R=0.26), colored by category. The green fitted line for translationally buffered genes deviates from the concordance diagonal toward the Ribo axis, indicating sustained translation despite reduced RNA. The lower R in NR2F6 reflects greater transcription-translation discordance.(g) GO enrichment bubble plot for translationally buffered genes, with bubble size indicating unique gene count and color intensity indicating significance (−log₁₀ P). Both factors enrich for immune response, stress response, and translation-related terms, with NR2F6 showing broader and more significant enrichment. This suggests that translational buffering sustains output of stress-response and immune-regulatory proteins despite transcriptional downregulation, potentially serving as a post-transcriptional resilience mechanism following injury.(h) Genomic screen shots for confirming that buffered genes maintain positive TE despite reduced RNA. (Right) Representative IGV tracks for buffered genes (POLB, STAT1, OPHN1, ZEB2, NFE2L2 in NR2F1; PAK2, FOXO1, NOTCH2, EHHADH, PARP2 in NR2F6), showing maintained Ribo-seq signal alongside reduced snRNA signal at each locus.

NR2F6 altered the ribosome occupancy of 19,156 transcripts relative to injured controls (8,394 decreased, 10,762 increased; Fig. 8b, Supplementary Table S15). NR2F1 altered 6,793 (3,188 decreased, 3,605 increased, Supplementary Table S16). The genome-wide Ribo-seq landscape thus reflects extensive translational reprogramming by both factors rather than a uniform directional shift.

Within this broader reprogramming, NR2F6 imposed a coordinated repression of the protein-synthesis machinery itself. Virtually every tier of the translational apparatus was attenuated: 79 ribosomal-protein genes, 33 tRNA-synthetase genes, 42 initiation-complex factors and 27 elongation-factor transcripts were simultaneously reduced (Fig. 8d). This selective repression of the biosynthetic core contrasts with the genome-wide pattern and indicates that NR2F6 specifically dials back the translational capacity of the cell while permitting, and in many cases increasing, ribosome loading at other loci. The functional-category heatmap (Fig. 8c) shows that this selectivity extends across both factors: NR2F6 represses translation, motor-protein and metabolic transcripts while preserving axon-guidance, synaptic- signaling and developmental modules; NR2F1 shows a qualitatively similar but less pronounced pattern, consistent with its more balanced transcriptional profile.

To ask whether the translational changes are merely a read-out of the transcriptional changes or reflect an independent regulatory layer, we integrated the snRNA-seq and Ribo-seq datasets and computed translational efficiency (TE, defined as Ribo-seq log2FC minus snRNA-seq log2FC) for each gene. Genes were classified as concordant (RNA and ribosome occupancy move together), translationally attenuated (RNA up, TE down) or translationally buffered (RNA down, TE maintained or elevated). NR2F6 generated a large translationally buffered set of 3,566 genes (Supplementary Table S17), substantially exceeding NR2F1 (1,449 genes; Fig. 8e). For these buffered genes, ribosome occupancy is maintained despite reduced mRNA, demonstrating active post-transcriptional preservation of protein output. Scatter plots of snRNA versus Ribo-seq log2FC confirm that the regression for the buffered set deviates toward the Ribo-seq axis, away from the concordance diagonal, with a lower transcription-translation correlation for NR2F6 (R = 0.26) than NR2F1 (R = 0.40), reflecting greater transcription-translation discordance in the NR2F6 condition (Fig. 8f).

GO analysis of the buffered genes (Fig. 8g) revealed enrichment for immune response, stress response and translation-related terms in both factors, with NR2F6 showing broader and more significant enrichment. These are precisely the functions a neuron must maintain at the protein level while its transcriptome is being reprogrammed: the stress-response and immune-regulatory programs need to remain online to manage the post-injury environment, even as bulk biosynthetic capacity is being restructured. Representative IGV tracks at buffered loci confirm maintained Ribo-seq signal alongside reduced snRNA signal at individual genes (Fig. 8h, Supplementary Fig. S12).

Importantly, this profiling captures a single early timepoint (7 days post-injury), not the full 8-to-12-week regenerative trajectory. The translational down-shift of the biosynthetic apparatus and the concurrent buffering of growth-relevant modules are best understood as a transient reallocation phase, in which the neuron pares back bulk translational load while protecting the programs needed to initiate and sustain axon growth. This interpretation is consistent with the sustained functional recovery observed over the following weeks (Fig. 5).

### NR2F1 and NR2F6 bind predominantly distal regulatory elements, re-deploying developmental enhancer landscapes while acquiring regeneration-specific targets

To map the genomic binding sites of NR2F1 and NR2F6 after injury, we performed CUT&RUN on GFP-positive CST nuclei isolated 14 days after thoracic crush from mice that had received AAV-NR2F6 or AAV-NR2F1 (Fig. 9a). For each factor, nuclei were incubated with a validated antibody against the native protein; a parallel no-antibody control was processed through the identical workflow and used as background for peak calling with MACS2 (the control track is displayed alongside the enrichment track in the genome-browser panels of Fig. 9f). Peak calling returned 1,096 high-confidence NR2F6 peaks and 493 NR2F1 peaks (Supplementary Table S18).

**Figure 9.**
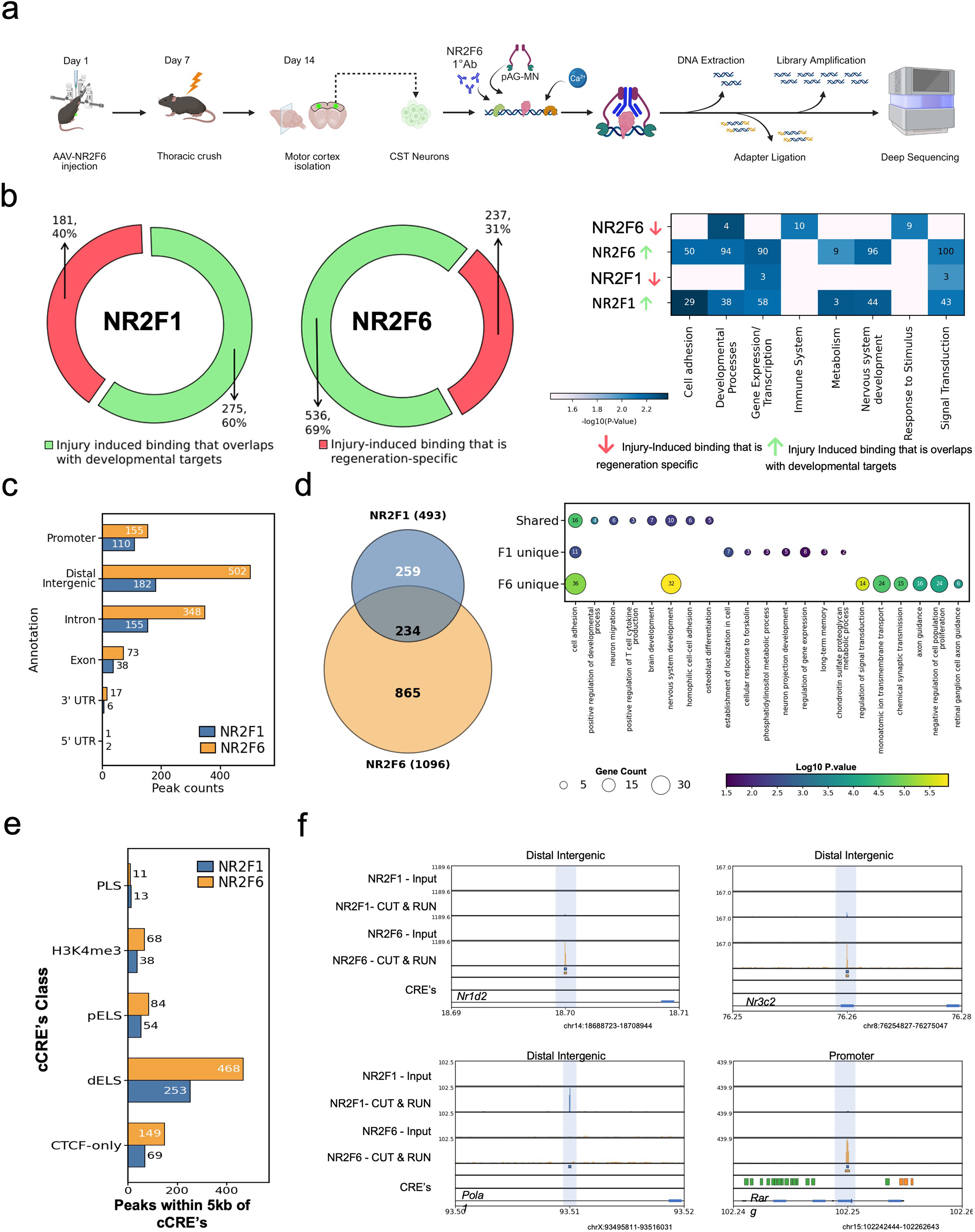
In vivo CUT&Run sequencing reveals the genomic binding landscape and regulatory elements targeted by NR2F1 and NR2F6. (a)Schematic representation of the in vivo Cleavage Under Targets and Release Using Nuclease (CUT&Run) experimental pipeline. The timeline depicts initial AAV injection (Day 1), subsequent thoracic crush injury (Day 7), and motor cortex isolation (Day 14). Following CST neuron isolation, targeted chromatin profiling was performed using transcription factor-specific primary antibodies followed by protein A/G-MNase targeted digestion, DNA extraction, and deep sequencing.(b) Donut plots for quantifying the NR2F1-bound (left) and NR2F6-bound (right) target genes categorized by their involvement in developmental ("Dev") versus non-developmental ("Non-Dev") processes. The accompanying heatmap (right) details the specific pathway enrichments associated with these developmental and non-developmental subsets for both transcription factors, with color intensity reflecting the - log10(P-Value) significance of the enrichment.(c) Bar chart detailing the genomic distribution of CUT&Run binding peaks for NR2F1 (blue) and NR2F6 (orange). Annotations map the precise localization of peaks across Promoters, Distal Intergenic regions, Introns, Exons, 3’ UTRs, and 5’ UTRs.(d) Venn diagram illustrating the extent of overlap between the identified target genes of NR2F1 and NR2F6, highlighting 234 shared targets alongside factor-specific unique targets. The adjacent dot plot categorizes the enriched biological processes associated with the shared, NR2F1-unique, and NR2F6-unique gene sets. Dot size represents the total gene count per pathway, while the color gradient corresponds to the statistical significance (Log10 P-value).(e) Bar graph quantifying the intersection of NR2F1 (blue) and NR2F6 (orange) binding peaks within a 5kb window of annotated candidate cis-Regulatory Elements (cCREs). Target regions are classified into specific regulatory signatures: promoter-like signatures (PLS), H3K4me3-rich regions, proximal enhancer-like signatures (pELS), distal enhancer-like signatures (dELS), and CTCF-only regions.(f) Representative genomic browser tracks confirming specific binding at select loci (e.g., Klf7, Neurog3, Nr1d2, Nr3c2, Pola1, Rarg). For each locus, CUT&Run signals from the targeted transcription factor ("T") are displayed alongside their respective input controls ("I"). The sequence tracks demonstrate highly specific peak enrichment normalized against background input. Corresponding cCRE annotations are displayed below the tracks to validate regulatory positioning.

Both factors bound predominantly distal regulatory elements rather than promoters (Fig. 9c). For NR2F6, the majority of peaks mapped to distal intergenic and intronic regions, with only a minority at promoters. NR2F1 showed a qualitatively similar distribution with a modestly higher promoter fraction. Intersection with ENCODE-annotated candidate cis-regulatory elements (cCREs) confirmed that peaks from both factors overlapped most frequently with distal enhancer-like signatures (dELS), followed by proximal enhancer-like signatures and H3K4me3-marked regions (Fig. 9e).

To ask whether the injury-induced binding recapitulates developmental NR2F occupancy, we compared each factor’s CUT&RUN peaks with the in-house curated embryonic COUP-TF target atlas (see Methods, Supplementary Table S19). For NR2F6, 69% of injury peaks overlapped developmental targets (Supplementary Table S20); the remaining 31% were unique to the injured adult cortex. For NR2F1, 60% overlapped developmental targets and 40% were regeneration-specific (Fig. 9b). Pathway enrichment of these two compartments (Fig. 9b, heatmap) showed that developmental-shared peaks in both factors were associated with cell adhesion, developmental processes, gene expression and transcription, nervous-system development and metabolism. Regeneration-specific peaks were enriched for immune-system, metabolic and signal-transduction functions, indicating that after injury both factors largely re-occupy their developmental enhancer landscapes while superimposing a new regulatory layer enriched for injury-relevant signaling and immune cues.

Although NR2F1 and NR2F6 bind at largely distinct genomic sites, they converge on a shared target set: 234 genes were bound by both factors (Fig. 9d). GO analysis of this shared set, alongside the NR2F1-unique and NR2F6-unique targets, revealed that the shared genes are enriched for cell adhesion, developmental regulation and nervous-system development, while NR2F6-unique targets are distinctively enriched for a wider range of processes including chondroitin-sulphate metabolism, axon guidance and negative regulation of cell proliferation (Fig. 9d, dot plot). Representative genome-browser tracks confirm specific binding at selected loci, with clear enrichment in the CUT&RUN signal relative to the matched input control (Fig. 9f).

To determine which binding events are accompanied by transcriptional change, we integrated the CUT&RUN data with the snRNA-seq differential-expression results (Supplementary Fig. S13). For NR2F6, of the bound genes testable in the snRNA-seq dataset, a defined subset showed concordant regulation (Supplementary Fig. S13a-b); the bound-and-regulated fraction was predominantly repressive (Supplementary Table S21), consistent with the transcriptional profile in Fig. 6. For NR2F1, a smaller fraction of bound genes was significantly regulated, with a more balanced direction of change (Supplementary Table S22). Most differentially expressed genes in both conditions were not directly bound, indicating that expression changes propagate indirectly through the upstream regulatory hubs identified in the Ribo-seq analysis (Fig. 8) and the three-dimensional chromatin reorganization described below (Fig. 10). Full gene lists for the bound-and-regulated, bound-not-regulated and regulated-not-bound categories are provided as supplementary tables for both factors (Supplementary tables 21,22).

**Figure 10.**
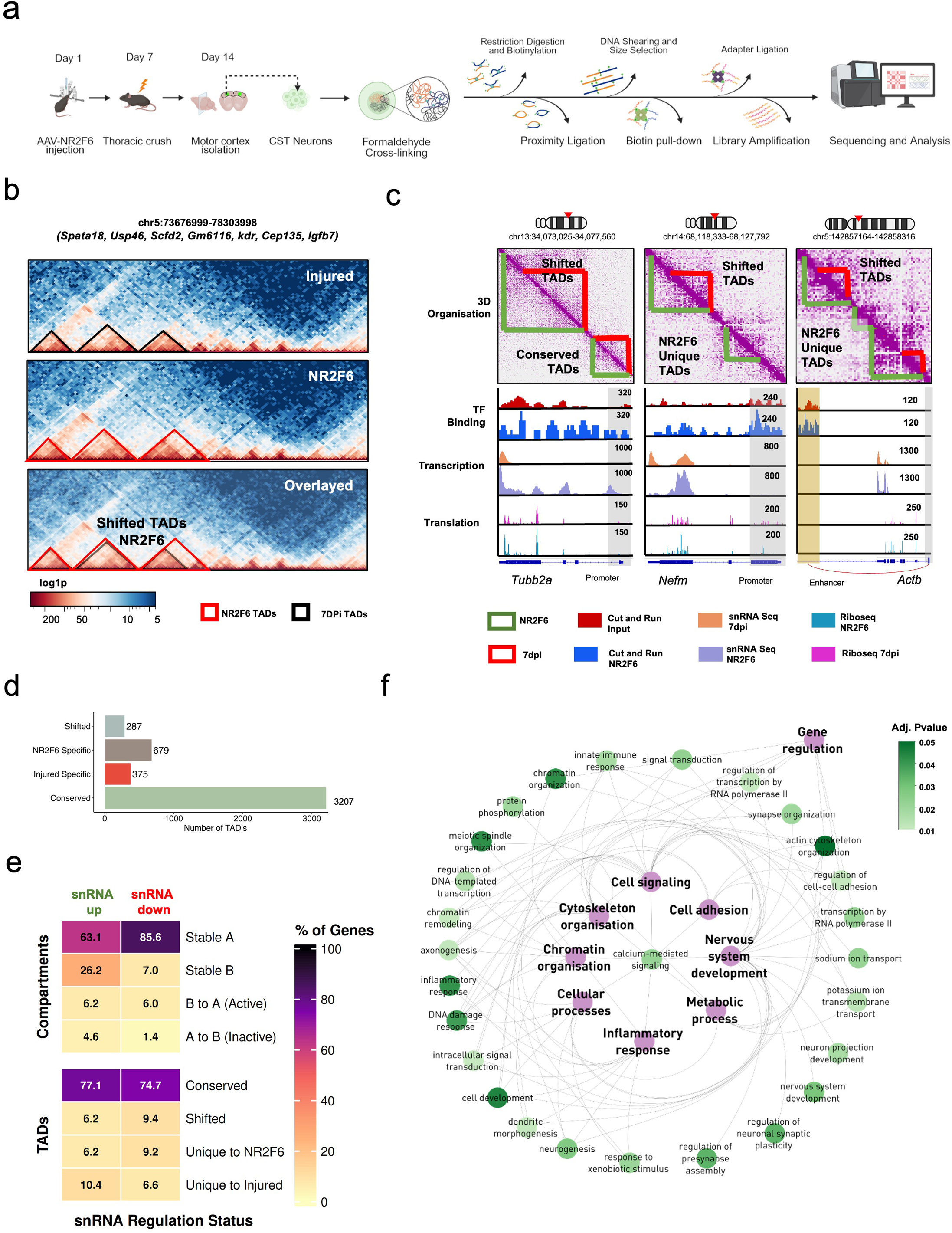
NR2F6-mediated chromatin reorganization following CNS injury revealed by Hi-C. (a) Schematic overview of the experimental workflow for Hi-C: Adult mice underwent AAV-mediated NR2F6 overexpression and spinal cord injury. Tissues were colected at Day 14 post-injury for chromatin conformation capture via in situ Hi-C followed by sequencing and downstream analysis. (b) Hi-C contact heatmaps depicting topologically associating domains (TADs) within a representative region of chromosome 5 (chr5:73,676,999–78,303,998) in Injured and NR2F6 samples. Overlayed maps show differential TAD boundaries: red outlines mark NR2F6-specific TADs, while black outlines mark TADs from the Injured condition. (c) Representative examples of TAD categories—Shifted, Conserved, and NR2F6- or Injured-specific TADs—alongside transcription factor (TF) binding profiles, transcription activity, and translation signals. Tracks show distinct epigenetic and transcriptional signatures associated with TAD reorganization.(d) Quantification of TAD classification across conditions: Conserved (n = 3207), NR2F6-specific (n = 679), Injured-specific (n = 375), and Shifted TADs (n = 287).(e) Heatmap integrating snRNA-seq differential expression data with Hi-C structural annotations. The color gradient and numerical values represent the percentage of differentially expressed genes (DEGs) mapping to specific global compartment dynamics (Stable A, Stable B, active transition [B to A], and inactive transition [A to B]) and TAD categories (Conserved, Shifted, Unique to NR2F6, and Unique to Injured).. (f) GO network visualization illustrating interconnections among selected GO terms enriched in NR2F6-specific TADs. Node color corresponds to term Adjusted P.value

Promoter occupancy showed essentially no correlation with expression change magnitude (NR2F6: Pearson R = -0.05 across 121 promoter-bound genes; NR2F1: R = +0.13 across 95 genes), consistent with regulation being determined by enhancer-target connectivity and three-dimensional genome organisation rather than by occupancy strength at the promoter (Supplementary Table S23).

### NR2F6 reorganises higher-order chromatin architecture, coupling new TAD formation and compartment switching to gene-expression changes

Because CUT&RUN showed that NR2F6 engages distal enhancers, we asked whether these binding events coincide with alterations in three-dimensional genome architecture. We carried out in-situ Hi-C on GFP-positive CST nuclei collected 7 days after injury from mice overexpressing NR2F6 (Fig. 10a). Injured GFP controls processed in parallel provided a baseline contact map.

Genome-wide insulation scoring identified 4,548 topologically associating domains (TADs) across both conditions. Most boundaries were stable: 3,207 TADs (70.6%) overlapped between datasets. However, NR2F6 introduced extensive remodeling: 679 entirely new TADs emerged in the NR2F6 condition, 375 were unique to the injured condition, and 287 boundaries shifted position (Fig. 10d, Supplementary Table S24). At a representative locus on chromosome 5, NR2F6 gained discrete sub-TADs that split a broad injury domain into finer compartments (Fig. 10b).

These structural changes are not inert: they are directly coupled to gene-expression alterations at multiple levels. At the locus level, integrative tracks (Fig. 10c) show this chain explicitly. At Tubb2a and Nefm, NR2F6 binding is present at new internal TAD boundaries, and at the same loci transcription (snRNA-seq) and ribosome occupancy (Ribo-seq) both rise. At Actb, which lies within a conserved TAD, there is no NR2F6 recruitment and no transcriptional change. The contrast between reorganised and conserved loci, with matched binding, transcription and translation tracks, demonstrates that boundary formation is spatially aligned with altered gene output.

At the genome-wide level, we integrated the Hi-C structural annotations with the snRNA-seq differential-expression data (Fig. 10e). This analysis reveals the relationship between chromatin reorganization and transcriptional change at two scales. First, at the compartment level: NR2F6 overexpression switched 3,783 genomic bins between A (active) and B (inactive) compartments relative to injured controls, approximately balanced between activating B-to-A transitions and repressive A-to-B transitions (Supplementary Table S25). Genes in B-to-A switching regions were enriched among the snRNA-seq up-regulated set, and genes in A-to-B regions among the down-regulated set, confirming that compartment relocation is associated with the expected transcriptional consequences. Second, at the TAD level: genes embedded in NR2F6-specific TADs were preferentially associated with transcriptional activation (Supplementary Table S26, S27), including regeneration-relevant genes such as Robo1, Nefm, Sptbn4 and Cacna1a, while genes in shifted TADs included hallmark growth genes such as Tubb3.

Gene Ontology analysis of the NR2F6-specific TADs (Fig. 10f, Supplementary Table S28) showed enrichment for chromatin remodeling, cell signaling, cytoskeleton organisation, intracellular signal transduction, nervous-system development and inflammatory response, with dense interconnections among these functional nodes. These are the same categories enriched in the snRNA-seq and CUT&RUN data (Figs. 6, 9), indicating convergent regulation across binding, expression and chromatin-architecture levels.

The Hi-C data show that NR2F6 overexpression reorganizes chromatin at two hierarchical scales: forming new TADs that embed growth-associated genes with their regulatory elements, and switching A/B compartments to relocate genes between active and inactive nuclear environments. These structural changes are accompanied by, and spatially aligned with, the corresponding transcriptional and translational alterations documented in Figs. 6 and 8, providing a coherent mechanistic chain from enhancer binding (Fig. 9) through three-dimensional reorganization to gene output.

## Discussion

We identify NR2F1 and NR2F6 as previously unrecognized nuclear-receptor transcription factors that promote adult corticospinal regeneration. Forced expression of either factor significantly increases neurite outgrowth in single-neuron tracing assays and, in two distinct spinal-cord injury models, supports long-tract axon growth with measurable motor recovery. Parallel multi-omic profiling of both factors (CUT&RUN, snRNA-seq, Ribo-seq) along with Hi-C of NR2F6 reveals distinct modes of action: NR2F1 re-engages chromatin-remodeling and cytoskeletal programs; NR2F6 re-occupies developmental enhancer targets, reorganizes three-dimensional chromatin architecture, and imposes a translational down-shift that selectively preserves growth-relevant modules through translational buffering. We present an integrated working model (Fig. 11) linking developmental enhancer closure, injury-induced re-expression, and the structural and metabolic reallocation that follows.

**Figure 11.**
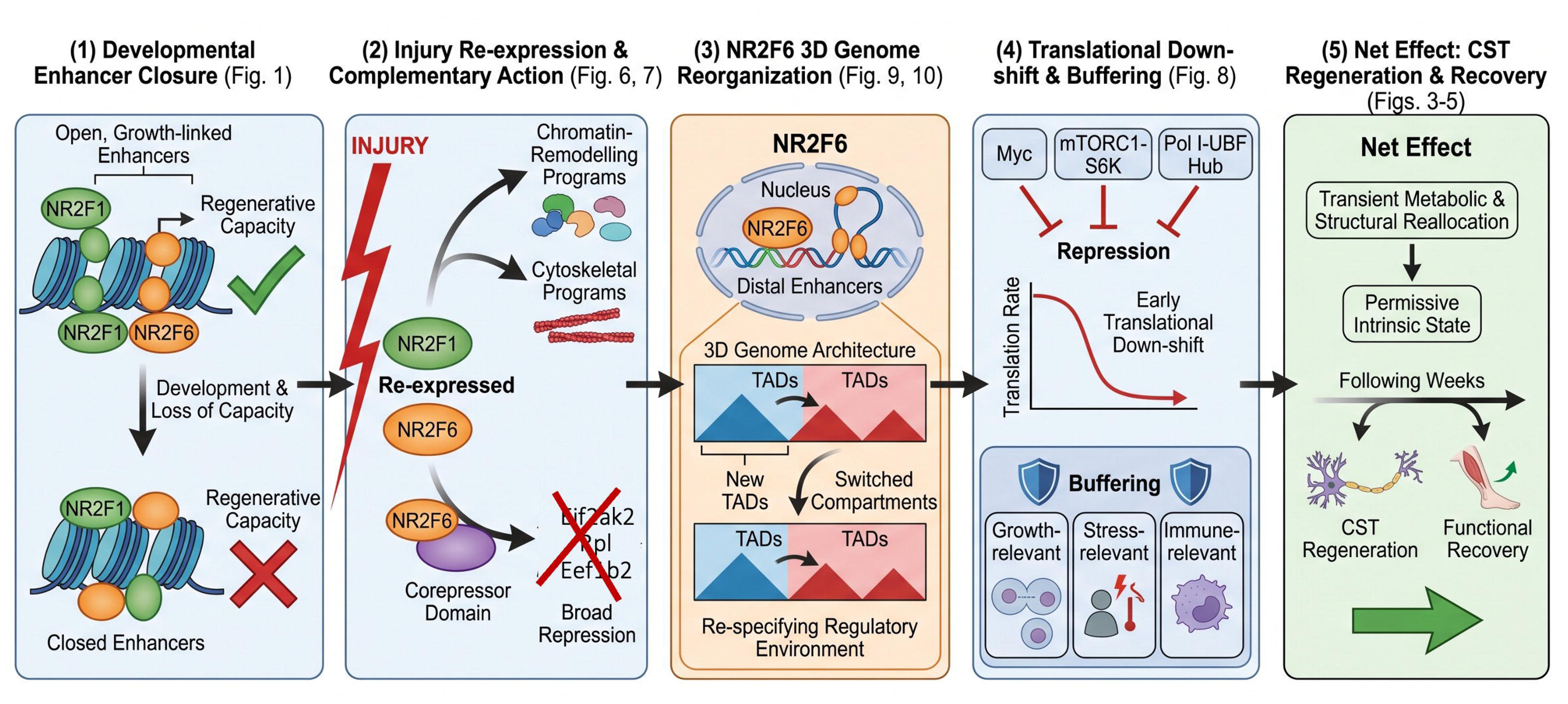
Integrated working model for NR2F1- and NR2F6-driven corticospinal regeneration. Schematic summarising how NR2F1 and NR2F6 reactivate intrinsic growth competence in the adult corticospinal tract, integrating the data across all figures. (1) Developmental enhancer closure (Fig. 1). During cortical development, NR2F1 and NR2F6 footprints occupy open, growth-linked enhancers in regeneration-competent neurons. As neurons mature these enhancers progressively close and regenerative capacity is lost, defining the pro-growth program that the factors are positioned to re-engage. (2) Injury re-expression and complementary action (Figs. 6, 7). Re-expressed after spinal cord injury, NR2F1 and NR2F6 act through complementary routes. NR2F1 reactivates chromatin-remodelling and cytoskeletal programs, while NR2F6, through its conserved corepressor domain, drives broad transcriptional repression. (3) NR2F6-mediated 3D genome reorganisation (Figs. 9, 10). NR2F6 binds predominantly distal enhancers and reorganises higher-order chromatin architecture, generating new topologically associating domains (TADs) and switching A/B compartments. These structural changes re-specify the regulatory environment of the enclosed genes. (4) Translational down-shift and buffering (Fig. 8). Repression of the upstream regulatory hub, comprising Myc, the mTORC1-S6K axis and the Pol I -UBF apparatus, produces an early genome-wide down-shift in translation. Translational buffering selectively preserves protein output from growth-, stress- and immune-relevant modules despite reduced transcript levels. (5) Net effect: CST regeneration and functional recovery (Figs. 3-5). The combined action produces a transient metabolic and structural reallocation that establishes a permissive intrinsic state. Over the following weeks this supports corticospinal tract regeneration and the recovery of motor function.

NR2F1 (COUP-TFI) is a well-established architect of brain development, directing cortical arealisation, progenitor maturation, layer specification, thalamocortical and callosal targeting, and postnatal synaptogenesis; human haplo-insufficiency underscores its importance for higher-order circuitry, yet its influence on injured adult axons has not been tested(Armentano *et al*, 2007; Bertacchi *et al*, 2019; Tomassy *et al*, 2010). NR2F6 (Ear2) is best known as an intracellular checkpoint that restrains effector T-cell activity and shapes anti-tumor immunity, with neural evidence limited to a knockout study showing disrupted locus-coeruleus formation, circadian instability and altered nociception (Hermann-Kleiter *et al*, 2015; Warnecke *et al*, 2005). Beyond this isolated developmental role, NR2F6 has no documented function in mature CNS neurons, and neither factor has been explored for the capacity to drive axon regeneration.

Our discovery of NR2F1 and NR2F6 emerged from a strategy that mines developmental chromatin landscapes for transcription factors that co-occupy pro-growth regulatory DNA. In earlier work, we anchored the search to KLF6 binding sites and uncovered the cooperative pairing of KLF6 with NR5A2, a combination that proved effective in enhancing corticospinal sprouting in vivo (Wang *et al*, 2018; Venkatesh *et al*, 2021). In the present study we removed the anchor constraint and scanned the full catalogue of developmentally open, growth-linked enhancers, revealing a convergence of NR2F1 and NR2F6 footprints on the same elements. Consistent with a developmental origin for this regulatory programs, both factors are expressed at high levels during the growth-competent period (E11 through P0) and are markedly reduced in the adult, with Nr2f6 declining approximately 78% and Nr2f1 approximately 69% from their respective developmental maxima (Supplementary Fig. S1). Importantly, Nr2f1 remains elevated throughout the perinatal period, when intrinsic axon-growth capacity is still intact, and drops only postnatally, a trajectory consistent with the known timeline of regenerative decline. After injury-induced overexpression, 69.3% of NR2F6 and 60% of NR2F1 CUT&RUN peaks overlap their respective developmental target sets (Fig. 9), and 25.1% of NR2F6 developmental targets are significantly differentially expressed after overexpression, indicating that re-expression re-engages a substantial fraction of the developmental programme at the transcriptional level. The present study therefore reinforces enhancer co-occupancy as a productive strategy for identifying transcription-factor combinations capable of promoting CNS repair.

NR2F1 and NR2F6 achieve comparable functional recovery (Fig. 5) through different molecular strategies. NR2F1 modulates 823 transcripts with a balanced activation/repression profile (430 up, 393 down), enriched for chromatin organisation, cytoskeletal remodeling and transcriptional regulation. NR2F6 drives a far larger transcriptional shift (4,076 genes, 93% repressed) and generates a substantially larger translationally buffered gene set (3,566 genes versus 1,449 for NR2F1). The CUT&RUN data reinforce this distinction: NR2F6 binds 1,096 sites (predominantly distal enhancers), while NR2F1 binds 493 with a modestly higher promoter fraction. Of the bound genes testable in the snRNA-seq, a defined subset shows concordant regulation, predominantly repressive for NR2F6 and more balanced for NR2F1 (Supplementary Fig. S13), while most differentially expressed genes are not directly bound, indicating that expression changes propagate indirectly through the upstream regulatory hub and three-dimensional genome reorganization described below. That two factors with such different molecular footprints produce equivalent functional outcomes suggests the regenerative state can be reached through more than one transcriptional route. This has direct implications for combinatorial gene-therapy design.

Although regeneration is usually framed as a high-energy process fueled by surges in protein synthesis, evidence from several systems points in a different direction: a transient, system-wide reduction of translation can free metabolic bandwidth and limit proteotoxic stress, focusing the cell on specialized growth programs. Quiescent muscle stem cells preserve their long-term regenerative capacity by maintaining eIF2α-mediated inhibition of global translation until commitment cues arrive (Eliazer & Brack, 2016); transient pharmacological dampening of the mTORC1/eIF4F axis improves functional recovery in the injured heart(Hofmann *et al*, 2024); and integrated-stress responses that throttle initiation rates are activated across multiple repair-competent vertebrate tissues(Wek, 2018). In the nervous system, however, transcriptional repression has historically been cast as an intrinsic brake, exemplified by HHEX(Simpson *et al*, 2015) and KLF9 (Moore *et al*, 2009), whose post-injury up-regulation coincides with regenerative failure and whose knock-down enhances neurite extension. Our data show that NR2F6 promotes CNS axon growth precisely because it represses. Acting through a conserved corepressor motif (LGIDNVCEL, residues 184-192), NR2F6 does not silence individual translation-gene promoters; none of the approximately 180 translational-apparatus genes whose ribosome occupancy falls carry an NR2F6 peak. Instead, NR2F6 binds distal enhancers and, through repression of the upstream Myc, mTORC1-S6K and Pol I -UBF regulatory hub (Supplementary Table S29), produces a coordinated down-shift of 79 ribosomal-protein genes, 42 initiation factors, 33 tRNA synthetases and 27 elongation factors, while preserving genes linked to cytoskeletal assembly, energy production and signaling. However, this is not a blanket shutdown, translational buffering maintains protein output from 3,566 genes, enriched for immune-response, stress-response and growth-relevant functions, even as their mRNA levels fall (Fig. 8, Supplementary Fig. S12). By reducing bulk biosynthetic load while protecting growth-relevant modules, NR2F6 reroutes cellular resources toward the specific task of rebuilding axons rather than sustaining general biosynthesis. Transcriptional repression, then, is not simply a barrier to regeneration; in the NR2F6 context it actively facilitates regrowth. Translational down-scaling coupled with selective buffering is a previously unrecognized mechanism for creating a permissive metabolic state in the injured CNS.

Recent work has placed long-range chromatin architecture, rather than promoter-proximal control, at the center of axon regeneration. In dorsal-root-ganglion neurons, regenerative competence correlates with selective opening of thousands of enhancer elements that remain closed when their regeneration-incompetent central branches are lesioned (Palmisano *et al*, 2019). We previously showed that transcription factors whose motifs populate both promoters and enhancers are disproportionately enriched among growth-promoting hits, exemplified by the Klf6/Nr5a2 pair uncovered through a co-occupancy pipeline(Venkatesh *et al*, 2021). Recent three-dimensional chromatin maps of sensory neurons revealed that injury induces complex promoter-enhancer loops and that cohesin is required for full activation of the regenerative programme, directly linking higher-order topology to axon growth(Palmisano *et al*, 2024). Our present data add a CNS counterpart: CUT&RUN shows that NR2F6 binds predominantly distal enhancers, and Hi-C demonstrates that it reorganizes the injured corticospinal genome into 679 new TADs and shifts 287 boundaries, while switching 3,783 genomic bins between A (active) and B (inactive) compartments. Promoter occupancy shows essentially no correlation with expression-change magnitude (NR2F6 Pearson R = -0.05 across 121 promoter-bound genes; NR2F1 R = +0.13 across 95 genes), confirming that regulation is determined by enhancer-target connectivity and three-dimensional organisation rather than by occupancy strength at the promoter. Genes within NR2F6-specific TADs are preferentially associated with transcriptional activation of regeneration-relevant targets, and compartment switching relocates loci between active and inactive environments with corresponding expression consequences (Fig. 10). Together, these findings show that enhancer recruitment and three-dimensional genome repackaging are mechanisms by which corticospinal neurons re-engage developmental growth circuitry after injury.

We discuss here a few caveats that bear on interpretation of the multi-omic data. The snRNA-seq, Ribo-seq and CUT&RUN profiles were collected at a single early timepoint (7 days post-injury) and therefore represent the initial transcriptional and translational response of the neuron, not the full 8-to-12-week trajectory over which functional recovery accrues. The early translational down-shift and the concurrent buffering should be read as a transient reallocation phase; whether this resolves into a later biosynthetic expansion as regeneration proceeds will require longitudinal profiling to determine. Because AAV was delivered by intracortical injection rather than retrograde tracing, the profiled population comprises layer-V extratelencephalic projection neurons of the motor cortex, of which corticospinal neurons are the principal but not exclusive component; the in vivo axon-tracing data (Figs. 3, 4), in which GFP-labelled axons project through the medullary pyramids into the spinal cord, confirm that genuine corticospinal neurons are present in the transduced population, but isolation of a pure corticospinal set would require retrograde labelling and is an informative direction for future work. Finally, the snRNA-seq was performed on a pooled sample per condition; our use of stringent effect-size and significance thresholds and the independent corroboration of the transcriptional findings by Ribo-seq, CUT&RUN and Hi-C provide confidence in the reported gene sets, though formal pseudobulk inference across biological replicates would strengthen the statistical framework.

In summary, we identify NR2F1 and NR2F6 as regulators of adult corticospinal repair, track their impact from developmental enhancer co-occupancy through in vitro validation, two in vivo injury models and parallel multi-omic dissection, and present an integrated working model (Fig. 11) linking enhancer redeployment, translational reprogramming and three-dimensional genome reorganization to axon regrowth. NR2F1 reactivates chromatin-remodeling and cytoskeletal modules; NR2F6, through a conserved corepressor motif, re-occupies developmental enhancers, restructures the translational economy through selective buffering, and reorganizes chromatin into growth-permissive domains. Their different strategies show that the regenerative state in corticospinal neurons can be reached through more than one transcriptional route and provide a mechanistic framework for designing transcription-factor-based interventions for spinal-cord repair.

## METHODS

### Animal husbandry

All the animal procedures were approved by the Institutional Animal Ethics Committee (IAEC) at CSIR-CCMB (IAEC 09/2025, IAEC 07/2025) Animals were housed under a 12-hour day and night cycle. All experiments were performed on wild-type C57BL/6J mice (mixed population).

### Cloning strategy

Mouse NR2F6 [Horizon Discovery, #MMM1013-202762257] and NR2F1 [Horizon Discovery, #MMM1013-202739977] were isolated from a cDNA library of mouse transcription factors. NR2F1 was cloned using traditional cut and paste cloning using 3’ HindIII [NEB, #R0104S] and 5’ KpnI [NEB, #R3142L]. NR2F6 gene was cloned into an AAV- compatible plasmid containing a CAG promoter [Addgene Plasmid, #59462] using standard PCR amplification protocol (98°C for 3 minutes, 95°C for 30 seconds, 59°C for 30 seconds, 72°C for 1.10 minutes, and a final extension at 72°C for 5 minutes for 35 cycles), PCR- amplified NR2F6 was digested with compatible restriction enzymes 3’ EcoRI [NEB, #R3101S] and 5’ XbaI [NEB, #R0145S]. The tdTomato reporter gene was excised from the plasmid backbone and replaced with the amplified gene using T4 DNA ligase [Takara, #2011B]. Successful cloning was confirmed by Sanger sequencing (CCMB facility) using CAG-forward (5’-GCAACGTGCTGGTTATTGTG-3’) and Amp-reverse (5’-ATAATACCGCGCCACATAGC- 3’) primers. A complete list of plasmids and primers used in this study is provided in Supplementary Table S32.

### AAV production

Adeno-associated viral particles (serotype 9) were produced in HEK293T cells and purified using a commercial AAV purification kit [Takara, #6675]. HEK293T cells were transiently transfected with pAAV-CAG-NR2F1 or pAAV-CAG-NR2F6, pAAV2/9n, and pAdDeltaF6 plasmids (gifted by Regalla lab, CSIR-CCMB) using polyethylenimine 40k [Polysciences, #24765]. 72 hours post-transfection, cells were harvested and viral isolation was performed according to the manufacturer’s instructions. AAV capsid integrity was assessed by 10% SDS-PAGE, run at 35 mA for 120 minutes, followed by Coomassie Blue staining for 20 minutes and overnight destaining in a methanol-acetic acid solution. For viral titer quantification, AAV particles were first treated with DNase I [Thermofisher, #EN0521] for 30 minutes at 37°C, then serially diluted ten-fold across four dilutions. Quantitative PCR was performed using TB Green [Takara, #RR82WR] to estimate viral titers by comparison with a plasmid standard curve. All AAV preparations achieved titers of 10 viral genomes/µl or higher, with intact capsid formation confirmed by SDS-PAGE analysis (supplementary figure S2).

### Primary Neuron Culture and Transduction

Acid-treated, autoclaved coverslips in a 12- well plate were coated with Poly-D-Lysine [Gibco, #A3890401] at 0.2 mg/mL and incubated at 37°C for 2 hours. After washing with autoclaved Milli-Q water, the coverslips were air- dried in a hood. C57BL/6J P0 pups were sacrificed according to IAEC guidelines. Motor cortices were isolated in pre-warmed HBSS (Hank’s Balanced Salt Solution), minced, and incubated in 0.1% Trypsin-EDTA [Sigma, #T4799] at 37°C for 10 minutes. Trypsin activity was stopped with Fetal Bovine Serum [Gibco, #10438026], followed by trituration in plating media (Neurobasal medium [Gibco, #21103049], 2% B27 supplement [Gibco, #17504044], 5% Fetal Bovine Serum, 1X PenStrep). Cells were counted and plated at low density of 35,000 cells per well in 1 mL plating media. After 24 hours, 50% of the media was replaced with fresh media containing 1 µM Ara-C [Sigma, #C1768]. Neurons were co-transduced with pAAV-CAG-GFP and the virus of interest (1:2). Transduction efficiency was assessed at 50 hours post-transduction, followed by immunostaining with βIII-Tubulin [CST, #2128S]

### Neuro-2a culture for in vitro neurite outgrowth assays

Neuro-2a cells [ATCC, #CCL-131] were cultured in DMEM [Gibco #11885084] supplemented with 10% FBS and 1X penicillin-streptomycin-gentamicin (PSG). For transfection, 15,000 cells were seeded per well in 96-well plates and incubated for 24 hours prior to transduction with AAV-NR2F1, AAV-NR2F6, or a combination of AAV-NR2F1 and AAV-NR2F6. In all conditions, the viruses were co-transduced with a tracer AAV-GFP at a 3:1 ratio. After 48 hours, cells were trypsinized with 10 µL of 0.1% trypsin [Sigma, #T4799] and reseeded onto 18 mm X 18 mm glass coverslips in 12-well plates. Following cell adhesion, the medium was replaced with a differentiation medium (DMEM and PSG) for 48 hours to promote neurite outgrowth. Cells were then fixed with 1 mL of 4% paraformaldehyde [Sigma, #30525-89-4] for 15 minutes and immunostained. Neurite outgrowth was visualized using a Zeiss Apotome. 2 microscopy system (20x magnification) with the following fluorescent markers: βIII-tubulin (1:500 dilution; TRITC filter,ex 557/576 nm, em 580/620 nm), GFP (FITC filter, ex 475/495 nm, em 510/525 nm), and nuclei (DAPI filter, ex 357/44 nm, em 447/60 nm). All image acquisition and quantification were performed in a double-blinded manner, with experimental groups coded until after completion of analysis and statistical testing. Neurite length was quantified by tracing polylines along neurites of transfected cells from overlay images(Supplementary Table S4). Data analysis was performed using GraphPad Prism (version 10.1.0), with statistical significance assessed by Kruskal-Wallis test followed by Dunn’s multiple comparisons. Results are presented as mean ± SEM.

### Animal Surgeries

All animal procedures were approved by the Animal Ethics Committee at CSIR CCMB and adhered to the IAEC guidelines. Adult C57BL/6J mice of mixed sex (>12 weeks old, 20-25g) were used for cortical injections and thoracic crush injury procedures, while 6–8-week-old mice were used for pyramidotomy. For all procedures, mice were anesthetized with a mixture of Ketamine (50 mg/ml) and Xylazine (100 mg/ml). For cortical injections, after shaving and incising the forehead, Bregma was identified. Using a stereotaxic apparatus, four injection sites were marked relative to Bregma (in mm): (M/L=+2, A/P=0), (M/L=+2, A/P=+1), (M/L=-2, A/P=0), and (M/L=-2, A/P=+1). Small holes were drilled at these sites, and a Hamilton syringe was used to deliver the virus at a depth of D/V=0.50-0.60 mm. For each condition, AAV vectors were mixed at a 3:1 volumetric ratio (transgene vector to GFP reporter) prior to injection: AAV-NR2F6 or AAV-NR2F1 with AAV-GFP for single-factor conditions, and AAV-NR2F6 with AAV-NR2F1 and AAV-GFP at a 3:3:1 ratio for the combined condition. All vectors were titre-matched before mixing. The GFP reporter, delivered at the minority ratio, served to identify transduced neurons without competing for viral-particle uptake with the transgene vector. Control animals received AAV-GFP alone at the equivalent total volume. All viral constructs and titer are summarized in Supplementary Table S31. For the thoracic crush injury, a midline incision exposed the spinal column, followed by blunt muscle dissection. Mice were secured in a custom spine stabilizer, and a laminectomy was performed at the T6-T10 vertebrae using spring scissors to expose the spinal cord. The spinal cord was then carefully crushed with equal force using blunt forceps (width-1mm). The pyramidotomy procedure was adapted from previous published work(Starkey *et al*, 2005). Briefly, a ventral midline incision was made between the forelimbs and jaw, and the medullary pyramid was exposed by partial removal of the occipital bone. The dura was cut, and the right pyramidal tract was incised using iridectomy scissors (0.5 mm width, 0.25 mm depth). Post-surgery, the oesophagus, trachea, and muscles were repositioned, and the skin was sutured.

### Axon-growth quantification: Pyramidotomy

Maximum of three 50 µm transverse sections per animal (C2 → C6; 1.5 mm spacing) were imaged on a Leica STED (63×, tile scan in 520 × 520 pixel, 1 µm z-steps, 400 frames/second ; intensity 1,250 V, laser power 40 %). Maximum-intensity projections were generated, a midline through the central canal was drawn, and parallel 10 µm-wide virtual lines were placed 200, 400 and 600 µm from the midline. GFP⁺ axons intersecting each line were counted. To normalise for tracer uptake, 50 µm vibratome sections of the medullary pyramids were imaged on a Zeiss ApoTome.2 at 20X magnification. Individual dots in the medullary section were counted on ImageJ for the whole medullary section. Maximum of 3 medullary section were taken whose average was taken final medulla count. All image acquisition and quantification were performed in a double-blinded manner, with experimental groups coded until after completion of analysis and statistical testing Lesion completeness was verified by PKCγ immunohistochemistry. PKCγ is selectively expressed in corticospinal tract neurons and their axons; following axotomy, the transected axons undergo Wallerian degeneration and lose PKCγ immunoreactivity, so its absence on the lesioned side, with retained staining on the intact side as an internal control, confirms a complete lesion.(Mori *et al*, 1990)

### Thoracic crush

Tissue processing matched the pyramidotomy protocol. Sections (50 µm) spanning the lesion were imaged on the same STED (100× objective). The lesion centre was identified as the peak of GFAP immunoreactivity, a reference line was drawn through this point, and additional 10 µm-wide lines were placed 500, 1,000 and 1,500 µm caudal to it. GFP⁺ axons crossing each line were counted and normalised to pyramid counts. To normalise for tracer uptake, 50 µm vibratome sections of the medullary pyramids were imaged on a Zeiss ApoTome.2 at 20X magnification. Individual dots in the medullary section were counted on ImageJ for the whole medullary section. In case of regeneration experiment since both side are labelled with tracer virus we imaged both the medulla from the slide and calculated by ImageJ. After calculating both medulla the medulla count from both the slide were added to give total axons labelled. All image acquisition and quantification were performed in a double-blinded manner, with experimental groups coded until after completion of analysis and statistical testing

### Functional Enrichment and Gene Ontology (GO) Analysis

To elucidate the biological relevance and functional architecture of the filtered gene list, over-representation analysis was performed using the Database for Annotation, Visualization and Integrated Discovery (DAVID) web server.The filtered gene symbols were uploaded into the DAVID interface and evaluated against the *Mus musculus* genomic background. Functional annotation was strictly restricted to the Biological Process (BP) domain of Gene Ontology (GO). Statistical significance for enriched biological themes was determined using a threshold of P <= 0.05, derived from the internal EASE score (a modified Fisher’s exact test). To minimize functional redundancy and provide a high-level biological overview, the significantly enriched specific GO identifiers (GO IDs) were hierarchically mapped to identify their overarching parent terms (edges are linear relationships). This lineage-based categorization was utilized to synthesize and consolidate the primary biological themes represented by the gene dataset. This lineage-based categorization is visualized as the network in Fig. 1a, where edges represent parent-child relationships within the GO hierarchy and node color encodes enrichment P-value. Full enrichment output is provided in Supplementary Table S1.

### Single-nucleus RNA-seq

C57BL/6 mice (8–10 wk, mixed sex) were injected anterogradely with either AAV-GFP (control) or AAV-NR2F1 + AAV-NR2F6 (0.6 µL per site; four sites: ±2 mm ML, 0- and +1-mm AP relative to bregma). One week later the animals received a T8 crush; motor cortices were collected 7 d after injury, flash-frozen and stored at –80 °C. Tissue was dounced in 2 mL Nuclei Lysis Buffer with RNase inhibitor, nuclei pelleted (500 g, 2 min, 4 °C), resuspended in Nuclear Suspension Buffer (PBS + 0.1 % BSA + RNase inhibitor) and passed through 20 µm mesh. GFP-positive nuclei were flow-sorted on a BD Melody (80 µm nozzle; debris and doublets excluded by FSC/SSC gating) into resuspension buffer (4500–5 000 events per sample). Single-nucleus libraries were generated with the 10x Genomics Chromium Next GEM v3.1 kit [#PN1000121] following the manufacturer’s protocol.

### CUT&RUN assays

Motor cortices were harvested 7 d after T8 crush from mice pre-injected with AAV9-NR2F6 and AAV9-NR2F1. Tissue was minced on ice, fixed in 0.1 % formaldehyde [CST, #12606P] for 2 min, quenched with 10X glycine [CST, #7005S] and washed. All subsequent steps followed the DNA–Protein Interaction Assay Kit [CUT&RUN CST, #86652]. Cells were dounced in kit wash buffer supplemented with spermidine [CST, #27287] and Protease Inhibitor Cocktail [CST, #7012], bound to Concanavalin-A magnetic beads, and incubated overnight at 4 °C with anti-NR2F6 [Thermofisher, #PA5-23321] and anti-NR2F1 (Abcam,# ab181137). Antibodies target exogenous NR2F6/NR2F1 protein, endogenous levels are very low in adult murine brain. Bead–nuclei complexes were washed and exposed to pAG-MNase [CST, #15338] in digitonin buffer [CST, #16359] containing spermidine and protease inhibitors for 1 h, then activated with CaCl₂ for 30 min at 4 °C. Digestion was stopped with kit Stop Buffer [CST, #48105] plus RNase A [CST, #7013], and cross-links were reversed with 10 % SDS and Proteinase K [CST, #10012] at 65 °C for 2 h. DNA was purified using kit spin columns [CST, #14209]. A no-antibody negative control was processed through the identical workflow, omitting only the primary antibody, to provide a matched background for peak calling; nuclei were lysed in DNA Extraction Buffer [CST, #42015], sonicated to 100–600 bp on a Covaris M220 (peak 75, duty 10, 200 cycles, 75 s) and processed in parallel. Libraries for both CUT&RUN and input samples were prepared with the NEBNext® Ultra™ II DNA Library Prep Kit [NEB, #E7103] and sequenced to ∼5 million paired end reads per sample.

### Ribosome profiling

Ribosome profiling protocol was adapted from previously published work(Froberg *et al*, 2023). Motor cortices were dissected from anaesthetised C57BL/6 mice 7 d after AAV9- NR2F6 injection and T8 crush, placed in ice-cold HBSS, and enzymatically dissociated in 0.1% trypsin-EDTA containing RNase-free DNase I (15 µL / 5 mL) at 37 °C for 20 min. Trypsin was quenched with FBS, tissue was transferred to Neurobasal medium and triturated sequentially through flame-polished pipettes. Cell suspensions were spun (2000 r.p.m., 15 s), supernatants pooled, and GFP⁺ cells isolated on a BD FACS Melody without nuclear dyes; ∼1 µg total RNA per animal was obtained. Sorted cells were resuspended in 2X polysome buffer (final 3 mL) and digested with RNase I [Promega, #N2515] (12 U mL⁻¹, 45 min, RT, rocking). Digestion was quenched with 10 µL of Superasin [Thermofisher Scientific, #2696], and lysates concentrated to ∼400 µL in Amicon Ultra-15 filters, (30 kDa, NMWL filter, EMD Millipore #UFC903024; 5000×g, 4°C). RNA was extracted using RNeasy Mini Kit [Qiagen, #74104]. Ribosome-protected fragments (25–37 nt) were size-selected on TBE–urea PAGE, visualised with SYBR Gold [Invitrogen, #S11494], excised, eluted overnight in 300 mM KOAc/1 mM EDTA, precipitated with isopropanol and re-purified on RNeasy columns. Fragments were 5Ö- phosphorylated with T4 PNK [NEB, #M0201S]; 50 µL reaction, 30 min, 37 °C) and cleaned on RNeasy. Libraries were prepared with the QIAseq miRNA Library Kit [Qiagen, #331502] and concentrated to 5 µL by speed-vac. For bulk polysome profiling, intact cortices were homogenised in ice-cold 1× polysome buffer + cycloheximide [Millipore Sigma, # C4859] (1 µg mL⁻¹) using 11 strokes of a chilled dounce. Post-nuclear supernatants (1660×g, 15 min, 4 °C) were brought to 6 mL, digested with RNase I (200–500 U, 1–2 h, RT) and stopped with Superasin. Lysates were concentrated to ∼400 µL (Amicon Ultra-15, 30 kDa) and RNA processed as above for library preparation.

### Behavioral assessments

For behavioural assessment, mice were familiarised with a horizontal ladder-walking task on two occasions, 7 days before and 3 days after stereotaxic AAV delivery to eliminate novelty effects. The apparatus consisted of a 23 × 2 × 6-inch aluminium frame fitted with evenly spaced rungs (1-cm centre-to-centre) and flanked by clear acrylic walls that kept the animal centred in the camera field. Beginning 7 days after thoracic crush and continuing at weekly intervals through week 9, each animal performed three uninterrupted traversals per session while being filmed at 30 fps with a side-mounted high-speed camera aligned to the ladder mid-line. Raw MOV files were imported into openshot video editor, and in every tenth frame the vertical distance from the greater trochanter (hip) to a baseline drawn through the tarsal joints was measured, yielding a mean hip-lift value that distinguished active limb elevation from passive dragging. For detailed kinematic analysis, the same videos were processed with DeepLabCut v3.1(Aljovic *et al*, 2022). For grip strength the mouse was held by the experimenter and gently brought toward the horizontal rod attached to the grip strength meter. As the animal approaches the rod, it reflexively attempts to push against it using its hindlimbs, generating a measurable compression force. The grip strength meter records the peak compression force in grams-force at the moment the animal is pushing away the rod. This value is then normalized to the body weight yielding normalized grip strength.

### Gait kinematic analysis

Kinematic analysis of hindlimb gait was performed using DeepLabCut (DLC, v3.1). Video recordings of mice walking on a horizontal ladder rung apparatus were acquired from a lateral perspective at 30 fps. A subset of frames (n = 200) was used to train a ResNet-50 deep neural network to track six anatomical landmarks on the hindlimb: iliac crest, hip, knee, ankle, metatarsophalangeal (MTP) joint and toe. Only frames with a tracking likelihood greater than 0.9 for all landmarks were retained for kinematic analysis. Raw pixel coordinates were extracted and transformed to a hip-centric reference frame by subtracting the median hip coordinates for each frame, allowing analysis of limb movement relative to the body independent of global animal displacement. Vertical coordinates were inverted such that higher values represent greater height from the walking surface. Gait cycles were identified based on periodic toe movement and normalised to 100% of the cycle to facilitate group comparisons. From these coordinates, several kinematic parameters were computed: knee and ankle joint angles (calculated from the vectors connecting adjacent landmarks), toe vertical excursion (maximum minus minimum y-coordinate during a stride), toe path length (cumulative Euclidean distance travelled by the toe across the gait cycle) and hip-rise amplitude (peak-to-trough vertical oscillation of the hip). For each animal, each metric was computed per gait cycle and averaged across cycles to yield a single value per animal, so that the unit of statistical analysis is the animal and not the individual video frame; this avoids pseudoreplication from treating non-independent within-animal frames as separate observations. Group comparisons across uninjured, injured (AAV-GFP), AAV-NR2F1 and AAV-NR2F6 animals were performed as described in the Statistical Analysis section. Per-animal kinematic metrics are provided in Supplementary Table S8.

### NR2F6 corepressor-domain deletion

Sequence alignment of NR2F1, NR2F2 and NR2F6 revealed that the repressor motif LSGYISLLL in NR2F1/2 aligns with a potential structurally analogous stretch LGIDNVCEL in NR2F6. To delete this domain, we employed a two-step, overlap- extension PCR strategy. Up- and downstream fragments flanking the motif were amplified with Phusion High-Fidelity DNA Polymerase [Thermo Scientific, #F530S] using primers bearing 18–20 bp overlaps (primer sequences in Supplementary Table S32). The first PCR comprised 7–8 cycles of 98 °C 2 min; 98 °C 15 s, 56–58 °C 30 s (primer-specific), 72 °C 3 min. The fragments were mixed, annealed via their overlaps and subjected to a second amplification (35 cycles: 98 °C 3 min; 95 °C 30 s, 69 °C 30 s, 72 °C 1.5 min; final 72 °C 5 min). Products were purified [QIAquick PCR Purification Kit, Qiagen, #28104] and assembled into the pAAV- CAG backbone using PCR-based enzyme cloning. Correct junctions and precise deletion were confirmed by Sanger sequencing.

### ATAC-seq data analysis

Paired-end ATAC-seq reads from ENCODE forebrain samples (E11, E12, E13, E14, E16, E18, P0 and adult; full accession list in the GitHub repository) were downloaded in FASTQ format. Adapter and low-quality bases were trimmed with fastp v0.23 (poly-G and poly-X filtering enabled; minimum read length 20 bp)(Chen, 2023). Cleaned reads were aligned to the mm10 reference genome with STAR v2.7.11a using the "--alignIntronMax 1 --alignEndsType EndToEnd" options to respect ATAC fragment structure(Dobin *et al*, 2013). Alignments with MAPQ < 10 were removed with SAMtools v1.17 and optical/PCR duplicates marked and dropped via Picard MarkDuplicates (Li *et al*, 2009). Library complexity, fragment-length periodicity, TSS enrichment and FRiP were evaluated with ATAQC (ENCODE pipeline) to confirm quality thresholds. Replicates passing QC were merged, and narrow peaks were called inside the TOBIAS-snake v0.14.0 workflow (MACS2 q ≤ 0.01, shifting +100/-50 bp). TOBIAS bias-correction, footprint scores and BINDetect motif scanning were run with default vertebrate parameters, producing per-stage BED, BigWig and motif-overlap files(Bentsen *et al*, 2020). Signal tracks were normalised to 1x genome coverage and exported as BigWigs for genome-browser visualisation(Raney *et al*, 2024). TF footprinting was performed genome-wide with the TOBIAS BINDetect module on the consensus ATAC-seq peak set across the three-stage developmental series. To focus on the pro-growth programme, footprint results were then intersected with the accessible regions associated with the 194 candidate genes, defined as ATAC-seq peaks overlapping the annotated gene body or lying within ±3 kb of the transcription start site (mm10 annotation). To define the developmental NR2F target atlas, we identified NR2F1/6 motif footprints that were present at high scores during early developmental stages and progressively lost into adulthood, using the TOBIAS binding-score output across the E11-to-adult series. Footprints recurring across multiple embryonic stages were collapsed to remove duplicate calls, yielding a single non-redundant gene list per factor. This in-house curated set constitutes the developmental NR2F1/6 target atlas referenced throughout the manuscript and is provided in Supplementary Table S19. All Snakemake rules, software versions, container recipes and command logs are hosted in our laboratory GitHub repository (https://github.com/VenkateshLab/nuclear-receptor-transcription-factor) for full reproducibility.

### Custom Reference Generation

To detect transgene-expressing nuclei, the AAV-GFP, AAV-NR2F1 and AAV-NR2F6 transgene sequences were appended as separate contigs to the mm10 reference FASTA, and matching transcript and exon entries were added to the GTF annotation. A custom reference was built with cellranger mkref (v7). Raw FASTQ reads were aligned to this augmented reference with cellranger count (v7), producing gene-barcode matrices and coordinate-sorted BAM files. To identify transgene-positive nuclei, BAM files were queried for reads overlapping each transgene locus; associated cellular barcodes were extracted and filtered to remove ambient RNA noise. These barcodes were added to the Seurat (v5.1) object metadata as a categorical label, enabling separation of transgene-positive and transgene-negative populations for UMAP visualisation and differential expression analysis. The custom annotation script is available at https://github.com/VenkateshLab/nuclear-receptor-transcription-factor.

### snRNA-seq processing and analysis

Raw BCL files were demultiplexed and FASTQ reads inspected with FastQC to check base quality and adapter content(Brown *et al*, 2017). Adapters and low- quality bases were removed with fastp v0.23(Chen *et al*, 2018), and clean reads were aligned to the mm10 reference using CellRanger v7.2 (default chemistry, 10x Genomics)(Satpathy *et al*, 2019). Each library yielded ∼3–4 k nuclei at ∼70 k reads per nucleus on an Illumina NovaSeq. Gene–barcode matrices were imported into Seurat v5 in R(Butler *et al*, 2018), filtered (nFeature > 200, < 6 % mitochondrial), log-normalised and integrated across samples with reciprocal PCA. Clustering (resolution 0.5) and UMAP visualisation yielded 13 transcriptional clusters (0-12). Transgene-expressing nuclei were identified using a custom annotation script that replaced the default Seurat automatic transgene-read assignment, which we found discarded a substantial number of genuinely transgene-positive nuclei owing to inconsistent calling; the custom script is deposited in our GitHub repository. Cell-type identity was assigned by evaluating the expression of canonical marker genes across each cluster (Supplementary Fig. S10, Supplementary Table S9): clusters expressing layer-V cortical projection-neuron markers (Satb2, Etv1, Crym, Ccnd2, Bcl11b) at high levels and lacking markers of non-cortical descending populations (paraventricular hypothalamus, red nucleus, locus coeruleus, raphe and hindbrain) were annotated as layer-V extratelencephalic projection neurons and retained for downstream analysis; small clusters expressing inhibitory-neuron (Slc32a1) or glial markers were classified as contaminants and excluded. After quality-control filtering and cluster assignment, 16,328 nuclei were retained (AAV-GFP: 5,403; AAV-NR2F1: 4,748; AAV-NR2F6: 6,177). Differential expression was then performed on the retained clusters with FindMarker (Wilcoxon rank-sum, min.pct = 0.25, log₂FC > 0.25). As a validation step, differential expression was independently repeated using the MAST framework with pseudobulk aggregation to account for the correlation structure among nuclei from a pooled sample; the two methods produced concordant gene sets with minimal differences (Supplementary Table S13). Custom figures such as UMAPs, heatmaps and volcano plots etc were generated with scCustomize(Marsh, 2021), and interactive exploration provided via ShinyCell(Ouyang *et al*, 2021); pre-built clusters and expression layers are browsable at https://nrtf.netlify.app/. All scripts, parameter files and processed matrices are deposited in our GitHub repository: https://github.com/VenkateshLab/nuclear-receptor-transcription-factor. All DEGs are summarized in Supplementary Table S33.

### CUT&RUN data processing

Adapter trimming and low-quality filtering were carried out with fastp v0.23(Chen *et al*, 2018). Clean reads were mapped to mm10 using BWA-MEM v0.7.17 (default settings), converted and sorted with SAMtools v1.17(Li *et al*, 2009), and PCR/optical duplicates were removed via Picard MarkDuplicates. Peaks were called against the no-antibody negative control with MACS2 v2.2.9.1 (default parameters), so every reported binding site is defined as enrichment over the matched background. Peak-to-gene annotation was performed with the annotatePeak function of ChIPseeker(Yu *et al*, 2015) in R against the mm10 reference, using TxDb.Mmusculus.UCSC.mm10.knownGene for transcript models. Each peak was assigned to the gene with the nearest transcription start site. Peaks within ±3 kb of a TSS were classified as promoter-associated; peaks outside this window were classified as distal (intronic or intergenic) and assigned to their nearest annotated gene by genomic distance.

### Multi-omic data integration

To link chromatin architecture, transcription-factor binding, transcriptional output and translational output at the gene level, we constructed a master integration table by mapping gene symbols across all four datasets (Hi-C, CUT&RUN, snRNA-seq and Ribo-seq) for NR2F6 using custom Python (v3.9) scripts. For Hi-C, A/B compartment status was determined by eigenvector decomposition, with positive PC1 values indicating active (A) and negative values indicating inactive (B) compartments; compartment switches were defined as transitions between conditions (B-to-A, A-to-B or stable). TAD boundaries were compared using reciprocal overlap to categorize domains as conserved, shifted or unique to a given condition. For CUT&RUN, genes were associated with a binding event if a peak fell within ±3 kb of the TSS (promoter) or within a distal regulatory element; binding status was recorded as a binary flag per factor, with fold-enrichment retained for quantitative analyses. For snRNA-seq and Ribo-seq, differential expression and differential translation were defined at p-adj ≤ 0.05 and | log2FC| ≥ 0.5. Translational efficiency (TE) was computed as Ribo-seq log2FC minus snRNA-seq log2FC, and genes were classified as concordant up (snRNA log2FC ≥ 0.5 and Ribo log2FC ≥ 0.5), concordant down (both ≤ -0.5), translationally attenuated (snRNA log2FC ≥ 0.5, TE ≤ -0.5) or translationally buffered (snRNA log2FC ≤ -0.5, TE ≥ 0.5). Genes were further classified by the intersection of binding and expression: bound and regulated (CUT&RUN peak present and DE threshold met), bound but not regulated (peak present, DE threshold not met) or regulated but not bound (DE threshold met, no detected peak). For the 194 pro-growth genes (Supplementary Table S24) and the developmental NR2F1/6 target atlas (Supplementary Table S19), each gene was additionally annotated for TAD status (conserved, shifted or NR2F6-specific), compartment switching (B-to-A, A-to-B or stable), CUT&RUN binding, snRNA-seq regulation, Ribo-seq regulation and translational-efficiency category, enabling a multi-layer view of how each gene is regulated across all four omic dimensions. All integration scripts are available in our GitHub repository. Global correlations between omics layers were assessed using Pearson’s r and Spearman’s rank correlation. Multi-omics heatmaps were generated using Seaborn(Waskom, 2021) and Matplotlib(Hunter, 2007) with rows normalized to show the percentage of genes within each regulatory category. Functional enrichment was performed using Gene Ontology (GO) analysis via custom scripts, with p-values adjusted for multiple testing using the Benjamini-Hochberg procedure.

### Ribosome-profiling data analysis

Raw FASTQ files were processed using the automated nf-core/riboseq pipeline (v1.2.0), running in Nextflow for quality control, adapter trimming, and sequence alignment. Briefly, low-quality bases and adapters were removed, and reads shorter than 20 nucleotides were discarded. Clean reads were depleted of contaminating ribosomal RNA (rRNA) and transfer RNA (tRNA) by mapping against dedicated reference databases. The remaining footprint reads were aligned to the mouse reference genome mm10 (GENCODE M25) using the STAR aligner (v2.7.10a) in two-pass mode, retaining only uniquely mapped alignments (--outFilterMultimapNmax 1, --alignEndsType EndToEnd). PCR duplicates were removed, and alignments with a MAPQ score < 30 were excluded. Ribosome occupancy footprints were quantified across annotated genomic regions (5Ö UTR, CDS, and 3Ö UTR). For downstream analysis of the unreplicated datasets (NR2F6, NR2F1, and Injured), raw footprint counts were imported into R. To account for sequencing depth variations across the single replicates, counts were normalized using the Trimmed Mean of M-values (TMM) method in edgeR (v3.38.4), and low-abundance genes with a Count Per Million (CPM) < 1 across all samples were filtered out.Differential translation changes were evaluated by directly calculating the log₂ fold-changes of TMM-normalized CPM values between conditions, rather than standard hypothesis testing.

To distinguish genuine translational regulation from passive transcriptional read-through, we computed translational efficiency (TE) for each gene as the difference between Ribo-seq log2 fold change and snRNA-seq log2 fold change (TE = Ribo-seq log2FC minus snRNA-seq log2FC), restricted to genes detected in both datasets. A log2FC threshold of ±0.5 was applied uniformly to both the snRNA-seq and Ribo-seq measurements, corresponding to an approximately 1.4-fold change and chosen to capture biologically meaningful shifts while excluding noise from near-zero changes. Genes were then classified into four mutually exclusive categories: concordant up (snRNA-seq log2FC ≥ 0.5 and Ribo-seq log2FC ≥ 0.5), in which both transcription and ribosome occupancy increase together; concordant down (snRNA-seq log2FC ≤ -0.5 and Ribo-seq log2FC ≤ -0.5), in which both decrease; translationally attenuated (snRNA-seq log2FC ≥ 0.5 but TE ≤ -0.5), in which mRNA rises but translational efficiency falls; and translationally buffered (snRNA-seq log2FC ≤ -0.5 but TE ≥ 0.5), in which mRNA abundance falls but ribosome occupancy is maintained or elevated, indicating active post-transcriptional preservation of protein output. This classification was performed independently for NR2F1 and NR2F6 conditions. GO enrichment of each category was performed with DAVID (Biological Process, EASE p ≤ 0.05), consistent with the enrichment framework used for Fig. 1a. Full per-gene TE values and category assignments are provided in Supplementary Table S17.

### In-situ Hi-C

C57BL/6 mice received bilateral cortical injections of AAV9-NR2F6 or AAV9- GFP (control) and a T8 crush 7 d later. Motor cortices (35–40 mg; pooled from 3–5 mice per group) were collected 7 d post-injury, flash-frozen, pulverised in liquid N₂, and processed according to the Ren-lab/ENCODE in-situ Hi-C protocol with tissue-specific optimisation(The ENCODE Project Consortium *et al*, 2020; Dixon *et al*, 2012). Chromatin was cross-linked in 1 % formaldehyde [Thermofisher, #12755] for 10 min (RT), quenched with 0.2 M glycine (MP 808822, 5 min), and nuclei were isolated in ice-cold Hi-C lysis buffer (5 mM CaCl₂, 3 mM MgAc₂, 2 mM EDTA, 0.5 mM EGTA, 10 mM Tris-HCl pH 8.0, 1 mM DTT, 0.1 mM PMSF + protease inhibitors, MCE HY-K0010). After 1 M sucrose gradient purification, nuclei were permeabilised in 0.5 % SDS (10 min, 62 °C), SDS was quenched with 10 % Triton X-100, and chromatin was digested overnight with 100 U MboI (NEB, #R0147, 37°C). MboI was heat-inactivated (62 °C, 20 min); 5Ö ends were filled in with biotin-14-dATP (0.4 mM; Thermofisher #19524-016) plus dCTP/dGTP/dTTP (10 mM each; Thermofisher #18254011/013/018) and Klenow (NEB, #M0210, 37 °C, 45 min). Blunt ends were proximity- ligated for 4 h at RT with T4 DNA ligase [NEB, #M0202] in ligation buffer containing 10 % Triton X-100 and 20 mg mL⁻¹ BSA [Sigma, #A7906]. Cross-links were reversed overnight with Proteinase K [Sigma, #P2308, 65 °C]. DNA was purified, sheared to 300–500 bp on a Covaris M220 (Peak 50, Duty 10, Cycles 350, 70 s) and size-selected with AMPure XP beads [Beckman, #A63881]. Biotinylated fragments were captured on Dynabeads MyOne Streptavidin T1 (Invitrogen 65601), end-repaired, dA-tailed, and ligated to Illumina TruSeq UD adapters (20040870). Libraries were PCR-amplified [Illumina, #20015964], bead-purified, and sequenced to ∼5 × 10⁸ paired-end reads on an Illumina NovaSeq.

### HiC data analyses

Paired-end Hi-C reads were first inspected with FastQC v0.11.9 to verify base quality(Brown *et al*, 2017), GC bias and adapter carry-over, then enzyme-specific 3Ö overhangs were trimmed with homerTools v4.11 (-mis 0 -matchStart 20 -min 20) to remove MboI/DpnII fragments (GATC)(Dixon *et al*, 2012). Because HOMER recommends single-end processing, R1 and R2 were aligned independently to the *mm10* reference with Bowtie2 v2.5.1 in unpaired mode (-U) using 30 parallel threads (-p 30)(Heinz *et al*, 2010). The resulting SAM files were collapsed into tag directories with makeTagDirectory (-tbp 1 to discard PCR duplicates; QC flags -checkGC -restrictionSite GATC -genome mm10) and converted to .hic format for interactive viewing in Juicebox via tagDir2hicFile.pl(Durand *et al*, 2016). Principal- component analysis of contact matrices was carried out with runHiCpca.pl at 25-kb resolution with a 50-kb smoothing window, producing PC1 bedGraphs that delineate open (A) and closed (B) compartments; replicate-aware comparisons between control and NR2F6 samples were performed with annotatePeaks.pl followed by getDiffExpression.pl (edgeR backend) to flag differential compartment shifts (Robinson *et al*, 2010). Chromatin compaction was quantified with analyzeHiC -compactionStats at 5-kb resolution/15-kb window, yielding distal-to-local interaction ratio (DLR) and inter-chromosomal interaction fraction (ICF) tracks; differential compaction profiles were obtained with subtractBedGraphs.pl. Topologically associating domains (TADs) and chromatin loops were identified using findTADsAndLoops.pl (3-kb bins, 15-kb sliding window), producing .tad.2D.bed and .loop.2D.bed files that were merged across conditions with merge2Dbed.pl, rescored with findTADsAndLoops.pl score, and subjected to edgeR-based differential testing via getDiffExpression.pl to catalogue NR2F6-specific gains or losses in TADs and loops. All command logs, Snakemake workflows and parameter files are archived at https://github.com/VenkateshLab/nuclear-receptor-transcription-factor.

### Data availability

All primary sequencing data generated in this study—single-nucleus RNA-seq, CUT&RUN, Ribo-seq and in-situ Hi-C have been deposited in the NCBI Gene Expression Omnibus under the superseries accession PRJNA1217058 and will be released upon publication. Public developmental ATAC-seq datasets re-analysed here were retrieved from ENCODE; the full accession list is provided in our GitHub repository (https://github.com/VenkateshLab/nuclear-receptor-transcription-factor). An interactive browser for the snRNA-seq data is available at https://nrtf.netlify.app/. All analysis pipelines, Snakemake workflows and custom scripts are openly accessible in the same GitHub repository.

### Code availability

All custom code used for data processing, statistical analyses, figure generation and interactive visualisation—including Snakemake workflows, Docker/Singularity container recipes, R and Python notebooks, and DeepLabCut post-processing scripts—is openly available at our laboratory GitHub repository: https://github.com/VenkateshLab/nuclear-receptor-transcription-factor. The repository contains detailed README files, environment.yml/requirements.txt specifications, and step-by-step instructions that enable full reproduction of the results presented in this study.

### Statistical Analyses

The choice of statistical test for each quantified panel was determined by the number of groups, the experimental design and the distribution of the data, and is detailed in full in Supplementary Table S30. Normality was assessed by the Shapiro-Wilk test and variance homogeneity by the Brown-Forsythe test. For multi-group comparisons of non-normally distributed data (neurite-length measurements, Figs. 2c, 2f, 7d; thoracic-crush fiber indices, Fig. 4d; grip strength, Fig. 5c), the Kruskal-Wallis test was used followed by Dunn’s multiple-comparisons test with adjusted p-values. For multi-group comparisons where data met normality and equal-variance assumptions (pyramidotomy fiber indices, Fig. 3e), ordinary one-way ANOVA was used followed by Dunnett’s post-hoc test against the AAV-GFP control, with variance homogeneity confirmed by the Brown-Forsythe test. For hip-rise recovery across multiple timepoints (Fig. 5c), repeated-measures two-way ANOVA with time as the repeated factor was used, followed by Sidak multiple-comparisons correction; residual normality was confirmed by quantile-quantile plots. For the bioinformatic analyses, differential expression in the snRNA-seq data was assessed with the Wilcoxon rank-sum test (FindMarkers, Seurat) using thresholds of p-adj ≤ 0.05 and | log2FC| ≥ 0.5; differential translation in the Ribo-seq data was assessed with the Wilcoxon rank-sum test with the same significance threshold. CUT&RUN peaks were called against the no-antibody negative control with MACS2 (default parameters). Gene Ontology enrichment was performed with DAVID (Biological Process domain, EASE p ≤ 0.05). Pearson and Spearman correlations were used to assess the relationship between CUT&RUN promoter occupancy and snRNA-seq expression change. All data are presented as mean ± SEM unless otherwise stated; for non-parametric analyses, data are presented as median ± interquartile range. Individual data points are shown on all bar graphs. A complete summary of all statistical tests, sample sizes, normality assessments, variance tests and exact p-values for every quantified panel is provided in Supplementary Table S30.

## Author contributions

Ishwariya Venkatesh (I. V.) and Yogesh Sahu (Y.S.) conceived and designed the study. Y.S., M. Kumaran, S. Banerjee, A. Narayan PS, K. Sanyal, M. Konda, S. Soni, D. Kumar, A.S. Menon, P. Chermakani, A. Sukumar, F. Farooq, S. Manjunath, D.S. Beji and A. Biswas performed the experiments, provided technical expertise, and contributed to data analysis and interpretation. Manuscript drafting was led by I. V., with input from Y.S. and M. Kumaran. Figure preparation was led by Y.S. and M. Kumaran. All authors reviewed and approved the final manuscript. * M. Kumaran, S. Banerjee and A. Narayan PS contributed equally. # K. Sanyal, M. Konda, S. Soni, D. Kumar, A.S. Menon and P. Chermakani contributed equally.

## Acknowledgments

We gratefully acknowledge financial support from the Council of Scientific and Industrial Research (CSIR – HCP, FBR), CSIR-Centre for Cellular and Molecular Biology (CSIR-CCMB), the Science and Engineering Research Board (SERB, SRG), the Department of Biotechnology (DBT, Biomedical Sciences), the Anusandhan National Research Foundation (ANRF-ARG) and the BFIBiome programme, all awarded to I. V. We thank the CCMB core facilities and their staff for invaluable assistance: Zareena (Tissue-Culture), Sairam and Prashanth (Animal House), Srinivas (FACS), Subhajit Sen (Genomics), Dr. Nitla Mahesh and Suman Bhandari (Microscopy) and FBC. We acknowledge the labs of Dr. Vinay Nandicoori and Dr. Sriram Varahan for their support. We are grateful to our lab manager Dhruva Kesireddy, grants manager Dr.Shringika Soni, and LabTech support Mr. Venkateshwarulu for dedicated laboratory support that kept day-to-day operations running smoothly.

## Supplementrary Figure

**Supplementary Figure S1:** Workflow and Developmental Expression Profiles of NR2F1 and NR2F6 in the Murine Forebrain.

**Supplementary Figure S2:** SDS-PAGE analysis of AAV9 viral capsid proteins from GFP, NR2F1, and NR2F6 preparations.

**Supplementary Figure S3 :** In-vitro screening identifies NR2F6 and NR2F1 as potentially significant pro-growth transcription factors

**Supplementary Figure S4:** AAV injection-site validation in motor cortex.

**Supplementary Figure S5:** High-magnification midline-crossing analysis after pyramidotomy

**Supplementary Figure S6:** GFAP staining confirms injury-site boundaries after thoracic crush injury.

**Supplementary Figure S7:** Transgene expression validation for NR2F1 and NR2F6.

**Supplementary Figure S8:** Gait analysis of uninjured, injured, and treated mice.

**Supplementary Figure S9 :** Experimental Timeline, Kinematic Gait Analysis, and Spatiotemporal Gait Parameters Following Spinal Cord Injury (SCI).

**Supplementary Figure S10:** Single-nucleus RNA-seq clustering and cell-type annotation of AAV-transduced neurons.

**Supplementary Figure S11:** Validation of ribosome profiling workflow and 30 bp ribosome-protected fragment (RPF) enrichment from sorted GFP-positive cells.

**Supplementary Figure S12:** Translational buffering of gene expression following NR2F1 and NR2F6 gene treatment.

**Supplementary Figure S13:** Integrative CUT&Run and snRNA-seq analysis reveals the direct transcriptional targets and genomic distribution of NR2F1 and NR2F6.

**Supplementary Figure S14:** Integrated multi-omics analysis of progrowth gene regulation by NR2F1 and NR2F6.

## Supplementary Table

**Supplementary Table S1:** GO enrichment of the 194 developmentally downregulated pro-growth genes

**Supplementary Table S2:** List of the 194 candidate pro-growth genes

**Supplementary Table S3:** NR2F1 and NR2F6 footprinting across the 194 pro-growth loci

**Supplementary Table S4:** In vitro neurite-growth quantification and statistics

**Supplementary Table S5:** Pyramidotomy CST sprouting quantification

**Supplementary Table S6:** Thoracic crush CST regeneration quantification

**Supplementary Table S7:** Behavioural recovery quantification

**Supplementary Table S8:** DLC-derived gait metrics and Statistical analysis

**Supplementary Table S9:** snRNA-seq cluster annotation and marker genes

**Supplementary Table S10:** snRNA-seq DEG results for NR2F6 versus GFP/injured control

**Supplementary Table S11:** snRNA-seq DEG results for NR2F1 versus GFP/injured control

**Supplementary Table S12:** Pseudobulk snRNA-seq differential-expression validation

**Supplementary Table S13:** Direct NR2F1 versus NR2F6 transcriptional comparison

**Supplementary Table S14:** NR2F6 corepressor-domain mutant neurite-growth statistics

**Supplementary Table S15:** NR2F6 Ribo-seq differential-translation results

**Supplementary Table S16:** NR2F1 Ribo-seq differential-translation results

**Supplementary Table S17:** Translational-efficiency and buffering analysis

**Supplementary Table S18:** NR2F6 and NR2F1 CUT&RUN peak annotation

**Supplementary Table S19:** Developmental target of NR2F6 and F1

**Supplementary Table S20:** Overlap between developmental NR2F6 and F1 targets and post-injury CUT&RUN peaks

**Supplementary Table S21:** NR2F6-bound and transcriptionally regulated genes

**Supplementary Table S22:** NR2F1-bound and transcriptionally regulated genes

**Supplementary Table S23:** Promoter occupancy versus gene-expression correlation data

**Supplementary Table S24:** Pro-growth 194-gene multi-omic categorisation

**Supplementary Table S25:** NR2F6-specific and shifted TADs

**Supplementary Table S26:** Genes within NR2F6-specific or shifted TADs linked to expression/Ribo-seq changes

**Supplementary Table S27:** A/B compartment switching after NR2F6 overexpression

**Supplementary Table S28:** GO enrichment of genes in NR2F6-remodelled TADs and switched compartments

**Supplementary Table S29:** Translation-regulatory hub genes

**Supplementary Table S30:** Statistical-analysis summary for all quantified panels

**Supplementary Table S31:** AAV-titter information

**Supplementary Table S32:** Cloning primers used

**Supplementary Table S33:** snRNA overexpression of NR2F1 and NR2F6 file

