## Supplementray_Figure for "Nuclear Receptor Transcription Factors promote axon regeneration in the Adult Corticospinal Tract"

a

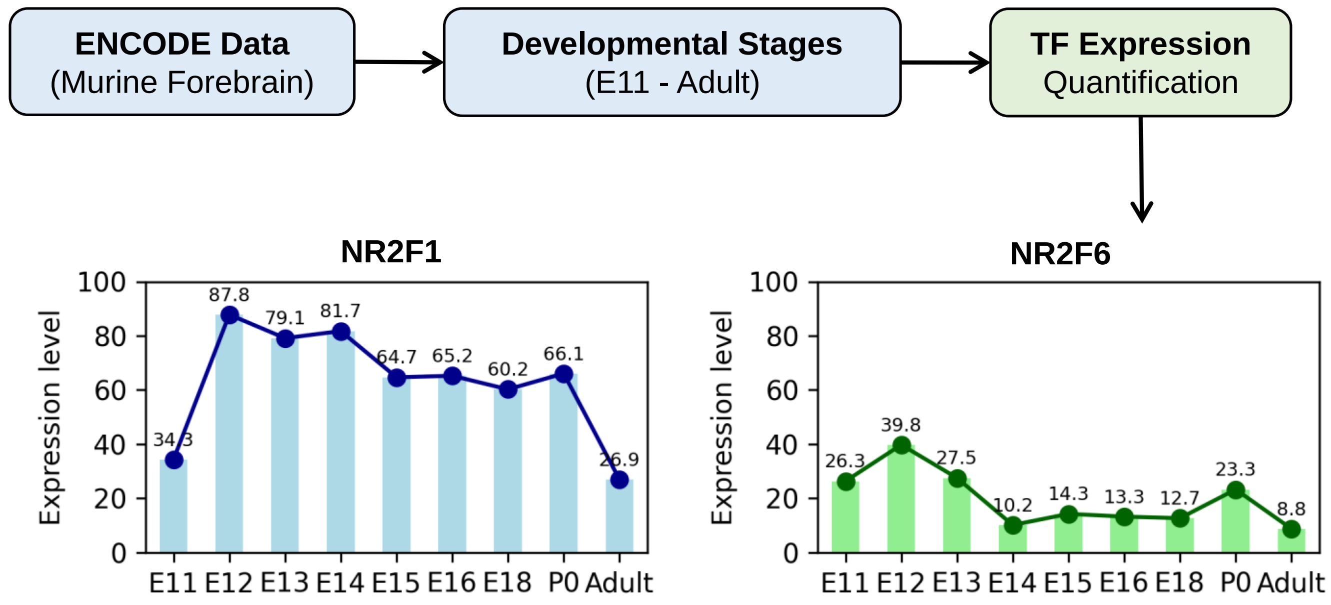


**Supplementary Figure S1: Workflow and Developmental Expression Profiles of NR2F1 and NR2F6 in the Murine Forebrain**
Schematic workflow and temporal expression analysis of target transcription factors. The top flowchart outlines the data mining pipeline, where public RNA-seq data from the murine forebrain was retrieved from the ENCODE database across a developmental continuum from embryonic day 11 (E11) to adulthood to quantify transcription factor (TF) expression levels. The bottom panels display the resulting quantified expression profiles for NR2F1 (left, blue) and NR2F6 (right, green) across sequential developmental time points (E11, E12, E13, E14, E15, E16, E18, P0, and Adult). The x-axis tracks the developmental stages, and the y-axis denotes the normalized expression levels. Numerical values labeled above each data point indicate the precise expression value at that specific stage, highlighting the distinct temporal regulation of each TF during forebrain development.

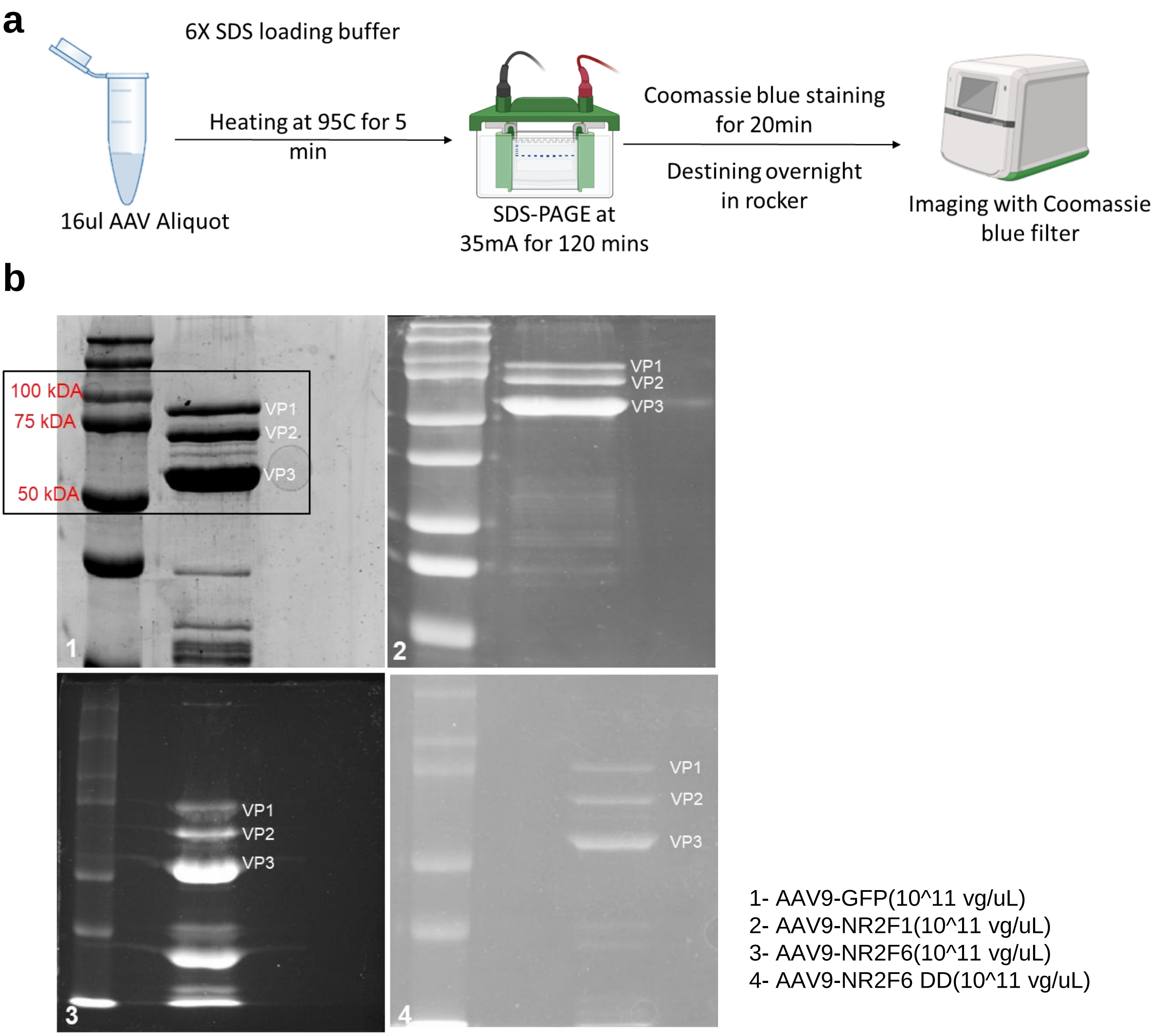


**Supplementary Figure S2: SDS-PAGE analysis of AAV9 viral capsid proteins from GFP, NR2F1, and NR2F6 preparations.**
(a) Workflow schematic for Coomassie-stained SDS-PAGE. AAV9 viral aliquots (16 µL) were mixed with 6X SDS loading buffer, denatured at 95°C for 5 minutes, run on SDS-PAGE at 35 mA for 120 minutes, stained with Coomassie blue for 20 minutes, and destained overnight before imaging.(b) Representative gels showing the capsid proteins VP1, VP2, and VP3 for (1) AAV9-GFP , (2) AAV9-NR2F1 ,(3) AAV9-NR2F6 (4) AAV9- NR2F6 DD . The presence of all three capsid proteins confirms successful packaging and purity of the AAV9 viral preparations. Approximate molecular weights are indicated by the protein ladder on the left.

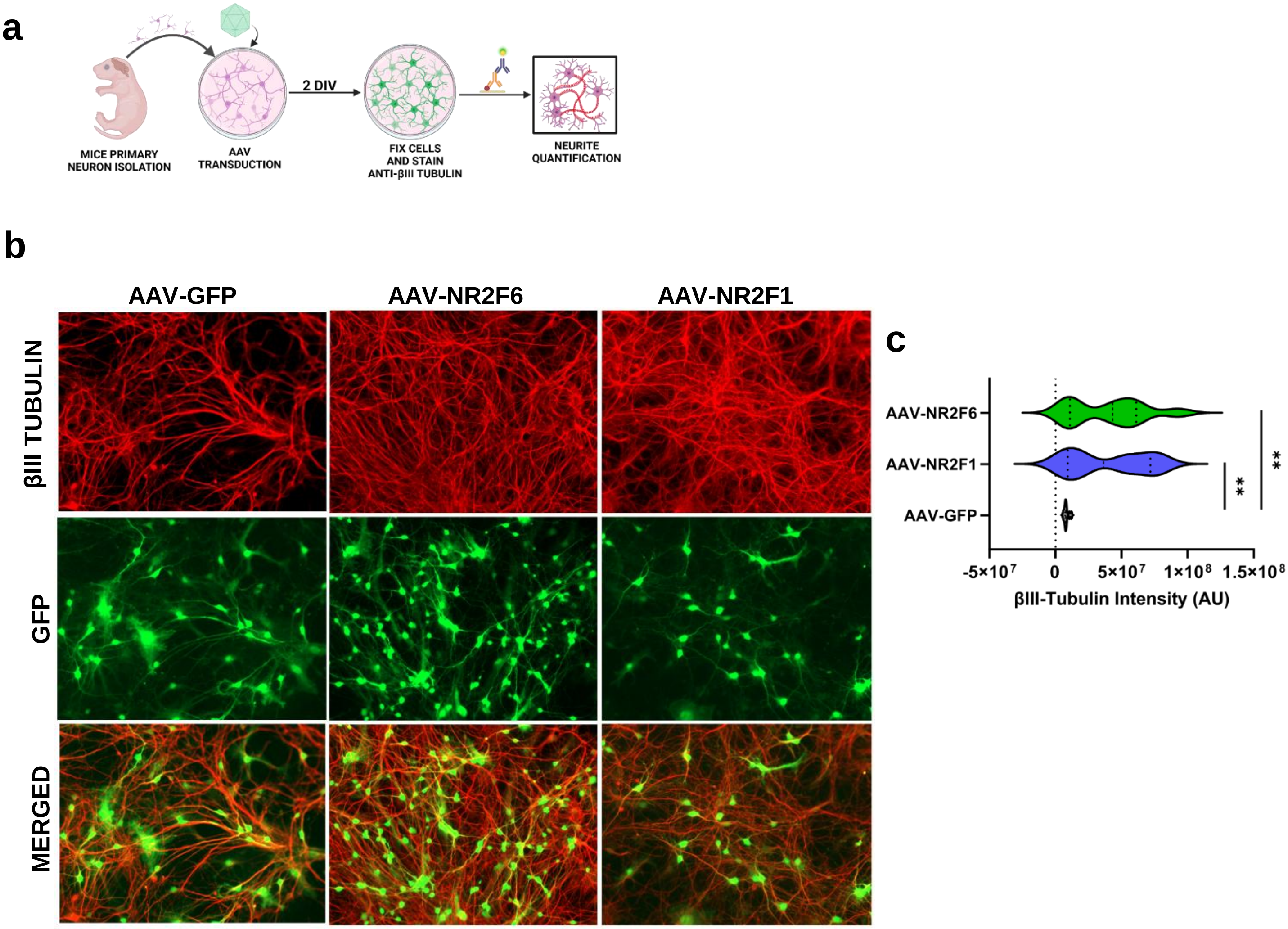


**Supplementary Figure S3 :** ***In vitro*** **screening identifies NR2F6 and NR2F1 as potentially significant pro-growth transcription factors**

(a) Schematic of the experimental timeline for mouse primary neuron culture. Mouse primary cortical neurons were isolated and transduced with AAV-GFP (control), AAV-NR2F1, or AAV-NR2F6. At 2 DIV, cells were fixed and stained for βIII Tubulin to quantify the extent of neurite outgrowth. (b) Representative immunofluorescence images of primary neurons transduced with AAV-GFP (control), AAV-NR2F6, or AAV-NR2F1. Cells were stained for βIII Tubulin(red) and GFP was visualized in green. Scale bar = 50 µm. (c) Quantification of βIII Tubulin intensity was quantified using ImageJ software. Around 1800 to 2000 neurons were counted for their βIII Tubulin intensity per treatment group. Statistical analysis was performed using one-way ANOVA followed by unpaired t-test . Data represent mean ± SEM ($p = 0.0089$, **$p <= 0.01$) ($n = 1$).

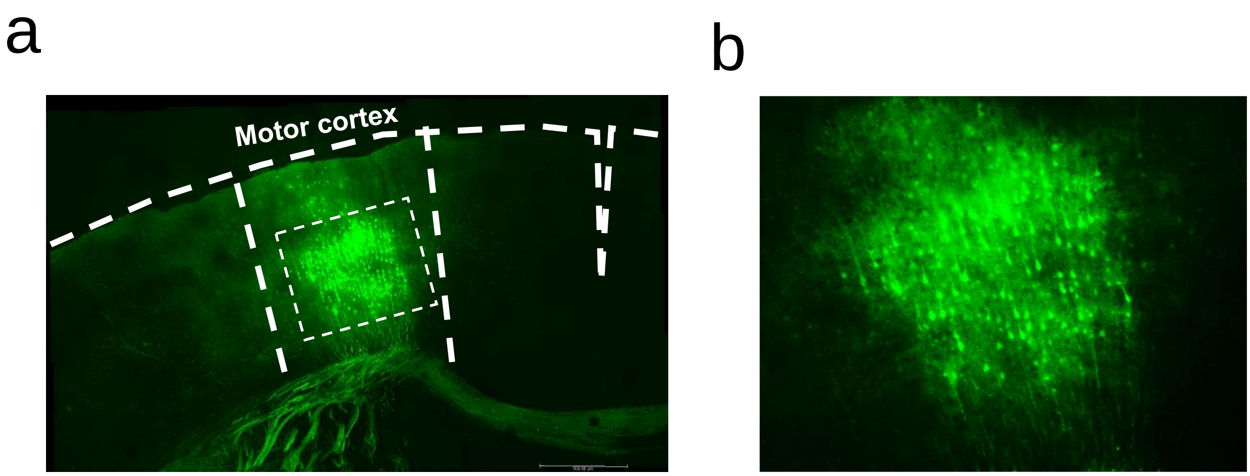


**Supplementary Figure S4: AAV injection-site validation in motor cortex**
(a) Low-magnification fluorescent microscopy image of the motor cortex (M1) showing labelled neurons. Dashed white lines demarcate the boundaries of the motor cortex and the area of interest. Scale bar at the bottom left of the image represents 500 µm. The inset dashed box zooms into the region shown in panel (b) High-magnification view of the boxed region in panel (a) showing the detailed neuron distribution within the motor cortex. Fluorescent labelling highlights the neurons involved in motor control, showing their morphology and arrangement in the cortical layers. A detailed information on co-ordinates are provided in methods section.

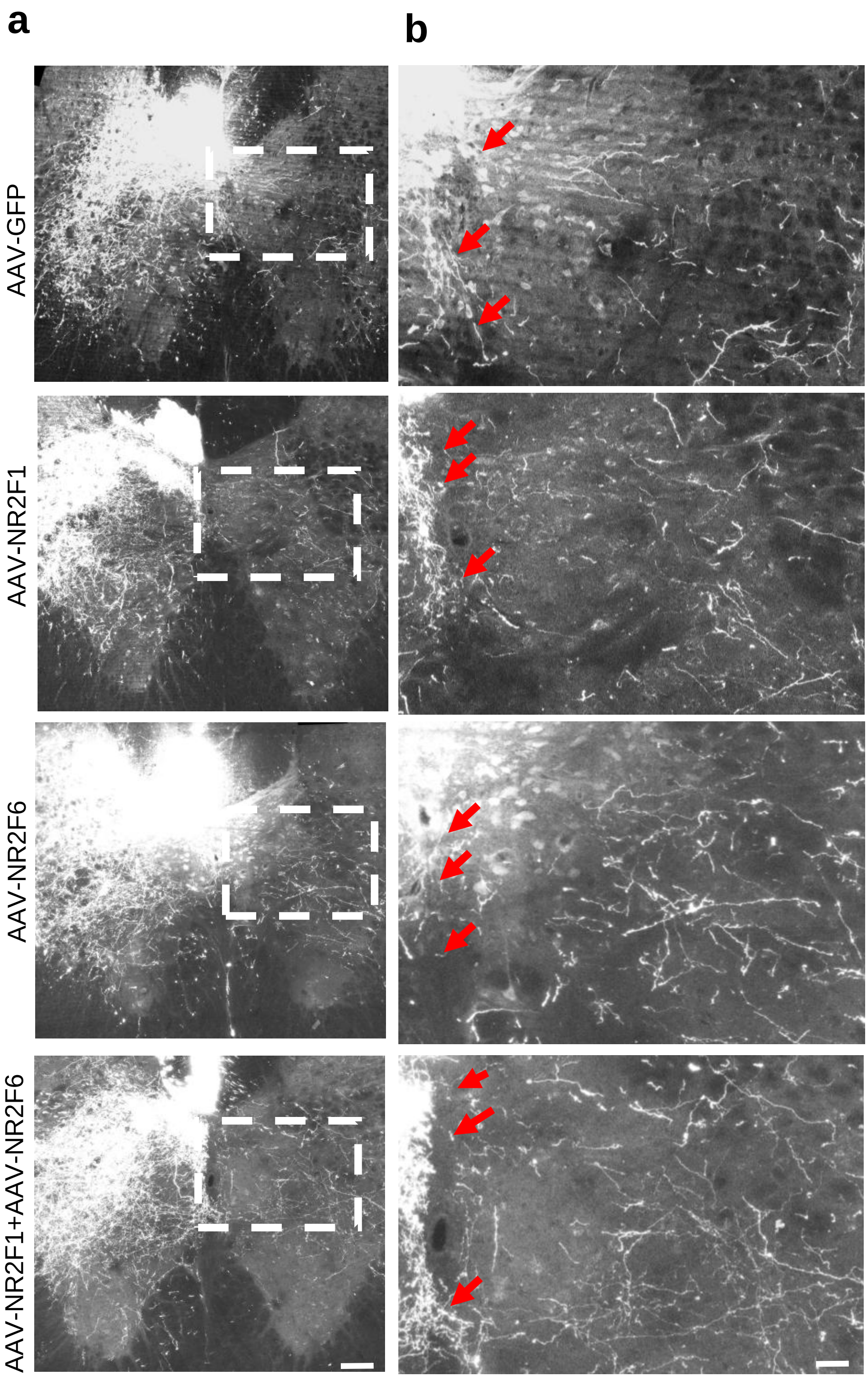


**Supplementary Figure S5: High-magnification midline-crossing analysis after pyramidotomy**
(a) Representative images of cervical spinal cord sections. Dotted boxes indicate regions of interest (ROI) for midline crossing. (b) High-magnification crops of the ROIs; red arrows indicate axons crossing the midline into the denervated side. Scale bars = 200 µm (overviews) and 50 µm (zooms).

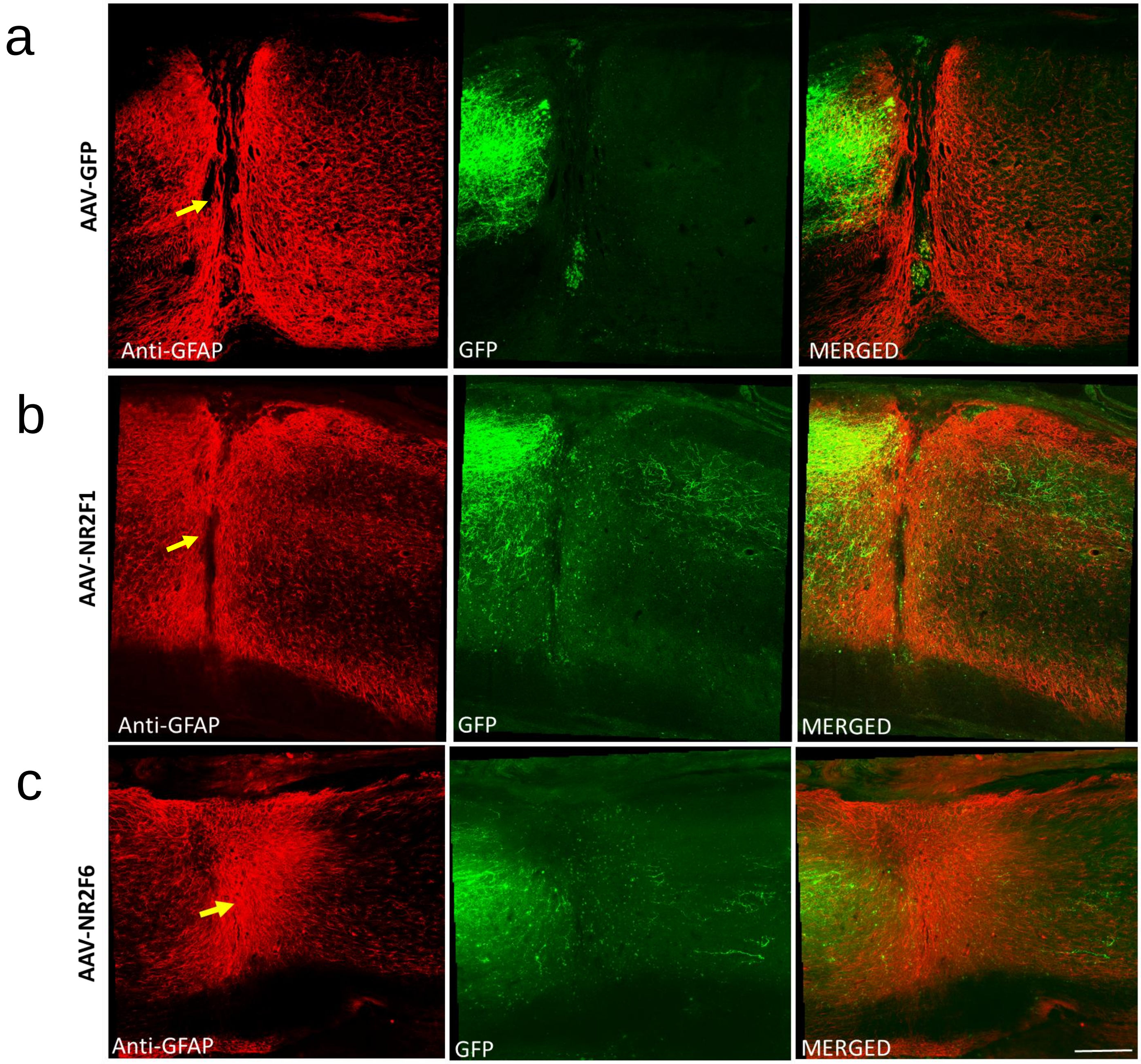


**Supplementary Figure S6: GFAP staining confirms injury-site boundaries after thoracic crush injury**
(a-c) representative images of sagittal sections of spinal cord (AAV-GFP, AAV-NR2F1, AAV-NR2F6) near injury site was confirmed with GFAP immunostaining. Injury site is shown with arrow head (yellow). "GFAP, red; GFP-labelled axons, green.Scale bar= 200μm

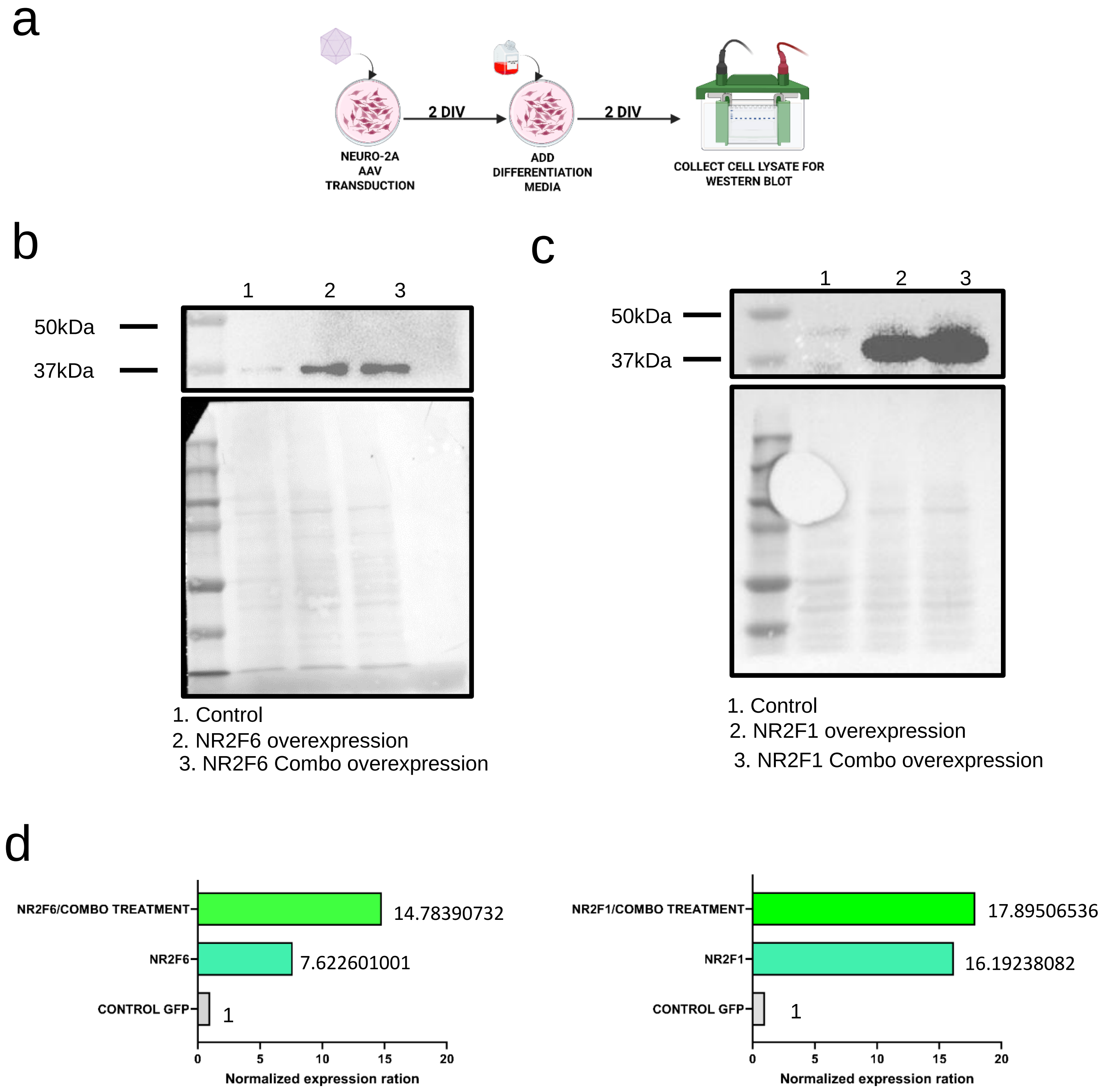


**Supplementary Figure S7: Transgene expression validation for NR2F1 and NR2F6**
(a) Schematic illustrating the experimental timeline for Neuro-2A cell transduction and differentiation. Cells were transduced with AAV, followed by the addition of differentiation media for 2 days in vitro (DIV). Cell lysates were collected for Western blot analysis. b Western blot showing NR2F6 expression in the three conditions: control, NR2F6 overexpression, and NR2F6+NR2F1 combined overexpression. The membrane was probed with an anti-NR2F6 antibody.
c Western blot showing NR2F1 expression under the same experimental conditions: control, NR2F1 overexpression, and NR2F1+NR2F6 combined overexpression. The membrane was probed with an anti-NR2F1 antibody.
(d) Bar plots displaying the normalized expression levels of NR2F6 (left) and NR2F1 (right) across the conditions. The expression levels are shown relative to the control GFP condition. Data are presented as the mean normalized expression ratio, with each bar representing a single replicate. The numeric values above the bars indicate the specific expression ratios for each condition. Detailed procedures are provided in the methods.

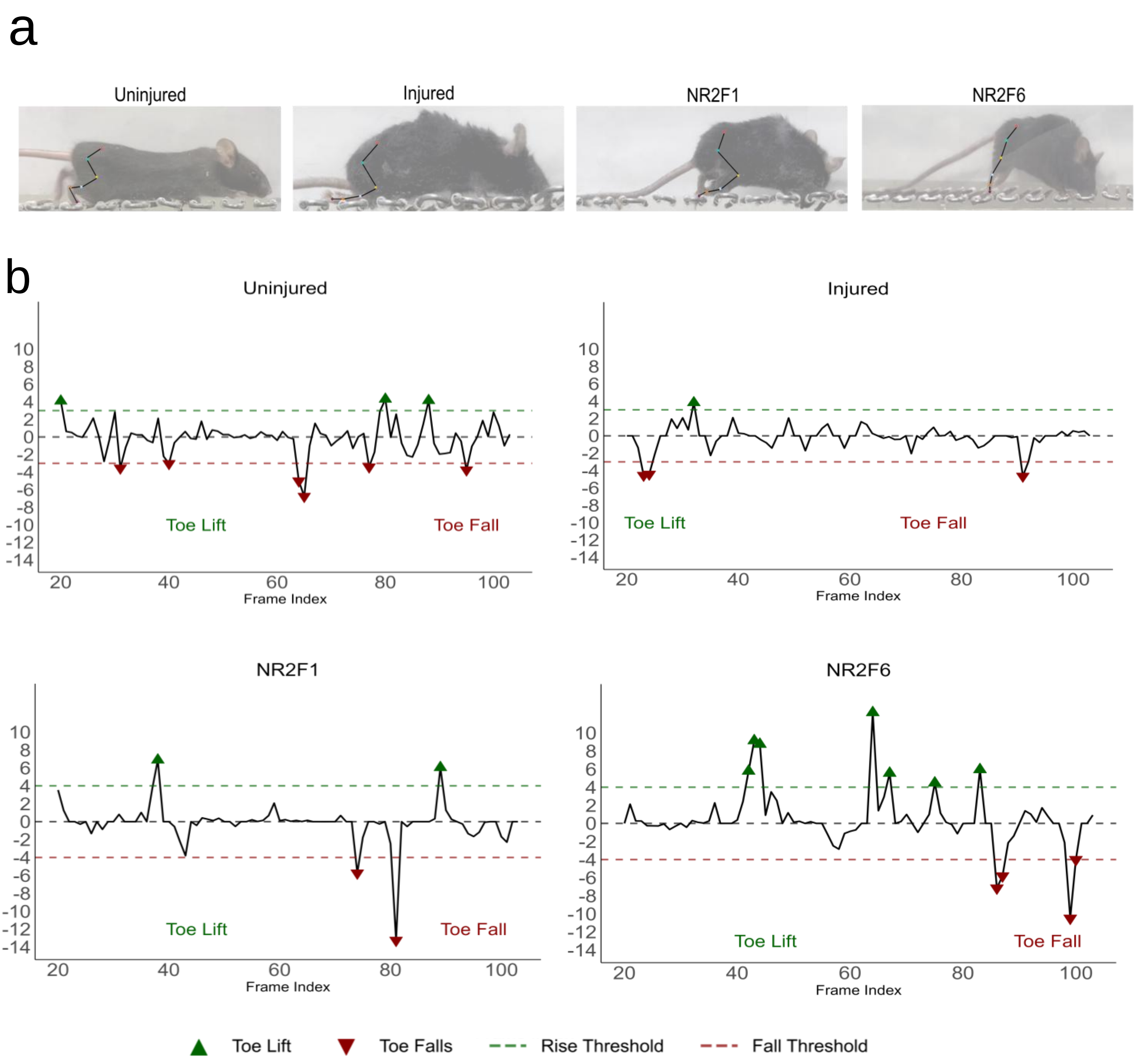


**Supplementary Figure S8: Gait analysis of uninjured, injured, and treated mice.** (a) Stick plot representations showing gait patterns during ladder rung experiment. Representative traces demonstrate normal gait in uninjured mice, impaired gait with toe dragging in injured mice, and improved gait with elevated hip positioning in mice treated with NR2F1 or NR2F6. (b) Quantitative analysis of toe lift and toe fall during ladder rung experiment. Dot plots show toe lift (green upward triangles) and toe fall (red downward triangles) measurements relative to the rung surface (0 line, solid horizontal line). Dashed horizontal lines indicate analysis thresholds: +0.5 cm for toe lift (rise threshold) and -0.5 cm for toe fall (fall threshold). Data are shown for uninjured, injured, NR2F1-treated, and NR2F6-treated groups across frame indices representing the gait cycle. Frame index represents temporal progression during the walking sequence.

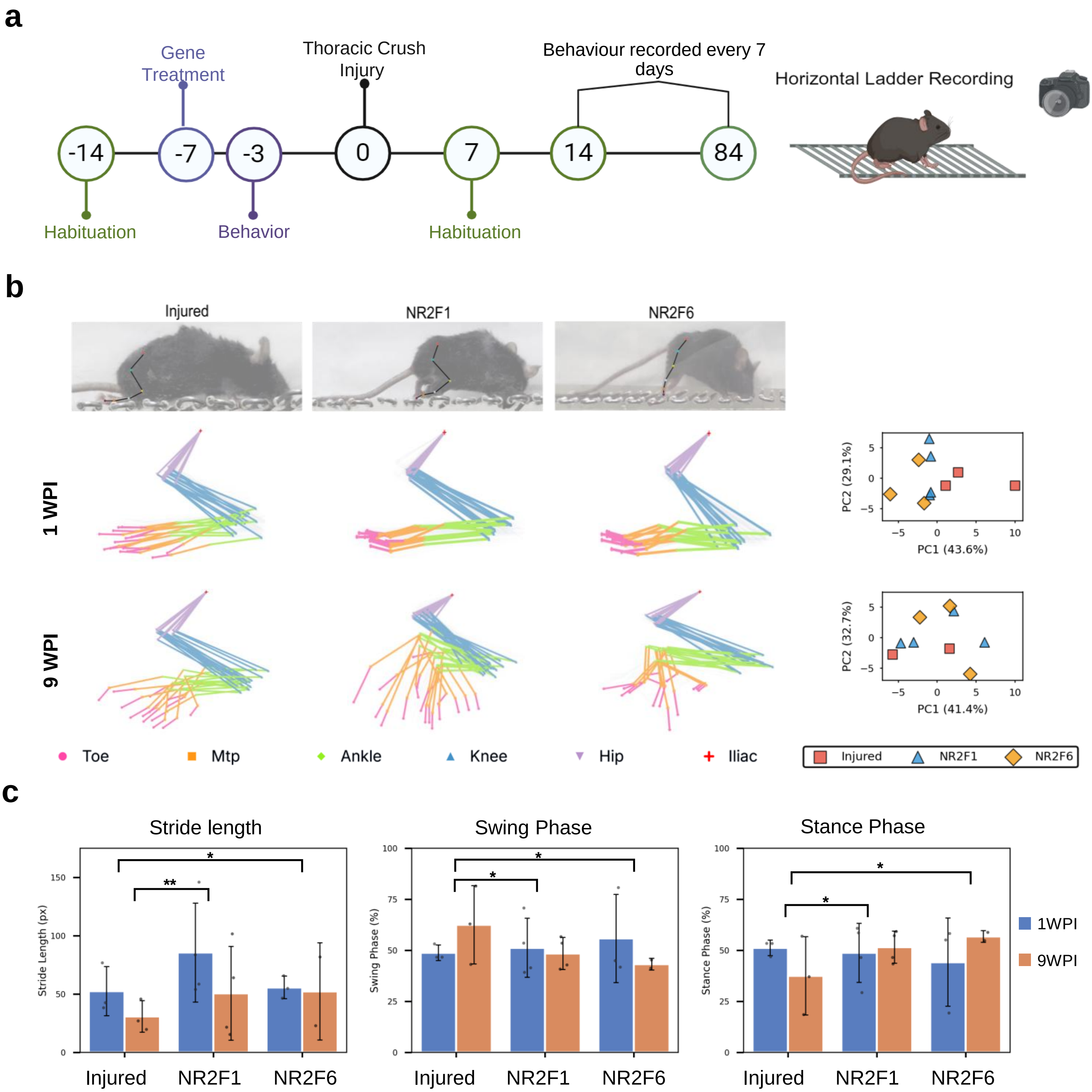


**Supplementary Figure S9 : Experimental Timeline, Kinematic Gait Analysis, and Spatiotemporal Gait Parameters Following Spinal Cord Injury (SCI)**
(a) Experimental Timeline and Behavioral Setup. Schematic representation of the experimental timeline for the thoracic crush injury model, gene treatment intervention, and subsequent behavioral assessments. Mice underwent habituation and baseline behavioral tracking prior to a thoracic crush injury (Day 0). Post-injury behavioral recovery was recorded every 7 days up to 84 days using a horizontal ladder rung walking test setup.
(b) Kinematic Joint Coordination and Principal Component Analysis (PCA). Representative stick plots and corresponding PCA plots characterizing hindlimb gait patterns across three experimental conditions: Injured (control), NR2F1-treated, and NR2F6-treated mice at 1 week post-injury (1 WPI) and 9 weeks post-injury (9 WPI). Stick plots illustrate the spatial trajectories and coordination of key anatomical landmarks during the stance and swing phases of locomotion. Precise joint position extraction and automated marker tracking were performed using DeepLabCut (DLC). The tracked anatomical landmarks include the Toe (pink circles), Metatarsophalangeal (Mtp) joint (orange squares), Ankle (green diamonds), Knee (blue upward-pointing triangles), Hip (purple downward-pointing triangles), and Iliac crest (red crosses). Accompanying PCA plots (right) demonstrate the clustering and segregation of gait profiles for each group based on kinematic variance along Principal Component 1 (PC1) and Principal Component 2 (PC2) at 1 WPI and 6 WPI.
(c) Quantitative Spatiotemporal Gait Parameters. Bar graphs quantifying discrete gait metrics across the Injured, NR2F1, and NR2F6 experimental groups at 1 WPI (blue bars) and 6 WPI (orange bars). Quantified parameters include Stride Length (pixels), Swing Phase duration (%), and Stance Phase duration (%). Data are presented as mean standard deviation (SD), with individual data points overlaid to show sample distribution.

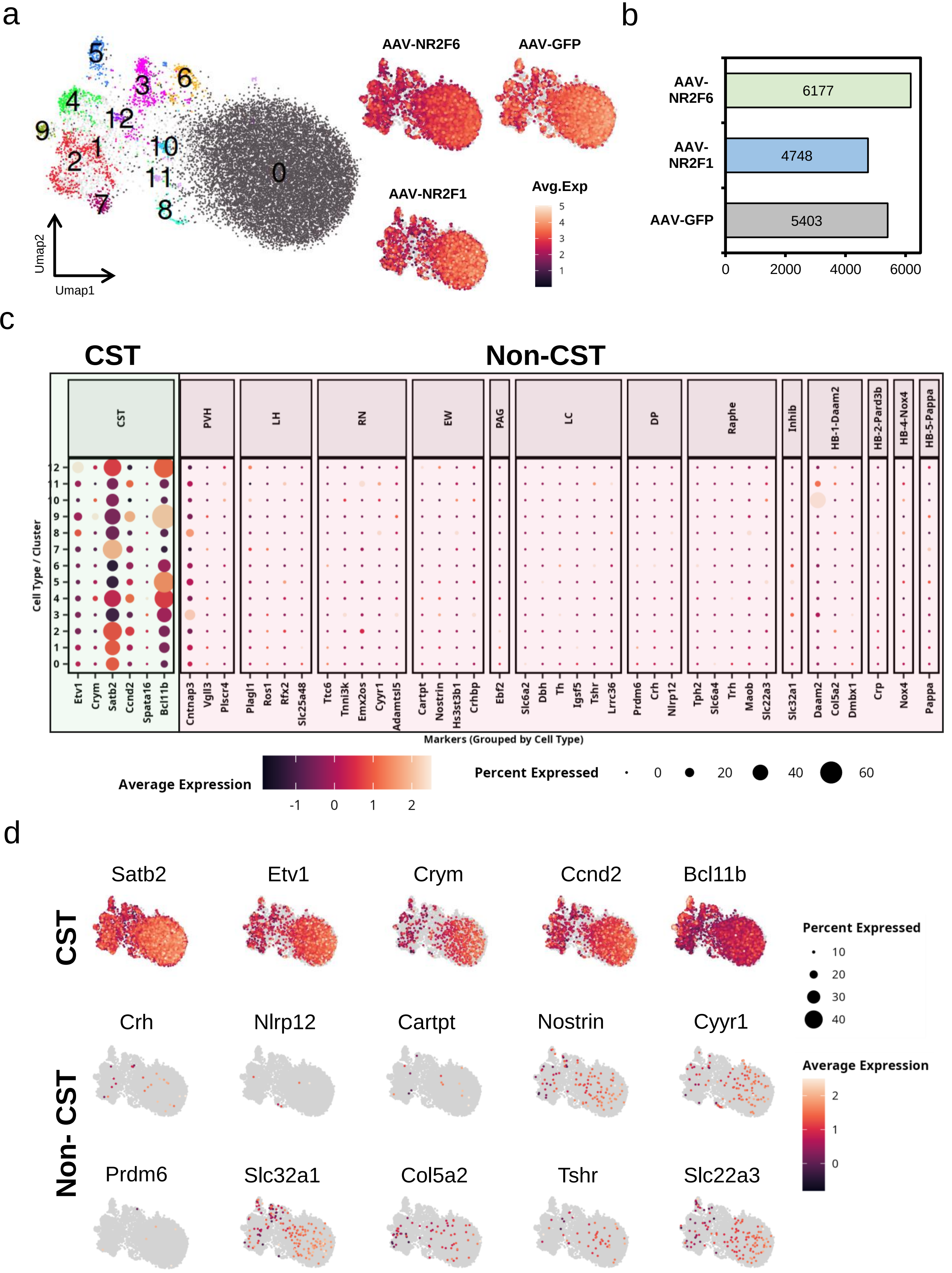
a
b
AAV-NR2F6
AAV-GFP
AAV-NR2F1
Avg.Exp
Umap2
Umap1
AAV-NR2F6
AAV-NR2F1
AAV-GFP
6177
4748
5403
0
2000
4000
6000
c
CST
Non-CST
CST
PVH
LH
RN
EW
PAG
LC
DP
Raphe
Inhib
HB-1-Daam2
HB-2-Pard3b
HB-4-Nox4
HB-5-Pappa
Cell Type / Cluster
Etv1
Crym
Satb2
Ccnd2
Spata16
Bcl11b
Cntnap3
Vgll3
Plscr4
Plagl1
Ros1
Rfx2
Slc25a48
Ttc6
Tnni3k
Emx2os
Cyyr1
Adamtsl5
Cartpt
Nostrin
Hs3st3b1
Crhbp
Ebf2
Slc6a2
Dbh
Th
Igsf5
Tshr
Lrrc36
Prdm6
Crh
Nlrp12
Tph2
Slc6a4
Trh
Maob
Slc22a3
Slc32a1
Daam2
Col5a2
Dmbx1
Crp
Nox4
Pappa
Markers (Grouped by Cell Type)
Average Expression
Percent Expressed
d
Satb2
Etv1
Crym
Ccnd2
Bcl11b
CST
Crh
Nlrp12
Cartpt
Nostrin
Cyyr1
Non- CST
Prdm6
Slc32a1
Col5a2
Tshr
Slc22a3
Percent Expressed
Average Expression

**Supplementary Figure S10: Single-nucleus RNA-seq clustering and cell-type annotation of AAV-transduced neurons.**

(a) Uniform Manifold Approximation and Projection (UMAP) embedding of snRNA-seq data revealing 13 distinct transcriptional clusters (0–12). The accompanying feature plots (right) display the average expression and distribution of viral transcripts within the diverse cellular populations across the three conditions: AAV-NR2F6, AAV-NR2F1, and AAV-GFP.
(b) Bar graph quantifying the total number of recovered nuclei for each respective AAV (AAV-NR2F6: n = 6177; AAV-NR2F1: n = 4748; AAV-GFP: n = 5403).
(c) Dot plot detailing the expression profiles of canonical marker genes utilized for cell-type annotation across all clusters. Evaluated lineages include Corticospinal Tract (CST), Paraventricular Hypothalamus (PVH), Lateral Hypothalamus (LH), Red Nucleus (RN), Edinger-Westphal nucleus (EW), Periaqueductal Gray (PAG), Locus Coeruleus (LC), Dorsal Pons (DP), Raphe nucleus, Inhibitory neurons (Inhib), and distinct Hindbrain populations (HB-1, HB-2, HB-4, HB-5). Dot size indicates the percentage of nuclei within a cluster expressing the specified gene, while color intensity represents the scaled average expression level. The robust expression of markers such as Satb2, Etv1, Crym, Ccnd2, and Bcl11b confirms that the major clusters represent CST neurons.
(d) Feature plots validating the spatial expression distribution of key CST marker genes (Satb2, Etv1, Crym, Ccnd2, Bcl11b) alongside representative non-CST markers across the UMAP space. The widespread and high-level expression of CST markers visually confirms that the vast majority of the captured nuclei belong to the CST lineage.

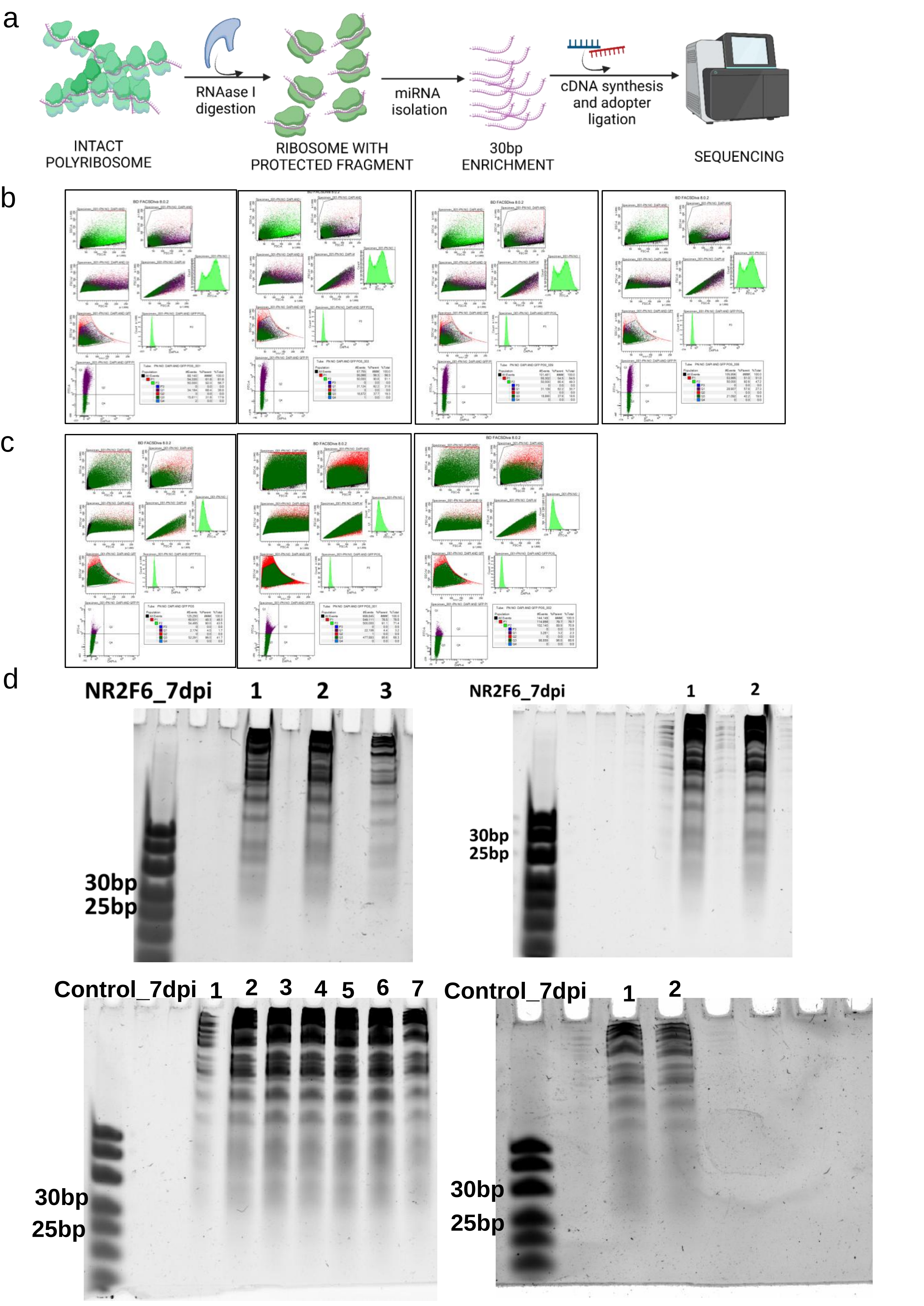
a
INTACT POLYRIBOSOME
RNAase I digestion
RIBOSOME WITH PROTECTED FRAGMENT
miRNA isolation
30bp ENRICHMENT
cDNA synthesis and adopter ligation
SEQUENCING
b
c
d
NR2F6_7dpi
1
2
3
30bp
25bp
NR2F6_7dpi
1
2
30bp
25bp
Control_7dpi
1
2
3
4
5
6
7
30bp
25bp
Control_7dpi
1
2
30bp
25bp

**NR2F1_7dpi**

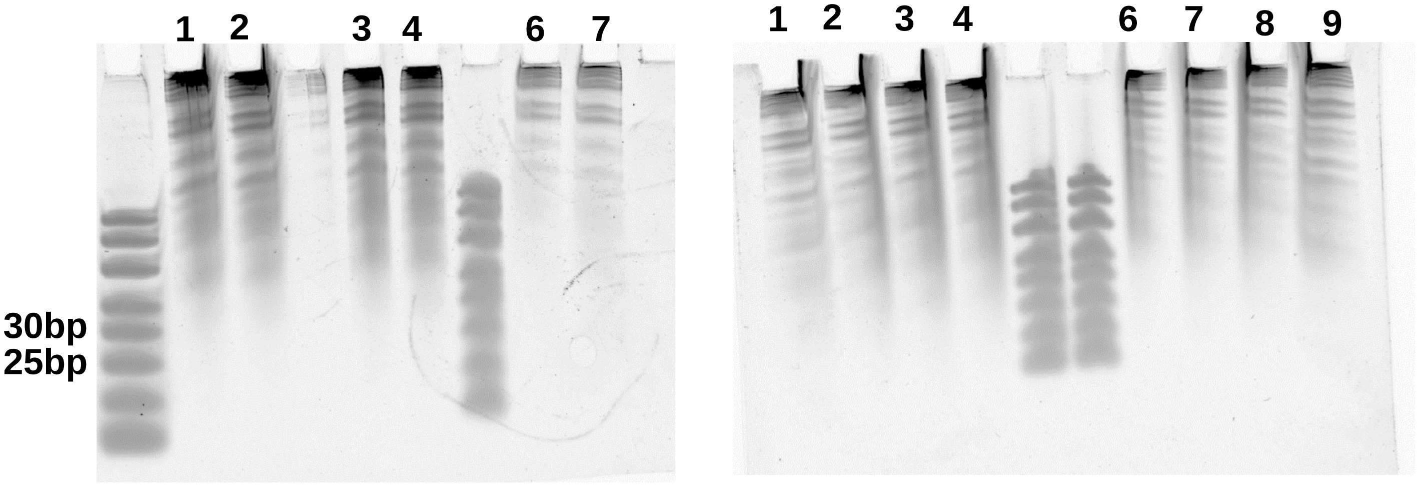


f

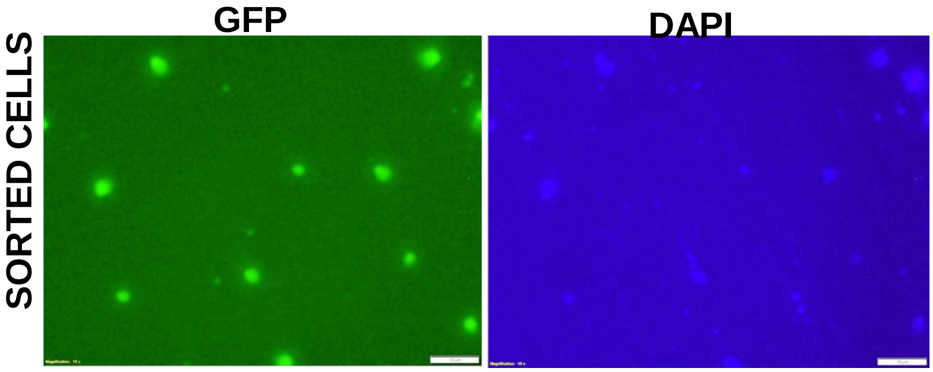


**Supplementary Figure S11: Validation of ribosome profiling workflow and 30 bp ribosome-protected fragment (RPF) enrichment from sorted GFP-positive cells.**
(a) Schematic illustration of the ribosome profiling workflow. Intact polyribosomes were digested with RNAse I to generate ribosome-protected fragments (~30 bp), followed by miRNA-sized fragment isolation, cDNA synthesis with adapter ligation, and sequencing. (b–c) Flow cytometry gating strategy for sorting GFP-positive neurons Control_7dp (b) and NR2F6_7dpi (c) samples. Sorted populations were used for downstream ribosome profiling. (d) TBS-UREA-PAGE images showing enrichment of 25–30 bp ribosome-protected fragments in NR2F6_7dpi ,Control_7dpi and NR2F1_7dpi ,across multiple replicates. Prominent enrichment of small RNA fragments confirms successful isolation of RPFs.(f) Fluorescent images of sorted cells showing GFP-positive neurons and DAPI-stained nuclei, confirming the presence of viable, labelled cells used for profiling Scale bar = 100µm.

A

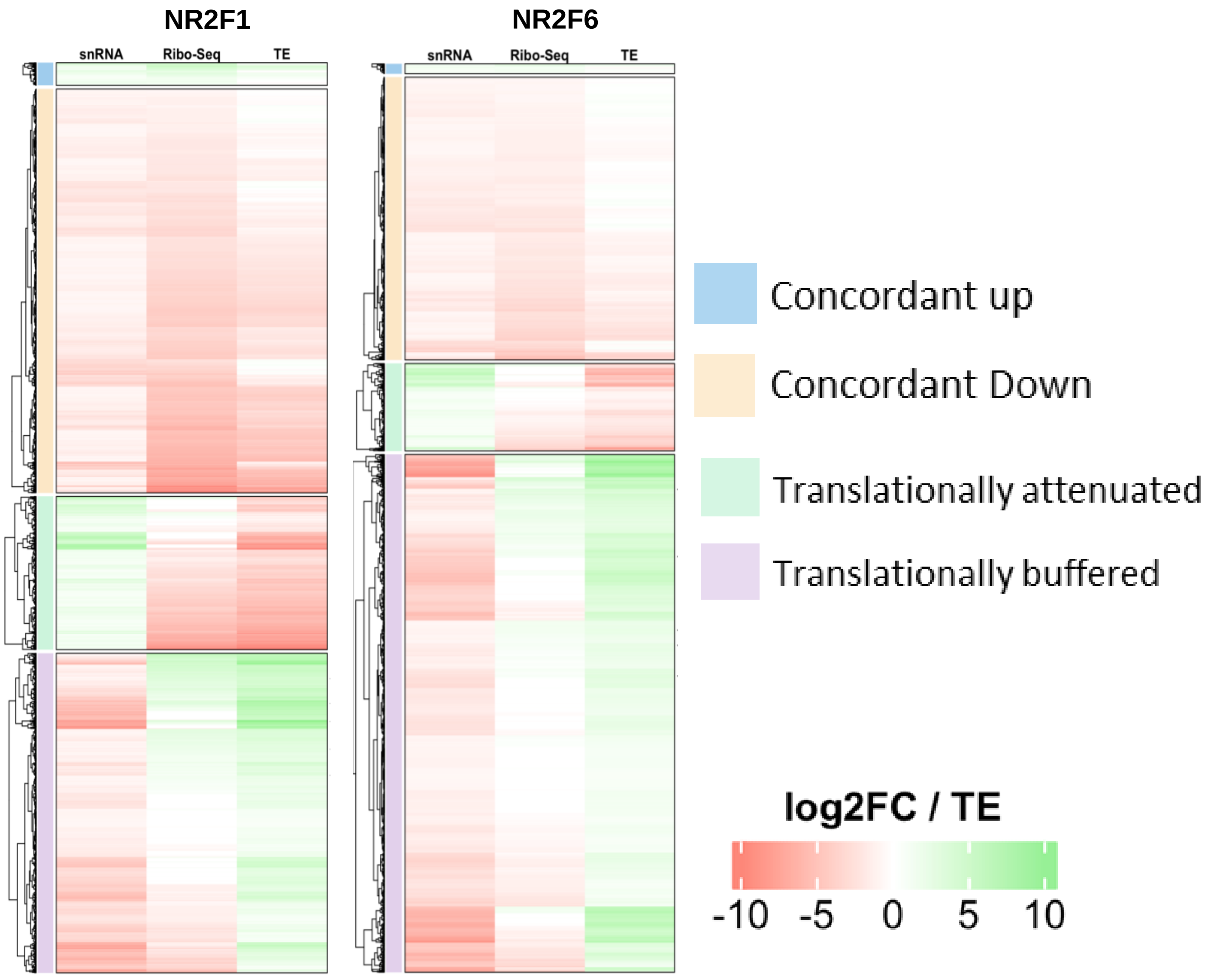


**Supplementary Figure S12: Translational buffering of gene expression following NR2F1 and NR2F6 gene treatment.**

(a) (Left) Heatmaps of snRNA-seq, Ribo-seq, and translational efficiency (TE) across gene categories for NR2F1 and NR2F6, confirming that buffered genes maintain positive TE despite reduced RNA. (Right) Representative IGV tracks for buffered genes (POLB, STAT1, OPHN1, ZEB2, NFE2L2 in NR2F1; PAK2, FOXO1, NOTCH2, EHHADH, PARP2 in NR2F6), showing maintained Ribo-seq signal alongside reduced snRNA signal at each locus.

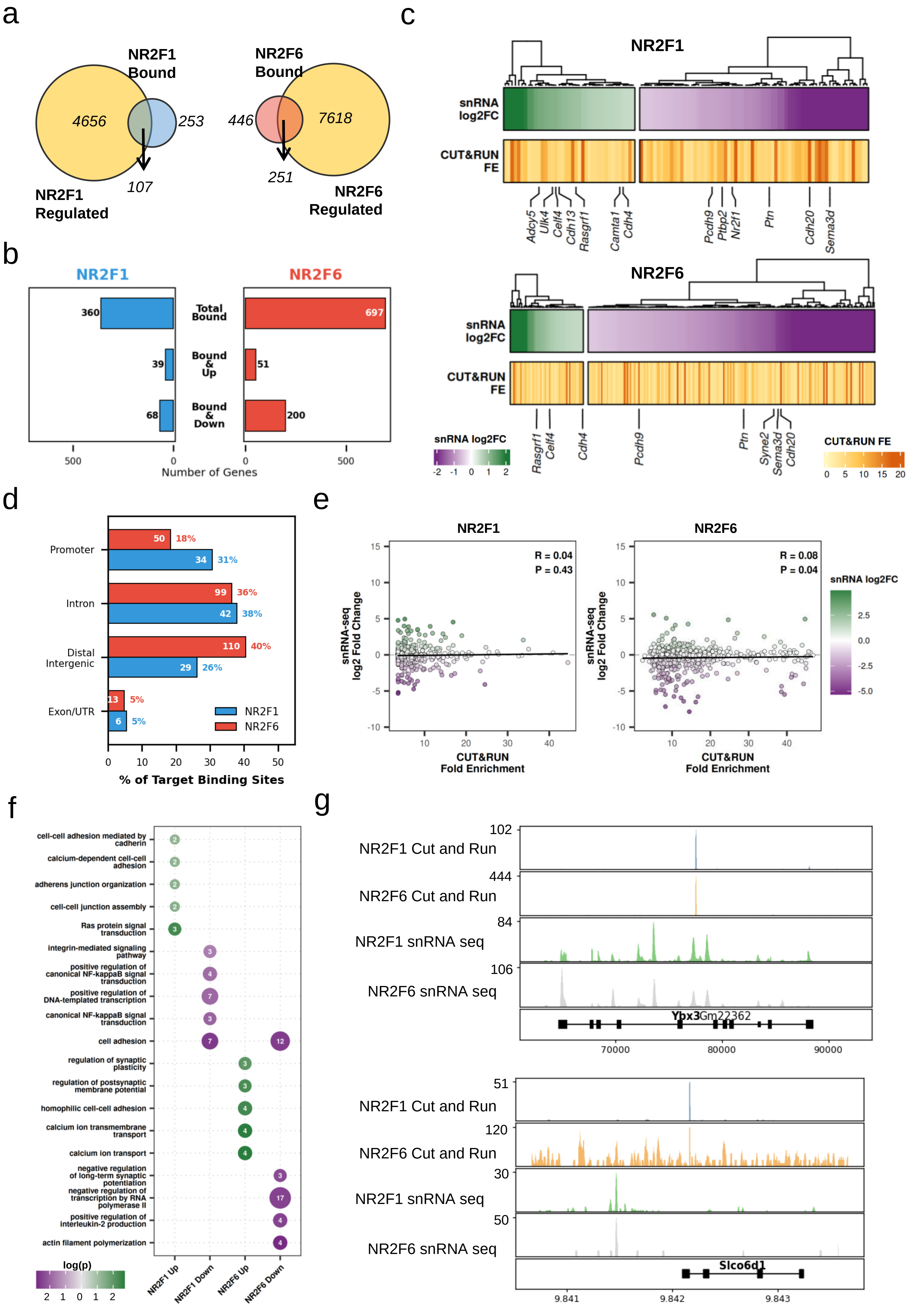

a
NR2F1 Bound
4656
253
107
NR2F1 Regulated
NR2F6 Bound
446
7618
251
NR2F6 Regulated
b
NR2F1
NR2F6
Total Bound
Bound & Up
Bound & Down
360
39
68
697
51
200
Number of Genes
c
NR2F1
NR2F6
snRNA log2FC
CUT&RUN FE
Adcy5
Ulk4
Celf4
Cdh13
Rasgrf1
Camta1
Cdh4
Pcdh9
Ptbp2
Nr2f1
Ptn
Cdh20
Sema3d
Syne2
CUT&RUN FE
d
Promoter
Intron
Distal Intergenic
Exon/UTR
50 18%
34 31%
99 36%
42 38%
110 40%
29 26%
13 5%
6 5%
NR2F1
NR2F6
% of Target Binding Sites
e
NR2F1
NR2F6
R = 0.04
P = 0.43
R = 0.08
P = 0.04
snRNA-seq log2 Fold Change
CUT&RUN Fold Enrichment
snRNA log2FC
f
cell-cell adhesion mediated by cadherin
calcium-dependent cell-cell adhesion
adherens junction organization
cell-cell junction assembly
Ras protein signal transduction
integrin-mediated signaling pathway
positive regulation of canonical NF-kappaB signal transduction
positive regulation of DNA-templated transcription
canonical NF-kappaB signal transduction
cell adhesion
regulation of synaptic plasticity
regulation of postsynaptic membrane potential
homophilic cell-cell adhesion
calcium ion transmembrane transport
calcium ion transport
negative regulation of long-term synaptic potentiation
negative regulation of transcription by RNA polymerase II
positive regulation of interleukin-2 production
actin filament polymerization
log(p)
NR2F1 Up
NR2F1 Down
NR2F6 Up
NR2F6 Down
g
NR2F1 Cut and Run
NR2F6 Cut and Run
NR2F1 snRNA seq
NR2F6 snRNA seq
Ybx3
Gm22362
Slco6d1

**Supplementary Figure S13: Integrative CUT&Run and snRNA-seq analysis reveals the direct transcriptional targets and genomic distribution of NR2F1 and NR2F6.**

(a) Venn diagrams illustrating the intersection between genes identified as directly bound by the respective transcription factors via CUT&Run (blue for NR2F1, salmon red for NR2F6) and the total pool of differentially regulated genes identified via snRNA-seq (yellow). The overlapping regions highlight 107 and 251 direct transcriptional targets for NR2F1 and NR2F6, respectively.
(b) Bidirectional bar plot quantifying the total number of CUT&Run bound genes, alongside the specific subsets of these bound targets that are significantly upregulated ("Bound & Up") or downregulated ("Bound & Down"). Transcriptional regulation was defined using a significance threshold of $p \leq 0.05$ and an absolute log2 fold change ($|log2FC| \geq 0.5$). NR2F1 data is depicted in blue; NR2F6 is depicted in red.
(c) Paired heatmaps detailing the expression and binding metrics for the overlapping target genes identified in (a) for NR2F1 (top) and NR2F6 (bottom). The upper tracks display the snRNA-seq log2FC, utilizing a diverging color scale (green for upregulation, purple for downregulation). The aligned lower tracks display the corresponding CUT&Run fold enrichment (FE) scores derived from MACS2 peak calling, represented by a yellow-to-orange continuous color gradient.
(d) Bar chart depicting the genomic distribution of NR2F1 (blue) and NR2F6 (red) binding sites. Annotations are categorized into Promoters, Introns, Distal Intergenic regions, and Exons/UTRs, expressed as a percentage of the total annotated target binding sites for each factor.
(e) Scatter plots assessing the global correlation between CUT&Run binding affinity (x-axis: Fold Enrichment) and transcriptional output (y-axis: snRNA-seq log2FC) for NR2F1 and NR2F6 targets. Data points are color-coded based strictly on their snRNA-seq expression profile (green: upregulated; purple: downregulated; grey: not significant). Pearson correlation coefficients (R) and associated p-values are denoted for each plot.
(f) Dot plot visualizing Gene Ontology (GO) biological process enrichment analysis for the directly bound and regulated targets. The plot displays the top 5 significantly enriched pathways for each of the four target categories (NR2F1 Up, NR2F1 Down, NR2F6 Up, NR2F6 Down). Dot color intensity reflects the statistical significance (log(p)).
(g) [Reserved for representative genomic browser tracks confirming specific binding peaks at key target loci].

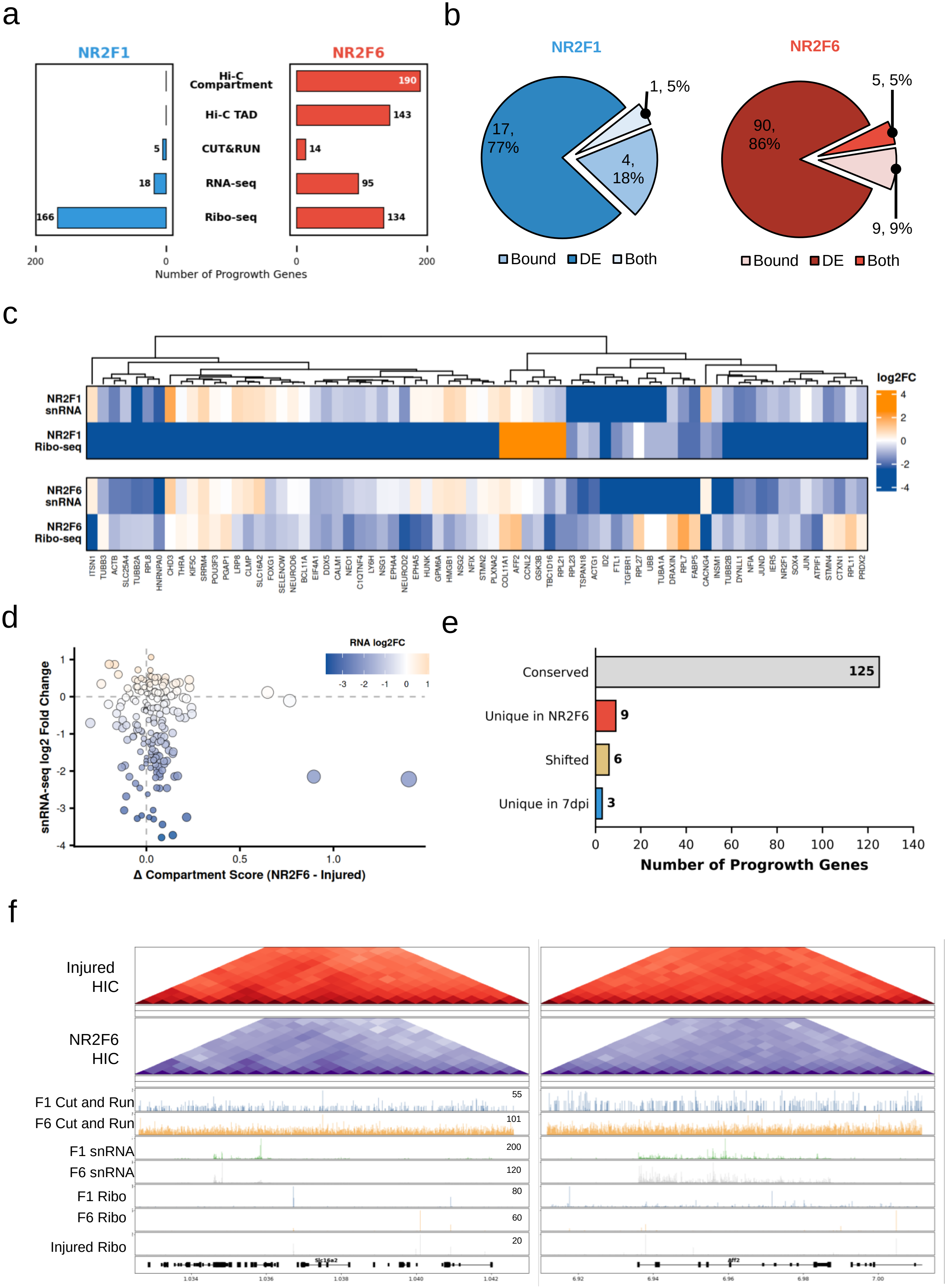
a
NR2F1
NR2F6
Hi-C Compartment
Hi-C TAD
CUT&RUN
RNA-seq
Ribo-seq
190
143
5
14
18
95
166
134
200
0
0
200
Number of Progrowth Genes
b
NR2F1
NR2F6
1, 5%
17, 77%
4, 18%
5, 5%
90, 86%
9, 9%
Bound
DE
Both
Bound
DE
Both
c
NR2F1 snRNA
NR2F1 Ribo-seq
NR2F6 snRNA
NR2F6 Ribo-seq
log2FC
4
2
0
-2
-4
ITSN1 TUBB3 ACTB SLC25A4 TUBB2A RPL8 HNRNPA0 CHD3 THRA KIF5C SRRM4 POU3F3 PGAP1 LRP8 CLMP SLC16A2 FOXG1 SELENOW NEUROD6 BCL11A EIF4A1 DDX5 CALM1 NEO1 C1QTNF4 LY6H NSG1 EPHA4 NEUROD2 EPHA5 HUNK GPM6A HMGB1 NSG2 NFIX STMN2 PLXNA2 COL11A1 AFF2 CCNL2 GSK3B TBC1D16 RPL21 RPL23 TSPAN18 ACTG1 ID2 FTL1 TGFBR1 RPL27 UBB TUBA1A DRAXIN RPL7 FABP5 CACNG4 INSM1 TUBB2B DYNLL1 NFIA JUND IER5 NR2F1 SOX4 JUN ATPIF1 STMN4 CTXN1 RPL11 PRDX2
d
RNA log2FC
-3 -2 -1 0 1
snRNA-seq log2 Fold Change
1
0
-1
-2
-3
-4
0.0
0.5
1.0
Δ Compartment Score (NR2F6 - Injured)
e
Conserved
125
Unique in NR2F6
9
Shifted
6
Unique in 7dpi
3
0 20 40 60 80 100 120 140
Number of Progrowth Genes
f
Injured HIC
NR2F6 HIC
F1 Cut and Run
F6 Cut and Run
F1 snRNA
F6 snRNA
F1 Ribo
F6 Ribo
Injured Ribo
55
101
200
120
80
60
20
Slc16a2
1.034 1.036 1.038 1.040 1.042
Aff2
6.92 6.94 6.96 6.98 7.00

**Supplementary Figure S14: Integrated multi-omics analysis of progrowth gene regulation by NR2F1 and NR2F6.**

(a) Bidirectional bar plot quantifying the number of progrowth genes exhibiting differential changes across varied multiomics data. The plot compares the regulatory impact of NR2F1 (blue, left) and NR2F6 (red, right) at the levels of translation (Ribo-seq), transcription (snRNA-seq), transcription factor binding (CUT&RUN), and chromatin architectur[illegible] (Hi-C TADs and Hi-C Compartments).

(b) Pie charts detailing the overlap between transcription factor binding and differential expression for the targeted progrowth genes. The diagrams illustrate the proportion and absolute number of genes that are exclusively bound, exclusively differentially expressed (DE), or both bound and DE under NR2F1 (blue) and NR2F6 (red) conditions.

(c) Heatmap representing the expression profiles of the progrowth gene panel. Rows indicate the specific omics modality (snRNA-seq and Ribo-seq) for NR2F1 and NR2F6, while columns represent individual genes. Color intensit[illegible] corresponds to the log2 fold change (log2FC) in expression.

(d) Scatter plot correlating the change in A/B compartment scores (Δ Compartment Score, NR2F6 - Injured) with the corresponding snRNA-seq log2 fold change, highlighting the relationship between structural chromatin shifts and transcriptional output.

(e) Horizontal bar plot summarizing the compartmental dynamics of the progrowth genes, categorized by their topological status: Conserved, Unique in NR2F6, Shifted, or Unique in 7dpi.

(f) Representative genomic tracks illustrating binding and accessibility landscapes at key progrowth loci (work in progress).
