## Supplementary_Table for "Nuclear Receptor Transcription Factors promote axon regeneration in the Adult Corticospinal Tract": Supplementary tables legend.docx

1. **Supplementary Table S1:** GO enrichment of the 194 developmentally downregulated pro-growth genes
2. **Supplementary Table S2:** List of the 194 candidate pro-growth genes
3. **Supplementary Table S3:** NR2F1 and NR2F6 footprinting across the 194 pro-growth loci
4. **Supplementary Table S4:** In vitro neurite-growth quantification and statistics
5. **Supplementary Table S5:** Pyramidotomy CST sprouting quantification
6. **Supplementary Table S6:** Thoracic crush CST regeneration quantification
7. **Supplementary Table S7:** Behavioural recovery quantification
8. **Supplementary Table S8:** DLC-derived gait metrics and Statistical analysis
9. **Supplementary Table S9:** snRNA-seq cluster annotation and marker genes
10. **Supplementary Table S10:** snRNA-seq DEG results for NR2F6 versus GFP/injured control
11. **Supplementary Table S11:** snRNA-seq DEG results for NR2F1 versus GFP/injured control
12. **Supplementary Table S12:** Pseudobulk snRNA-seq differential-expression validation
13. **Supplementary Table S13:** Direct NR2F1 versus NR2F6 transcriptional comparison
14. **Supplementary Table S14:** NR2F6 corepressor-domain mutant neurite-growth statistics
15. **Supplementary Table S15:** NR2F6 Ribo-seq differential-translation results
16. **Supplementary Table S16:** NR2F1 Ribo-seq differential-translation results
17. **Supplementary Table S17:** Translational-efficiency and buffering analysis
18. **Supplementary Table S18:** NR2F6 and NR2F1 CUT&RUN peak annotation
19. **Supplementary Table S19:** Developmental target of NR2F6 and F1 **Supplementary Table S20:** Overlap between developmental NR2F6 and F1 targets and post-injury CUT&RUN peaks
20. **Supplementary Table S21:** NR2F6-bound and transcriptionally regulated genes
21. **Supplementary Table S22:** NR2F1-bound and transcriptionally regulated genes
22. **Supplementary Table S23:** Promoter occupancy versus gene-expression correlation data
23. **Supplementary Table S23:** Promotor vs gene expression correlation of Cut and Run and snRNA seq.
24. **Supplementary Table S24:** Pro-growth 194-gene multi-omic categorisation
25. **Supplementary Table S25:** NR2F6-specific and shifted TADs
26. **Supplementary Table S26:** Genes within NR2F6-specific or shifted TADs linked to expression/Ribo-seq changes
27. **Supplementary Table S27:** A/B compartment switching after NR2F6 overexpression
28. **Supplementary Table S28:** GO enrichment of genes in NR2F6-remodelled TADs and switched compartments
29. **Supplementary Table S29:** Translation-regulatory hub genes
30. **Supplementary Table S30:** Statistical-analysis summary for all quantified panels
31. **Supplementary Table S31:** AAV-titter information
32. **Supplementary Table S32:** Cloning primers used
33. **Supplementary Table S33:** snRNA overexpression of NR2F1 and NR2F6 file
